# Ninhydrin as a covalent warhead for chemical proteomic-enabled discovery and selective engagement of reactive arginines

**DOI:** 10.64898/2026.01.05.697388

**Authors:** Andrew K. Ecker, Andreas Langen, Chloe S. Fields, José L. Montaño, Minh Tran, Ian B. Seiple, Balyn W. Zaro

## Abstract

Covalent molecules have emerged as next-generation therapeutics and as powerful tools for perturbing fundamental biological processes. Chemical proteomic methods to screen for reactive proteinaceous amino acids have transformed small-molecule discovery pipelines, but their application remains mostly limited to sites where reactive cysteines and lysines are present. Here we report a ninhydrin-based warhead that selectively modifies arginine residues, thus expanding the repertoire of amino acids targetable by covalent molecules. Specifically, we developed alkyne-functionalized variants of ninhydrin to establish an arginine-specific chemical proteomics platform, enabling the classification of more than 6,800 unique reactive arginines. These studies uncovered potential modification sites on disease-relevant proteins, including reactive arginines within catalytic sites that are essential for function. By endowing a reversible small molecule inhibitor of cyclophilin A with a ninhydrin warhead, we achieved selective, covalent engagement, and attenuation of enzymatic activity, highlighting the potential for targeting arginines in future therapeutic development campaigns. These findings establish ninhydrin as a warhead for studying arginine reactivity and modulating protein function.

## INTRODUCTION

Covalent therapeutics have revolutionized drug discovery by enabling precise and irreversible engagement of protein targets. Cysteine has been the predominant residue leveraged for covalent modifications, with approved drugs such as ibrutinib (BTK Cys481), afatinib (EGFR Cys797), and adagrasib/sotorasib (KRAS G12C) exemplifying this strategy. Evaluating and exploiting proteome-wide cysteine reactivity has been transformed by the use of broad-spectrum electrophiles, such as iodoacetamide-alkyne, that enable both reactivity profiling and competitive screening to assess a molecule of interest’s selectivity^1,2^. More recently, lysine residues have also emerged as attractive covalent drug targets, aided by probes that form imines or related adducts^3–5^. Beyond warheads selectively targeting a specific amino acid residue, such as cysteine and lysine, activity-based protein profiling (ABPP) strategies have begun to extend toward other nucleophilic side chains, although these remain less comprehensively mapped. SuFEx-based probes also enable the profiling of lysines and less conventional nucleophilic residues across the proteome, including histidine and tyrosine^6–9^.

Arginine plays indispensable roles in enzymatic catalysis, molecular recognition, and nucleic acid interactions. The positively charged guanidinium side-chain engages in a diverse array of noncovalent interactions, including salt-bridging and cation-π contacts, enabling arginine to stabilize transition states, mediate protein–protein interaction, and recognize aromatic functional groups within biomolecular targets^10^. However, its high pKa, resonance-stabilized charge distribution, and limited nucleophilicity complicate the development of covalent modification strategies. The reaction between vicinal dicarbonyl compounds with amidine and guanidine moieties has long been investigated from initial studies by Diels and Schleich^11^ to the Maillard reaction of sugar fragments with arginine^12^, yet the direct application of these reactions in biological systems has been quite limited. α,β-diketoamides^13^ and phenylglyoxal derivatives^14–17^ have been explored. Phenylglyoxals were found to have significant cross reactivity with lysines when used as a probe for activity-based protein profiling studies^17^. Chemoproteomic workflows using α,β-diketoamides as warheads have not yet been reported. The development of a selective and robust warhead for covalent pan-arginine modification would provide a powerful tool for studying arginine reactivity and function in complex biological contexts and facilitate the design of new therapeutic strategies.

Ninhydrin, an indanetrione, has long been utilized for detecting amino acids through its chromogenic reaction with primary amines at high temperature. Despite its established role in analytical chemistry, its potential as a selective covalent modifier of arginine has remained underexplored. Given its electrophilic properties and ability to rapidly form stabilized cyclic adducts with guanidine^18^, we hypothesized that ninhydrin could be repurposed for targeted covalent modification of arginine in biological systems. Furthermore, we postulated that if ninhydrin were reactive enough across proteinaceous arginine residues, further functionalization with a bioorthogonal handle, such as an alkyne, would allow for the development of a pan-arginine chemical proteomics platform akin to the iodoacetamide-alkyne probe and the isoTOPP-ABPP workflow, as developed by Cravatt and co-workers for cysteine^1,19^.

## RESULTS

### In vitro characterization of arginine engagement of ninhydrin

Many vicinal dicarbonyl compounds in aqueous environments exist predominantly in their hydrate state, reducing their electrophilicity. The equilibrium between this hydrate state and the reactive dehydrated state governs their ability to engage in covalent modification^20^. To investigate this equilibrium for ninhydrin versus the known arginine-reactive but promiscuous warhead phenylglyoxal (PGO)^21^, we performed density functional theory (DFT) calculations (**Figure 1A**). These values predict the relative occupancy of each state, allowing us to assess their potential for selective arginine modification. Our calculations revealed that both probes predominantly exist in their hydrate state; however, ninhydrin has a calculated K_eq_ of 3.55 × 10⁻³, indicating a higher fraction of the molecule resides in the reactive dehydrated state. PGO has a significantly lower calculated K_eq_ of 1.33 × 10⁻⁶, suggesting a much lower occupancy in the reactive form (**Figure 1A**). This favorable equilibrium positioning suggests ninhydrin would have a strong potential for arginine modification.

**Figure 1.**
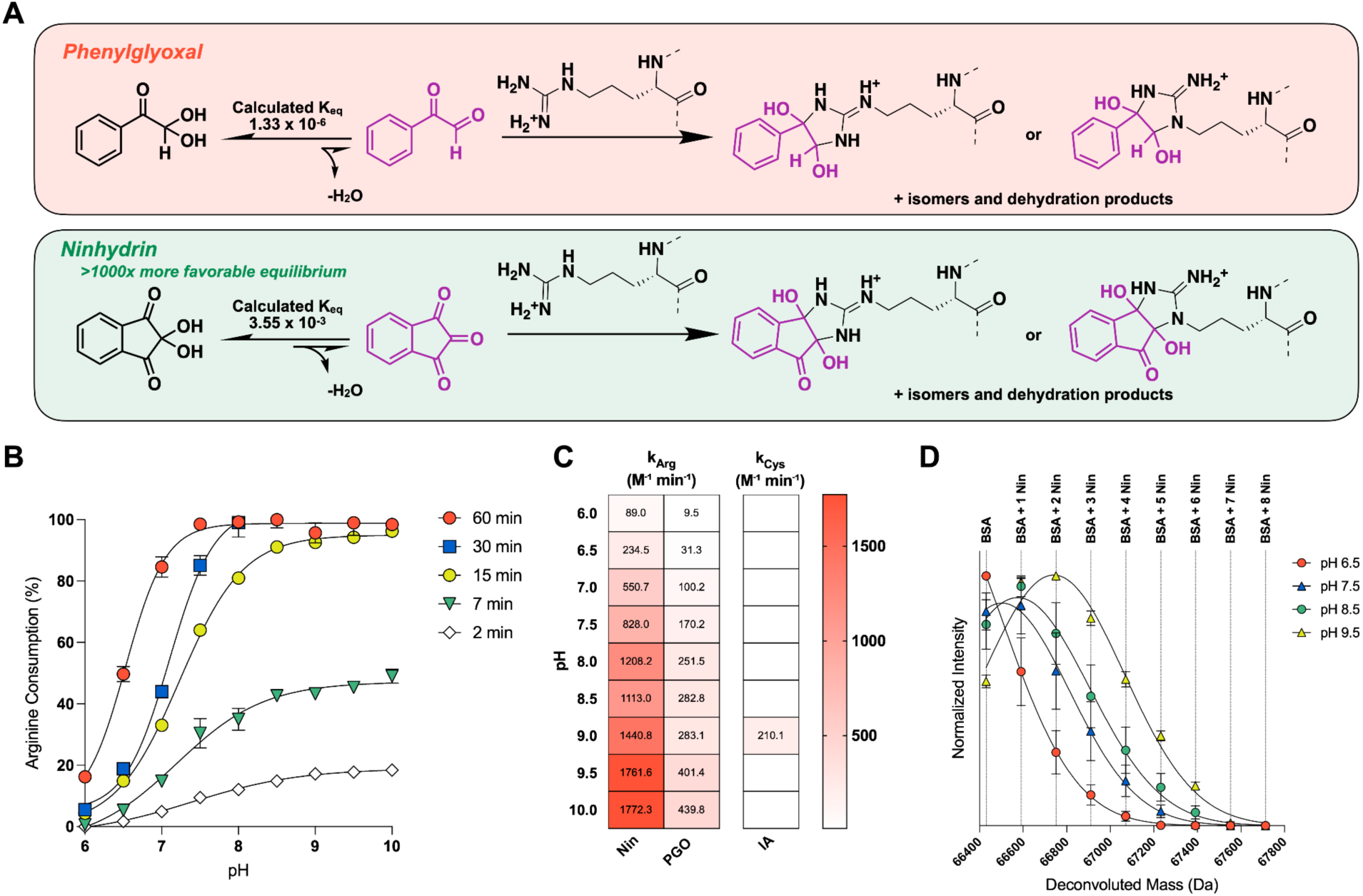
Profiling the reactivity of ninhydrin with arginine in vitro. **A.** Proposed reactions between phenylglyoxal and ninhydrin with arginine, highlighting differences in hydration equilibrium. Calculated equilibrium constants (K_eq_) were obtained by density functional theory (DFT) at the B3LYP/6-311G** level with implicit water solvation. **B.** Kinetic analysis of pH-and time-dependent consumption of Fmoc-Arg (500 µM) by ninhydrin (15 mM, 30x) in PBS at 20 °C. Arginine consumption was quantified by LC-MS over 2 to 60 minutes. Error bars represent SD. Representative HPLC traces in SI. n = 4 **C.** Heatmap depicting the second order reaction kinetics of ninhydrin or phenylglyoxal with Fmoc-arginine determined by UHPLC analysis compared to the reaction of iodoacetamide with cysteine determined by Ellman’s reagent. **D.** Intact protein mass spectrometry analysis of BSA (1.5 µM) treated with 150 µM ninhydrin at 37 °C for 30 minutes across pH 6.5 to 9.5. Reactions were quenched by dilution into 0.1% aqueous TFA and analyzed by LC–MS on a Agilent 6230 TOF System. Error bars represent SEM. n = 3

To validate these predictions, we characterized ninhydrin reactivity toward arginine using an HPLC-based kinetics assay monitoring the consumption of Fmoc-Arginine (**Figure 1B**). This approach provided direct evidence that the reaction proceeds at the terminal guanidine, while also enabling precise quantification via UV absorption. The 2nd order reaction kinetics were evaluated across a pH gradient ranging from 6.0 to 10.0, with time points recorded at 2, 7, 15, 30, and 60 minutes. At pH 7.5, complete consumption of free arginine was observed within 60 minutes (k_Arg, pH 7.5_ = 828.0 ± 86.8 M^−1^ min^−1^), whereas in more basic conditions reaction completion was achieved within 30 minutes (k_Arg, pH 9.5_ = 1761.6 ± 1.2 M^−1^ min^−1^) (**Figure 1B**). The relatively fast 2nd order reaction kinetics are comparable to that of iodoacetamide with cysteine (k_cys, pH 9.0_ = 210.1 ± 30.2 M^−1^ min^−1^) and further supported our hypothesis that ninhydrin may be a warhead amenable to engaging arginines in biological systems on a reasonable timescale^22^. When we compared these rates to the reaction of PGO towards arginine we found that on average a 6-fold decrease in reactivity(k_Arg, pH 7.5_ = 170.2 ± 0.2 M^−1^ min^−1^) with PGO never fully consuming the Fmoc-Arg at an hour even at more basic pHs (**Figures 1C and 3C, Extended Data Figure 3A**). A similar study was performed with ninhydrin towards Fmoc-lysine and glutathione^23^, which did not result in appreciable engagement (**Extended Data Figure 1**). Conversely, PGO showed engagement of glutathione, consistent with prior reports of PGOs reactivity towards other nucleophilic sidechains^17^ (**Extended Data Figure 1B**).

To probe the extent of ninhydrin labeling and its potential to react with proteinaceous arginines, we examined its modification stoichiometry on a model protein, bovine serum albumin (BSA), which contains 23 arginine residues. Following 30 minutes of ninhydrin treatment, intact protein mass spectrometry revealed up to 6 labeling events under physiologic conditions (**Figure 1D** and **Supplemental Figure 1A**), with increased reactivity in a pH-dependent manner, as we observed in our free-arginine kinetics assay. The mass shift for each labeling event indicated addition of ninhydrin, consistent with engagement of ninhydrin with a nucleophilic sidechain forming dehydration products. Similar studies were also conducted for lysozyme, revealing a comparable pH-dependent reactivity profile modifying up to 11 residues on this protein with 11 arginines^24^ (**Extended Data Figure 1C** and **Supplemental Figure 1B**). While it was possible that engagement of other sidechains could result in a similar mass shift, it was unlikely that all labeling events could be attributed to non-arginine sidechains due to the reversibility of those reactions compared to arginine, especially considering that free lysine and cysteine are not detectably engaged by ninhydrin. Therefore, though possible that some of this labeling could be attributed to lysine or cysteine engagement, we postulated that it was unlikely that all of the labeling could be explained through this mechanism.

### In vitro characterization of arginine engagement of ninhydrin

Many vicinal dicarbonyl compounds in aqueous environments exist predominantly in their hydrate state, reducing their electrophilicity. The equilibrium between this hydrate state and the reactive dehydrated state governs their ability to engage in covalent modification^20^. To investigate this equilibrium for ninhydrin versus the known arginine-reactive but promiscuous warhead phenylglyoxal (PGO)^21^, we performed density functional theory (DFT) calculations (**Figure 1A**). These values predict the relative occupancy of each state, allowing us to assess their potential for selective arginine modification. Our calculations revealed that both probes predominantly exist in their hydrate state; however, ninhydrin has a calculated K_eq_ of 3.55 × 10⁻³, indicating a higher fraction of the molecule resides in the reactive dehydrated state. PGO has a significantly lower calculated K_eq_ of 1.33 × 10⁻⁶, suggesting a much lower occupancy in the reactive form (**Figure 1A**). This favorable equilibrium positioning suggests ninhydrin would have a strong potential for arginine modification.

To validate these predictions, we characterized ninhydrin reactivity toward arginine using an HPLC-based kinetics assay monitoring the consumption of Fmoc-Arginine (**Figure 1B**). This approach provided direct evidence that the reaction proceeds at the terminal guanidine, while also enabling precise quantification via UV absorption. The 2nd order reaction kinetics were evaluated across a pH gradient ranging from 6.0 to 10.0, with time points recorded at 2, 7, 15, 30, and 60 minutes. At pH 7.5, complete consumption of free arginine was observed within 60 minutes (k_Arg, pH 7.5_ = 828.0 ± 86.8 M^−1^ min^−1^), whereas in more basic conditions reaction completion was achieved within 30 minutes (k_Arg, pH 9.5_ = 1761.6 ± 1.2 M^−1^ min^−1^) (**Figure 1B**). The relatively fast 2nd order reaction kinetics are comparable to that of iodoacetamide with cysteine (k_cys, pH 9.0_ = 210.1 ± 30.2 M^−1^ min^−1^) and further supported our hypothesis that ninhydrin may be a warhead amenable to engaging arginines in biological systems on a reasonable timescale^22^. When we compared these rates to the reaction of PGO towards arginine we found that on average a 6-fold decrease in reactivity(k_Arg, pH 7.5_ = 170.2 ± 0.2 M^−1^ min^−1^) with PGO never fully consuming the Fmoc-Arg at an hour even at more basic pHs (**Figures 1C and 3C, Extended Data Figure 3A**). A similar study was performed with ninhydrin towards Fmoc-lysine and glutathione^23^, which did not result in appreciable engagement (**Extended Data Figure 1**). Conversely, PGO showed engagement of glutathione, consistent with prior reports of PGOs reactivity towards other nucleophilic sidechains^17^ (**Extended Data Figure 1B**).

To probe the extent of ninhydrin labeling and its potential to react with proteinaceous arginines, we examined its modification stoichiometry on a model protein, bovine serum albumin (BSA), which contains 23 arginine residues. Following 30 minutes of ninhydrin treatment, intact protein mass spectrometry revealed up to 6 labeling events under physiologic conditions (**Figure 1D** and **Supplemental Figure 1A**), with increased reactivity in a pH-dependent manner, as we observed in our free-arginine kinetics assay. The mass shift for each labeling event indicated addition of ninhydrin, consistent with engagement of ninhydrin with a nucleophilic sidechain forming dehydration products. Similar studies were also conducted for lysozyme, revealing a comparable pH-dependent reactivity profile modifying up to 11 residues on this protein with 11 arginines^24^ (**Extended Data Figure 1C** and **Supplemental Figure 1B**). While it was possible that engagement of other sidechains could result in a similar mass shift, it was unlikely that all labeling events could be attributed to non-arginine sidechains due to the reversibility of those reactions compared to arginine, especially considering that free lysine and cysteine are not detectably engaged by ninhydrin. Therefore, though possible that some of this labeling could be attributed to lysine or cysteine engagement, we postulated that it was unlikely that all of the labeling could be explained through this mechanism.

### Ninhydrin probe development

Encouraged by Fmoc-arginine engagement and robust, rapid labeling of two purified proteins, we pursued the development of a ninhydrin chemical probe for testing in complex proteomes. We synthesized a ninhydrin-derived probe (**Nin-Alk**) and the reported arginine-reactive probe **PGO**^17^ (**Figure 2A**). We visualized their relative reactivity across the proteome using gel-based chemical probe techniques (**Figure 2B**, and **Supplemental Figure 2A**). Mino cell lysates were treated with the indicated probe for 1h. Following incubation, lysates were subjected to treatment with an azide-conjugated TAMRA fluorochrome under Cu(I)-catalyzed Azide-Alkyne Cycloaddition (CuAAC)^26^ conditions. Methanol was added, and precipitated proteins were resuspended and separated by SDS-PAGE. In-gel fluorescence scanning revealed dose-dependent labeling for both probes across the proteome, with **Nin-Alk** exhibiting higher levels of labeling at all concentrations tested (**Figure 2B**). As a benchmark, we compared proteome-wide labeling by **Nin-Alk** and **PGO** to the pan-cysteine probe iodoacetamide-alkyne (**IA-Alk**)^1^. Both **Nin-Alk** and **PGO** exhibited reduced labeling compared to **IA-Alk** (**Figure 2B**). Curious as to if a regioisomer of **Nin-Alk** would have similar labeling, we also synthesized **4-Nin-Alk** (**Figure 2A**), which exhibited similar labeling of cell lysates compared to **Nin-Alk** (**Extended Data Figure 2A**).

**Figure 2.**
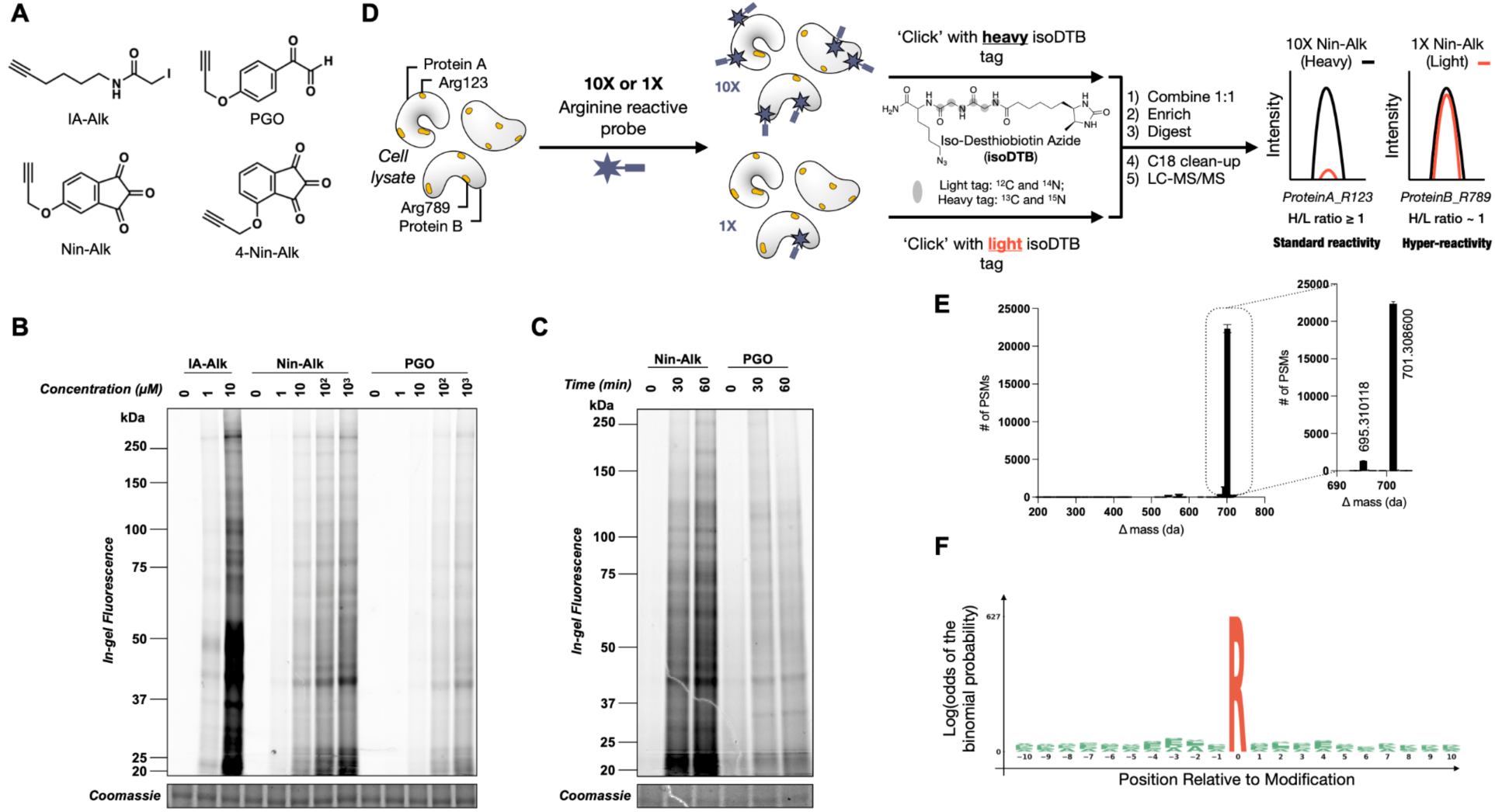
Comparative profiling and biochemical characterization of arginine-reactive electrophiles. **A.** Chemical structures of electrophilic probes used in this study, including **IA-Alk** (iodoacetamide alkyne, cysteine-reactive control), phenylglyoxal alkyne (**PGO**), and two alkyne-functionalized ninhydrin derivatives, **4-Nin-Alk** and **Nin-Alk**. **B.** Qualitative, gel-based assessment of proteome-wide reactivity of alkyne-bearing covalent probes in Mino cell lysate. Soluble proteome (1.5 µg µL⁻¹) was treated with each probe at the indicated concentration for 1 h, followed by CuAAC conjugation with TAMRA-N₃, separation by SDS–PAGE, and in-gel fluorescence scanning. **C.** Qualitative time-dependent reactivity of **Nin-Alk**, and **PGO** in live cells. Mino cells were treated with 100 µM of the indicated probe for 0, 30, or 60 min, followed by lysis, CuAAC conjugation with TAMRA-N₃, separation by SDS–PAGE, and in-gel fluorescence scanning. Uncropped gel images available in SI. **D.** Schematic of the Reactive Arginine Profiling (RAP) workflow to assess arginine reactivity. Cell lysates were treated with an arginine-reactive electrophile at high (1 mM) or low (100 µM) concentrations and subjected to copper-catalyzed azide-alkyne cycloaddition (“click” chemistry) with isotopically light or heavy isoDTB tags. Samples were combined 1:1, enriched, digested, and analyzed by LC-MS/MS. Peptides from hyper-reactive arginine residues show similar heavy/light (H/L) intensities, whereas less reactive sites are preferentially labeled at higher probe concentrations, yielding elevated H/L ratios. **E.** Open-mass search analysis of modified peptides reveals a predominant Δmass of 701.308600 and 695.310118 Da, corresponding to covalent modification of arginine by **Nin-Alk** and heavy or light isoDTB tag conjugation, respectively. Error bar represents SD **F.** pLogo motif analysis^25^ of the ±10-residue sequence window surrounding all modified arginines (foreground: 6,913 modified sites; background: 773,893 human arginines.

We next investigated whether **Nin-Alk** was capable of labeling live cells in a time-dependent manner. Mino cells were treated for 0, 30, or 60 min with the indicated probe (100 µM). Labeled cells were pelleted, lysed, and subjected to conjugation with TAMRA-azide as described above. In-gel fluorescence scanning of SDS-PAGE separated proteins revealed improved time-dependent labeling with **Nin-Alk** compared to **PGO** (**Figure 2C** and **Supplemental Figure 2B**). Notably, **4-Nin-Alk** also exhibited markedly reduced labeling in live cells compared to **Nin-Alk**, affirming our focus on **Nin-Alk** as a potential probe for characterizing arginine reactivity.

To evaluate **Nin-Alk**’s potential as a pan-arginine chemical probe, we developed a workflow utilizing the isotopically distinct azide-bearing desthiobiotin (isoDTB) tags developed by Hacker and co-workers^27^ (**Figure 2D**). Cell lysates were treated with **Nin-Alk** at either 100 μM or 1 mM for 1h, and the resulting alkyne-labeled arginines were conjugated to light or heavy IsoDTB tags via copper(I)-catalyzed azide-alkyne cycloaddition (CuAAC). Labeled proteome pairs were combined and processed by single-pot, solid-phase sample preparation (SP3), yielding enriched, desalted peptides for subsequent mass spectrometry analysis^28^. We call this pipeline Reactive Arginine Profiling (RAP).

We first performed an open search to validate detection of intact probe modification. Open searches characterize the frequency of mass modifications to peptides, which allows for the identification of large mass differences between unmodified peptide sequences and experimentally observed precursors^29^. For our **Nin-Alk**-treated lysates, we observed one major mass modification, +701.308600 da, which corresponds to the cyclic mono-dehydration adduct of arginine and **Nin-Alk** clicked to the heavy isoDTB tag. A less abundant mass corresponding to the cyclic mono-dehydration adduct of arginine and **Nin-Alk** clicked to the light isoDTB tag was also detected (+695.310118 da, **Figure 2E** and **Table 1**), which we expected given the 10-fold lower concentration of **Nin-Alk** used in the light channel. pLOGO analysis of peptides with a modified arginine found no significant 2D sequence determinants of labeling with some minor enrichment for acidic residues 3-4 residues away (**Figure 2F**)^25^. Finally, in further support of arginine selectivity, our RAP samples were prepared in the absence of a reducing agent, which is often required for the trapping of lysine modifications.

We next compared the performance of **Nin-Alk** to the **PGO** probe in a comparative RAP experiment in which paired lysates were each treated with either probe (1mM; 1h) and conjugated to either a heavy or light isoDTB tag. UpSet plot analysis revealed that the majority of **PGO**-engaged peptides were also engaged by **Nin-Alk** (n = 2862 of 2871 arginines are shared) while remarkably few arginines are uniquely engaged with **PGO** (n = 9) (**Figure 3A**, **Table 2**). This high overlap suggests that most **PGO**-reactive arginines are also accessible by **Nin-Alk**. Conversely, an additional 713 arginines are uniquely engaged by **Nin-Alk**, suggestive that the probe engages a broader subset of arginines in the proteome. While less intrinsically reactive, **PGO** remains an important electrophile within the arginine-targeting toolkit, offering a more tempered reactivity profile that can facilitate more discriminating labeling and may prove advantageous in experimental contexts demanding precise or attenuated engagement of reactive residues.

**Figure 3.**
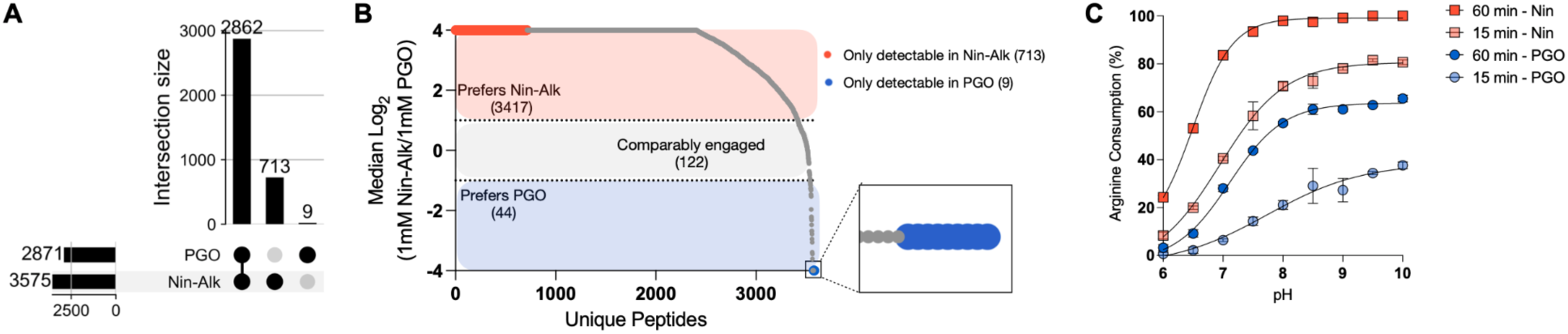
Comparative profiling of phenylglyoxal (PGO) and Nin-Alk demonstrates increased saturation and promiscuity for Nin-Alk. **A.** UpSet plot demonstrating high overlap between **PGO**-engaged arginines and **Nin-Alk**-engaged arginines at 1 mM (1 h), with **Nin-Alk** engaging more targets. **B.** Waterfall plot showing median log_2_ ratios of 1 mM **Nin-Alk** to 1 mM **PGO** isoDTB enrichment (n = 6). Peptides with log_2_(**Nin-Alk**/**PGO**) ≥ 1 were classified as preferring **Nin-Alk**, whereas those with log_2_(**Nin-Alk**/**PGO**) ≤ −1 preferred **PGO**. Because **PGO** generates a distinct oxidation side product, quantification was performed by merging closed searches corresponding to oxidized and unoxidized species; intensities for both light channels (± ox) were summed for each peptide, and the heavy channel was averaged across searches to prevent double-counting. **C.** Reactivity analysis of pH- and time-dependent consumption of Fmoc-Arg (500 µM) by ninhydrin (Nin, 15 mM, 30x) or phenylglyoxal (PGO, 15 mM, 30x) in PBS at 20 °C. Arginine consumption was quantified by UHPLC analysis of reactions quenched at 15 or 60 minutes. Error bars represent SD n = 4

Most arginines engaged by both probes were preferentially engaged by **Nin-Alk** (2704 arginines), whereas a significantly smaller fraction of sites were favored or exclusively engaged by **PGO** (n = 44) (**Figure 3B**). This pattern suggests that **PGO** frequently does not reach equivalent occupancy of shared arginines under identical conditions. Solution-phase Fmoc-Arginine reactivity mirrored these proteomic trends, with ninhydrin showing faster and more extensive arginine consumption than **PGO** across all tested pH values and time points (**Fig 3C** and **Extended Data Figure 3A**). Together, these findings suggest that **Nin-Alk** achieves a broader and more uniform engagement of reactive arginines and is therefore well suited for proteome-wide screening applications analogous to iodoacetamide-alkyne in cysteine profiling. It is also worth noting that the **PGO** adduct has been reported to oxidize during sample preparation and data acquisition^17^. We also observed this limitation (**Table 3**), which further complicates quantitative analysis. For our comparative studies we were careful to perform searches with both the non-oxidized modification and oxidized modification (**Extended Data Figures 3B** and **3C, Tables 2, 4** and **5**). These data revealed comparable reactivity profiles and combining both searches into our quantitative pipeline did not alter our results (**Figure 3B**, **Extended Data Figures 3B** and **3C** and **Tables 2, 4** and **5**). In contrast, **Nin-Alk** yields a single dominant mass adduct and clean isotopic labeling profiles, affirming its suitability as a general arginine-targeting chemical probe.

### Chemical proteomic profiling of arginine labeling

Having validated the arginine-targeted pan-reactivity of **Nin-Alk**, we next leveraged the data to identify reactive arginines embedded in proteins. Raw data collected from the RAP workflow comparing 1 mM and 100 μM **Nin-Alk** labeling events (which has been standard for arginine-targeted datasets, given their increased frequency in the proteome compared to cysteine) were subjected to a modified isoDTB closed-search workflow (**Figure 2D**)^27^. From Mino cell lysates, we captured 10,642 modified peptide ions mapping to 6,888 unique arginines across 2,076 distinct proteins (**Figure 4A** and **Table 6**). Despite a recent observation from the Hacker group^17^ that modified arginines are very rarely (0.2 %) recognized by trypsin as a proteolytic cleavage site, we found that a greater subset (6.74 %) of our modifications were still being recognized by trypsin as a cleavage site. We performed similar RAP studies using **Nin-Alk** in lysates and live cells at a lower concentration range, 100 μM and 10 μM, as we felt these concentrations were more appropriate for live cell studies (**Extended Data Figure 4** and **Extended Tables**). Strikingly, our lysate and in cell studies at these lower concentrations resulted in less than 100 quantified arginines for each condition. While live cell data contained intracellular proteins of interest such as the Akt modulator TCL1A, mitochondrial GTPase EFTU, and the endoplasmic reticulum disulfide isomerase PDIA3, suggestive of cell uptake of the **Nin-Alk** probe, the low coverage nature of our lower concentration pairing deprioritized these experimental conditions (**Extended Data Figure 4A**). Therefore, we moved forward analyzing our cell lysate dataset comparing 1 mM and 100 μM **Nin-Alk** engagement of nearly 7,000 arginines.

**Figure 4.**
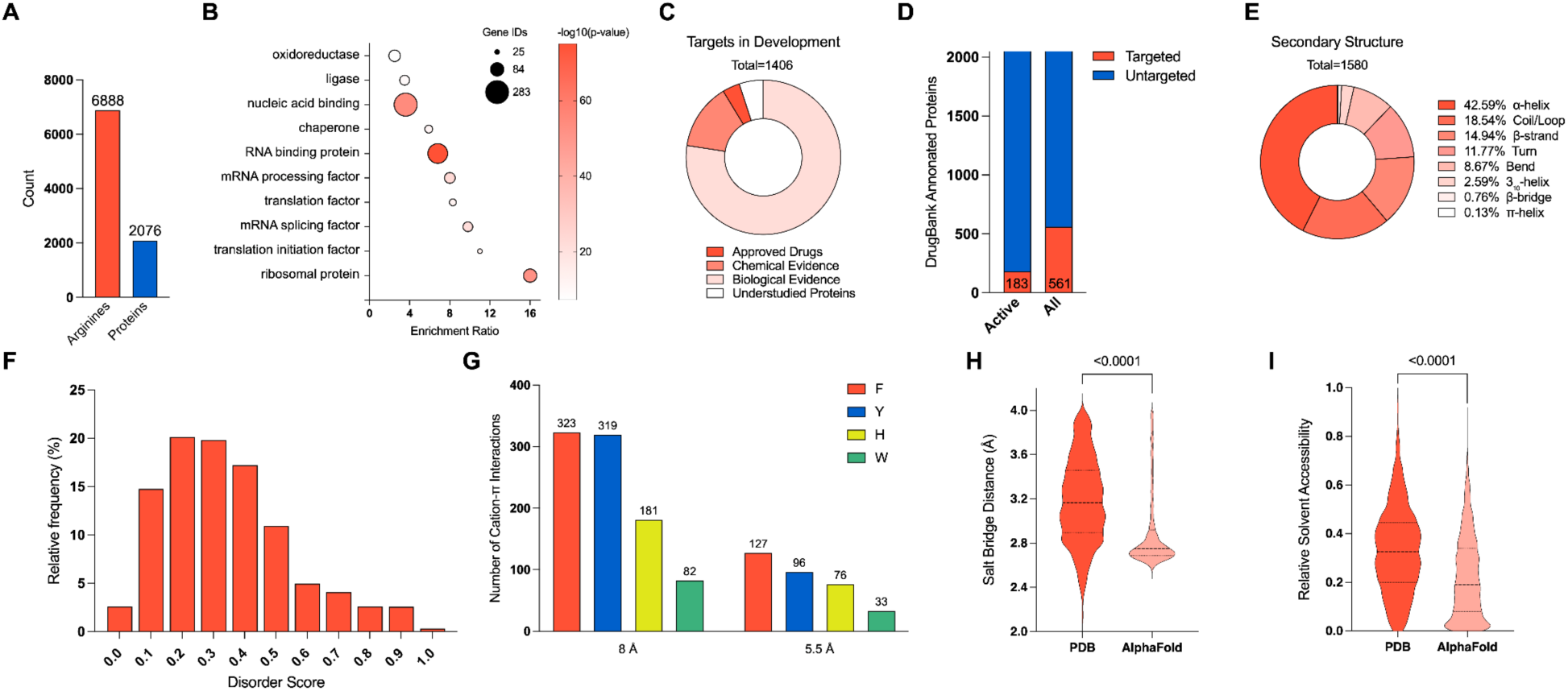
Structural and functional context of arginines identified by RAP in Mino cell lysates. **A.** Total number of unique arginines and corresponding proteins identified using the RAP workflow (see **Figure 2D** and **Table 6**). **B.** Bubble plot of the top 10 most significantly enriched PANTHER protein classes. Statistical significance was calculated using Fisher’s Exact test and adjusted using the Benjamini–Hochberg procedure to control the false discovery rate at α = 0.05; only terms with adjusted p-value < 10⁻⁹ are shown. Bubble size reflects the number of proteins in each class. **C.** Target development level of proteins containing reactive arginines, based on Target Central Resource Database classification, where Tclin = Approved Drug, Tchem = Chemical Evidence, Tbio = Biological Evidence, Tdark = Understudied Proteins**. D.** Number of proteins detected in our RAP dataset that are annotated as targeted in the DrugBank. **E.** Distribution of secondary structure assignments for reactive arginines, as annotated using DSSP on PDB structures(**Table 11**). **F.** Histogram of predicted intrinsic disorder scores at modified arginine sites, calculated using IUPred2A. Scores > 0.5 indicate disordered regions. (n = 3630) **G.** Quantification of cation–π interactions between reactive arginines and aromatic residues in PDB deposited structures. Interaction counts are shown at distance thresholds of 8.0 or 5.5 Å, measured from the guanidinium CZ atom to the centroid of the aromatic ring. Geometric constraints were applied for inclusion of parallel interactions (face-on) when the angle between the CZ-to-centroid vector and the normal vector of the aromatic plane was ≤ 30°, and T-shaped interactions (edge-on) when deviation from 90° was ≤ 20°(**Table 7**). **H.** Violin plot showing the distribution of salt bridge distances between guanidinium nitrogen atoms of reactive arginines and carboxylate oxygens of acidic residues (Asp/Glu) with discrepancies between PDB (n = 1594) and AlphaFold (n = 1445) models (**Tables 7 and 8**). Welch’s t-test (two-tailed, parametric) **I.** Predicted relative solvent accessibility (RSA) of reactive arginines, computed using the Shrake–Rupley algorithm on PDB (n = 1649) and predicted AlphaFold (n = 4309) structures, indicating an over-representation of arginine surface burial in AlphaFold models compared to PDB models(**Tables 9 and 10**).Welch’s t-test (two-tailed, parametric).Inspired by the unique roles arginines play in protein structure and scaffolding, we sought to characterize the broader structural and functional context of **Nin-Alk** enriched residues. Through secondary-structure mapping across the PDB, we found reactive arginines are enriched in helices and more flexible loops compared to β-sheets (**Figure 4E** and **Table 11**). To gauge local flexibility, we used IUPred2A, an energy-based predictor of intrinsic disorder, to compute per-residue scores from 0 (ordered) to 1 (disordered). Reactive arginines peaked in the moderate-disorder regime (0.3 - 0.4) yet ∼ 40% reside in well-ordered contexts (scores < 0.2), demonstrating that both structural support and some degree of flexibility facilitate efficient covalent engagement (**Figure 4F**).

Statistical overrepresentation analysis of proteins containing reactive arginines revealed enrichment for protein classes associated with protein translation, mRNA processing, and nucleic acid binding, among others (**Figure 4B**)^30,31^. We also cross-referenced proteins harboring reactive arginines with Pharos^32^, a database developed by the NIH to categorize a protein’s status as a drug target based upon available chemical and biological evidence. This analysis revealed that the majority of proteins harboring reactive arginines are potential therapeutic targets supported by strong biological evidence but currently lack chemical tools for modulation (**Figure 4C**). Complementary annotations from DrugBank^33^ further suggest an opportunity to expand covalent small-molecule engagement to a broader set of potentially druggable proteins through reactive arginine engagement (**Figure 4D**).

Beyond these structural contexts, we turned to cation-π contacts with aromatic side chains, quantifying interactions across all available PDB models for each protein (**Figure 4G and Table 7**). We applied two distance thresholds, 8 Å, which is the commonly used cutoff, and a more stringent 5.5 Å cutoff for true energetic relevance, measured from the arginine CZ atom to the aromatic ring-centroid. To ensure only geometrically plausible face-on and edge-on interactions were included, we added angular filters: for parallel (face-on) interactions, the CZ-to-centroid vector was required to lie within 30° of the ring normal, while for T-shaped (edge-on) interactions, deviation from 90° had to be ≤ 20°. Using these criteria, we identified 905 total cation-π contacts at 8.0 Å, while only 332 satisfied the stricter 5.5 Å threshold (**Figure 4G and Table 7**). Phenylalanine and tyrosine accounted for the majority of these contacts, compared to histidine and tryptophan. At the more stringent distance threshold, the distribution of aromatic partners mirrors the natural abundance of aromatic residues in the human proteome (**Table 7**). Unlike recent efforts to selectively label tryptophans that showed enrichment for tryptophans engaged in cation-π interactions^34^, we do not see an enrichment of arginines engaged in cation-π interactions (**Extended Data Figure 5A and Table 7**). This indicates that arginine reactivity is not biased by a specific type of π-donor or the stabilizing effect of a cation–π interaction.

We next measured salt bridges as the minimum distance between any arginine guanidinium nitrogen and Asp/Glu carboxylate oxygens for both PDB entries and AlphaFold^35^ models. In AlphaFold structures, these distances cluster tightly around ∼2.6 to 2.8 Å (**Extended Data Figure 5B, 5C, and 5D, Table 8**), whereas PDB entries span a broader 2.5 to 4.0 Å range with a median closer to 3.2 Å (**Figure 4H, Extended Data Figure 5E** and **5F, Table 7**). Likewise, relative solvent accessibility values derived from solvent accessible surface area (SASA)^36^ analysis on PDB models are systematically higher and more variable than those from AlphaFold (median RSA for AlphaFold predicted models = 0.19, median RSA for PDB experimental models = 0.33; **Figure 4I, Extended Data Figure 6A, 6B, Tables 9 and 10**). Although AlphaFold models offer near proteome-wide coverage, their monomeric, in-silico nature often yields overly “idealized” geometries that may not reflect the full range of biologically relevant conformations^37–39^. This was exemplified in our secondary structure analysis on AlphaFold models, which showed a greater enrichment of alpha helices than determined in the PDB models (**Figure 4E and Extended Data Figure 6C, Tables 11 and 12**), which is a common feature of AlphaFold structures^40^. To address this, we prioritized experimentally determined PDB structures, trading breadth for depth by focusing on a smaller set of proteins with multiple crystallographic or complexed snapshots that better approximate physiological contexts. In both cases, the experimental data capture greater conformational diversity and exposure compared to the predicted models, which we believe are critical parameters for accurately mapping covalent-probe hotspots.

Transitioning from these broader structural contexts, we sought to dissect the biophysical and functional determinants that govern differential arginine reactivity using datasets of reactive and hyper-reactive arginines (**Figure 5A**). We defined hyper-reactive arginines as those which display similar enrichment at high and low concentrations (comparative ratio of intensity between conditions of less than 2), whereas less reactive arginines display concentration-dependent labeling (comparative ratio of at least 2) (**Figure 5A and Figure 2D**). The median log_2_ ratio of enrichment between 1 mM and 100 µM **Nin-Alk** across all arginines was 2.50, reflecting dose-dependent labeling of most reactive arginines. Notably, 410 sites (∼ 6.0% of detected arginines) were hyper-reactive, fully engaged at 100 µM, while the remaining 6478 sites exhibited clear, concentration-dependent labeling. This percentage of arginines annotated as hyper-reactive is consistent with reports for cysteine and arginine^1,2,4^. Moreover, proteins typically harbor a single hyper-reactive arginine but can contain multiple concentration-dependent reactive arginines (**Figure 5B**).

**Figure 5.**
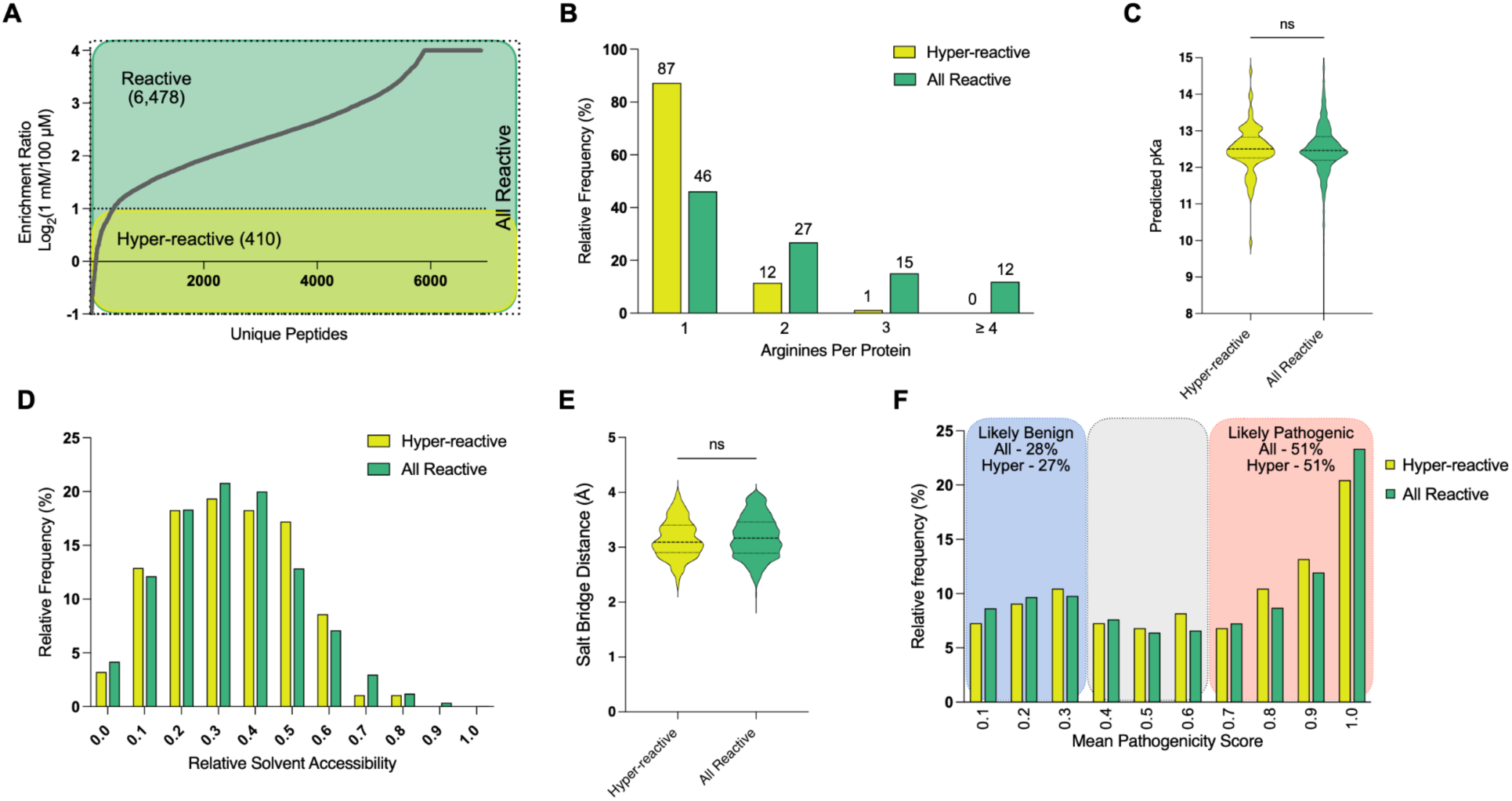
Biophysical and functional features between reactive arginine residues. **A.** Waterfall plot showing the distribution of median RAP enrichment scores (log_2_ heavy-to-light [H/L] ratios) for all quantified arginines across 4 biological replicates. Arginine residues with log_2_(H/L) ≤1 were classified as hyper-reactive (n = 410), representing a subset of the total reactive arginines identified (n = 6,888). See Figure 2D for a schematic of the RAP workflow. **B.** Distribution of reactive and hyper-reactive arginines per unique protein. **C.** Distribution of predicted pKa values for hyper-reactive (n = 85) and all reactive arginine residues (n = 1570), calculated using PROPKA3 and the PDB.^41,42^ Welch’s t-test (two-tailed, parametric) **D.** Predicted relative solvent accessibility (RSA) of hyper-reactive (n = 93) and reactive arginines (n = 1649), computed using the Shrake–Rupley algorithm^36^ on PDB structures, indicating a shared preference for surface exposure(**Table 9**). **E.** Violin plot showing the distribution of salt bridge distances between guanidinium nitrogen atoms of hyper-reactive (n = 101) or all reactive arginines (n = 1445) and carboxylate oxygens of acidic residues (Asp/Glu) in PDB models(**Table 7**). Welch’s t-test (two-tailed, parametric) **F.** Mean pathogenicity score of mutations to each hyper-reactive (n = 220) and reactive arginine (n = 4865), predicted using AlphaMissense(**Table 14**).^43^

With our lists of either all reactive (any arginine engaging ninhydrin) or hyper-reactive (enrichment ratio less than 2) arginines, we set out to identify distinguishing features between the two sets. Even though arginine remains protonated under physiological conditions, cysteine and lysine reactivity have been linked to residue pKa values^44,45^, so we first asked whether arginine labeling follows a similar trend. Predicted pKa values for all reactive arginines within the PDB were, as expected, tightly clustered around 12.5. Distributions for hyper-reactive versus all reactive arginine sites showed no significant differences (**Figure 5C and Table 9**), indicating that, unlike lysine (**Extended Data Figure 7B**), intrinsic protonation state does not drive labeling efficiency. This finding was consistent for predicted AlphaFold structures (**Extended Data Figure 7B, Table 13**). Given the established importance of solvent accessibility for cysteine modification by iodoacetamide^46^, we next assessed the relative solvent exposure of reactive arginines. SASA analysis^36^ revealed comparable relative solvent exposure distributions for hyper-reactive and all reactive sites, with both populations exhibiting partial rather than full surface exposure (median RSA for hyper-reactive = 0.31, median RSA for all reactive = 0.33, mean RSA for both = 0.33; **Figure 5D and Table 9**). The pKa and solvent accessibility independence of arginine labeling implies that: arginine-directed probes can be designed to engage arginines across diverse structural and environmental contexts, and selectivity would be based on interactions between the probe’s scaffold and the binding site, not on the intrinsic reactivity of the engaged arginine. This highlights a notable difference compared to cysteine-reactive probe design, wherein pKa and solvent accessibility of targeted cysteines play major roles in the reactivity and selectivity of covalent probes.

To examine whether salt-bridge geometry contributes to hyper-reactivity, we compared the salt bridge distances for hyper-reactive and all reactive arginines in PDB models (**Figure 5E and Table 7**). Both populations exhibit broad, overlapping distributions (median distances = 3.2 Å for both hyper-reactive and all reactive sites), indicating that salt-bridge proximity alone does not distinguish the hyper-reactive subset. Lastly, we explored functional constraints by mapping mean pathogenicity scores^43^ (AlphaMissense) onto each reactive arginine (**Figure 5F and Table 14**). A high pathogenicity score indicates that mutations of a given residue are associated with significant functional impairment or disease-related phenotypes. Both hyper-reactive and all reactive residues are enriched at the high end of the pathogenicity scale, with 51% of enriched arginines falling into the likely pathogenic range, indicating a labeling bias towards arginines that are mutationally constrained, likely reflecting their critical structural or functional roles in maintaining cellular integrity. Since these arginines play critical roles in cellular function, they are less likely to undergo genetic mutations that could evade covalent engagement. Thus, covalent engagement at these essential positions likely mimics genetic perturbation, directly impacting residue functionality and consequently cellular processes, making them attractive and robust targets for drug and tool compound development.

### Potential mechanism for ninhydrin inhibition of aconitase

Ninhydrin has been reported to inhibit the catalytic activity of mitochondrial aconitase at doses comparable to those used in our assays^47^, but the mechanism of this inhibition has not been elucidated. Mitochondrial aconitase plays a central role in the tricarboxylic acid cycle, linking cellular energy production to redox homeostasis and iron-sulfur cluster maintenance^48^. Dysregulation of its activity has been implicated in metabolic disorders, neurodegeneration, and tumor progression^49^. While not widely targeted clinically, mitochondrial aconitase has garnered interest as a metabolic vulnerability in cancer and as a redox-sensitive node in neurodegenerative diseases, with small molecules and metabolic poisons such as fluoroacetate demonstrating proof-of-concept for pharmacological modulation^50^. Our **Nin-Alk** hyper-reactivity RAP dataset revealed four modified arginines on mitochondrial aconitase (R479, R474, R564 and R607) which are all located within or proximal to the enzymes active site playing roles in the potioning of the native citrate/isocitrate. Among these, R479 was consistently detected across all samples and occupies a central position in substrate coordination (**Table 6**). Structural analysis of an aconitase crystal structure (PDB 1B0J)^51^ identified R479 as a key residue in the positioning of aconitase’s native substrate (**Figure 6A**). Covalent docking simulations to this crystal structure further demonstrated that ninhydrin modification at this site would sterically occlude the active site, likely preventing substrate binding (**Figure 6B**). The presence of multiple reactive arginines clustered around the catalytic pocket supports competitive engagement of the ninhydrin scaffold within the same binding region, providing mechanistic insight into how covalent modification of these residues may contribute to the observed inhibitory effect.

**Figure 6.**
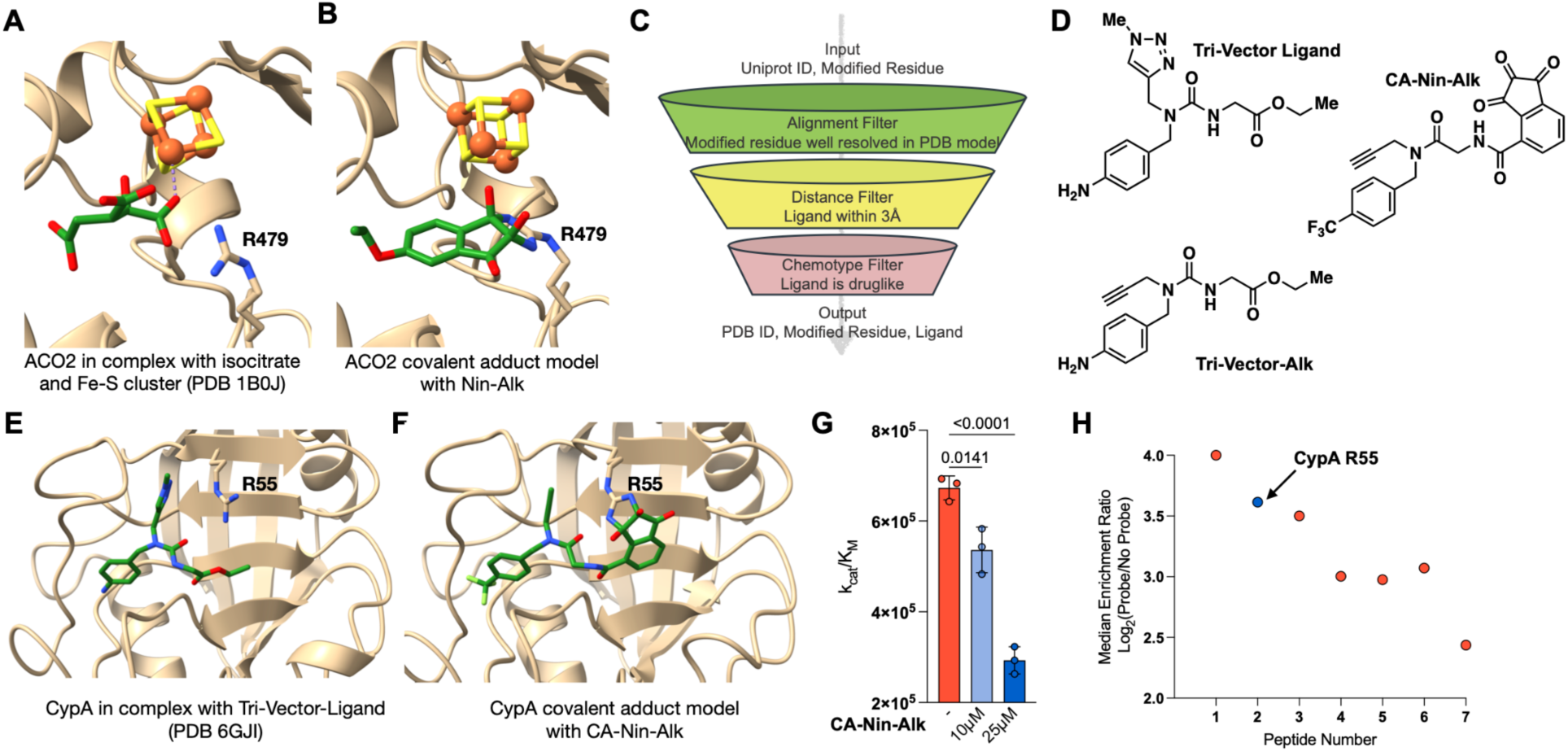
Structure-guided development of arginine-selective ligands. **A.** Crystal structure of mitochondrial aconitase (ACO2) bound to isocitrate and a [4Fe-4S] cluster, highlighting the proximity of R479 to the active site (PDB: 1B0J). **B.** Covalent docking model of **Nin-Alk** to R479 of ACO2, showing spatial alignment with the native isocitrate ligand. **C.** Schematic of the in-house structure-based filtering pipeline used to identify candidate ligands in proximity to modified arginines. Input UniProt IDs and modified residue positions were filtered by PDB alignment, ligand proximity (≤3 Å), and drug-likeness of the ligand to prioritize hits for follow-up. **D.** Chemical structures of the tri-vector ligand and tri-vector alkyne, a clickable analog previously shown to bind cyclophilin A (CypA). The tri-vector alkyne contains an alkyne moiety that enables CuAAC-based enrichment and inspired the synthesis of **CA-Nin-Alk**. **E.** Published co-crystal structure of CypA bound to the tri-vector ligand (PDB: 6GJI),^52^ showing R55 positioned at the periphery of the ligand binding site. **F.** Covalent docking model of **CA-Nin-Alk** to R55 of CypA, showing similar binding orientation as the tri-vector ligand. **G.** CypA activity measured by a coupled peptidyl-prolyl isomerase assay. Pre-incubation of 1 µM recombinant CypA with **CA-Nin-Alk** (10 µM, 25 µM; 1 h) decreases CypA catalytic efficiency (k_cat_/K_m_) by 20.3 ± 8.1% and 56.4 ± 4.8%, respectively compared to DMSO. Data represent mean ± s.d. (n = 3). **H.** Chemoproteomic enrichment of peptides labeled by **CA-Nin-Alk** (10 µM, 1h) in a RAP experiment. Each data point represents a distinct modified peptide; covalent labeling of R55 in cyclophilin A (CypA) is highlighted in blue

### Development of a pipeline for ligand-directed arginine modification

To systematically identify which reactive arginines in our RAP dataset would be amenable to targeting with ninhydrin-endowed ligands, we developed an in-house computational structural proteomics pipeline (**Figure 6C**). The workflow compiles a list of UniProt IDs^53^ alongside their respective modified arginine residues and cross-references them with available high-resolution structures in the Protein Data Bank (PDB).^42^ The analysis parsed through 9,348 unique PDB files, identifying 33,402 occurrences of arginines from our dataset. For each structurally resolved arginine, the pipeline identifies nearby ligands within 3 Å of the guanidinium group. From these occurrences, 292 arginines were found to engage in 884 unique contacts with 461 distinct ligands (**Table 15**). To prioritize functionally relevant ligands, we exclude primary human metabolites, crystallographic ions, and buffer components.

Applying this strategy, we identified R55 in cyclophilin A (CypA) as a reactive arginine proximal to recently reported tri-vector non-covalent binders^52^ that provide synthetically tractable starting points for medicinal chemistry (**Figure 6D** and **6E**). However, we note that while these molecules have been previously reported to non-covalently engage CypA, they have not been evaluated for their inhibition of CypA activity. Structure-guided ligand optimization led to the development of **CA-Nin-Alk** (**Figure 6D**), where the ethyl ester was replaced with a ninhydrin warhead, the urea was substituted with an amide, and the aniline NH₂ was replaced with an isosteric trifluoromethyl group. This substitution leveraged the trifluoromethyl group’s ability to electronically and spatially mimic an aniline while avoiding potential liabilities of a free amine noted in the original tri-vector ligand report, thereby maintaining the binding geometry observed in the tri-vector ligands. This design positioned the ninhydrin warhead appropriately for covalent engagement with R55 in our docking models to the crystal structure of cyclophilin A in complex with a tri-vector ligand (PDB 6GJI)^52^ (**Figure 6F**). An *in vitro* washout-like activity assay revealed that **CA-Nin-Alk** attenuated CypA catalytic activity^54^. Using a modified coupled peptidyl-prolyl isomerase (PPIase) assay with the chromogenic substrate suc-AAPF-pNA, recombinant CypA (1 µM) was pre-incubated with **CA-Nin-Alk** (10 µM or 25 µM; 1 h) or vehicle and subsequently diluted 100-fold into the reaction mixture for a final CypA concentration of 100 nM immediately before initiating catalysis. Importantly, this dilution step would be expected to dissociate noncovalent complexes^55^ thereby mimicking a washout step, which would help indicate whether the interaction was covalent or non-covalent. Under these conditions, **CA-Nin-Alk** treatment resulted in a dose-dependent decrease in catalytic efficiency (**Figure 6G**, **Table 16 and Supplemental Figure 3**). Specifically, **CA-Nin-Alk** treatment at 10 µM and 25 µM reduced (k_cat_/K_m_) by 20.3 ± 8.1% and 56.4 ± 4.8%, respectively, relative to DMSO. These findings suggest that **CA-Nin-Alk** is either capable of covalently inhibiting CypA enzymatic activity or can function as a non-covalent inhibitor at 100 nM.To quantitatively assess the proteome-wide selectivity of **CA-Nin-Alk** and validate covalency for CypA at R55, we performed a probe-versus-no-probe RAP experiment at two concentrations. At 100 µM, **CA-Nin-Alk** enriched 287 modified peptides, indicating that **CA-Nin-Alk** selectively modifies a limited subset of reactive arginines under physiologic conditions (**Table 17**). At 10 µM, only 7 modified peptides were detected, demonstrating a marked reduction in off-target engagement at lower concentrations (**Figure 6H** and **Table 18**). Notably, R55 of cyclophilin A was among these 7 peptides. The ligand scaffold likely plays a key role in directing this selectivity by engaging the binding pocket containing R55 and restricting the modification of other reactive arginines throughout the proteome. Together with the reduction in CypA catalytic efficiency observed in the PPIase assay (**Fig 6G**), these results suggest that **CA-Nin-Alk** engages CypA through its active-site arginine, consistent with a covalent mode of inhibition.

A comparative RAP analysis at 1 mM between **CA-Nin-Alk** and its regioisomer **iso-CA-Nin-Alk**, which repositions the ninhydrin warhead away relative to R55, revealed both a significantly reduced overall reactivity for **iso-CA-Nin-Alk** at CypA R55 and a median log_2_ selectivity between the probes ratio of 2.49 in favor of **CA-Nin-Alk**, likely driven by a combination of improved interactions with the binding pocket and better electrophile placement for improved engagement (**Extended Data Figure 8** and **Table 19**).

## DISCUSSION

Covalent probes have transformed chemical biology and drug discovery, but broadly expanding covalent strategies to amino acids beyond cysteine and lysine has been challenging. Our study establishes ninhydrin-based probes as selective and robust tools for covalent arginine labeling, significantly expanding the chemical proteomics toolkit and enabling new approaches to study arginine-mediated processes. Our ninhydrin-derived probes leverage the equilibrium between the hydrate and reactive dehydrate states of vicinal dicarbonyls. This strategy optimizes probe reactivity while minimizing non-specific interactions and positions arginine as a promising target for broadly applicable covalent strategies. Although **Nin-Alk** demonstrates more complete proteome engagement than **PGO** (**Figure 3A, 3B**), the latter will be valuable in contexts where selective or attenuated reactivity is advantageous. **PGO**’s greater rotational freedom and lower intrinsic electrophilicity suggest that it could function analogously to chloroacetamides in cysteine-directed ABPP, providing a less globally-reactive warhead that enables competition-based ligand screening or focused mapping of highly nucleophilic arginines. Future efforts might leverage this tunability in designing arginine-targeting covalent inhibitors with site-specific, tailored electrophilicity.

Unlike cysteine and lysine, whose reactivities heavily depend on intrinsic residue pKa and high solvent accessibility, arginine residues demonstrate consistent reactivity independent of pKa variations or extensive solvent exposure. This independence could be advantageous or disadvantageous for probe development: it allows for covalent engagement in structurally diverse and otherwise challenging environments, but privileged intrinsic reactivity is minimal for arginines compared to cysteines. Cell lysate labeling studies revealed that modified arginines are often partially buried rather than fully solvent-exposed, emphasizing that determinants of arginine reactivity remain poorly defined, unlike other amino acids, where factors such as pKa or solvent exposure more clearly predict reactivity (**Figures 1C, 4I and 5D**).

Arginines are frequently localized at protein-protein interaction (PPI) sites, where their covalent modification is poised to directly modulate or disrupt critical interaction surfaces. Both cation-π interactions and salt bridges are more prevalent in proteins from thermophilic organisms, enhancing protein stability. We observe that reactive arginines frequently engage in these stabilizing interactions, although these features were not enriched in hyper-reactive arginines compared to generally reactive ones. This trend is consistent with the absence of significant differences in predicted pKa or solvent accessibility between the two groups. In our RAP dataset, we identify 290 unique arginines stabilizing PPIs within the PDB, forming 519 unique PPI pairs. Among these, 145 arginines participated in 193 cation-π interactions, and 212 arginines were involved in 326 unique salt bridge pairs (**Table 7**). Despite the prevalence of these specific interactions, our analysis found that fully solvated arginines are underrepresented based on their physicochemical properties. This rarity may partly reflect the inherent dynamic nature of fully solvated arginines, as their positions might only be reliably resolved when stabilized at interaction interfaces, resulting in relatively lower RSA values being captured in our bioinformatic analyses. Importantly, these interactions provide valuable insights into structural contexts that could guide rational optimization of arginine-targeted covalent probes.

Integration of experimental PDB structures significantly enhanced our structural analyses compared to predictions based on computational AlphaFold models. Although AlphaFold offers extensive near-proteome-wide coverage, these models frequently depict idealized conformations, lacking critical contextual interactions present in biological environments. By contrast, the use of PDB structures, though computationally more intensive, owing to the need for extensive sequence alignments, the handling of large cryo-EM complexes, and the presence of multiple structural entries for individual proteins, provided more realistic and biologically representative data. Specifically, experimental structures captured greater diversity in salt bridge geometries, solvent accessibility, and interaction distances(**Extended Data Figure 9A and 9B**), all crucial for accurately predicting arginine reactivity(**Figure 4H, 4I, and Extended Data Figures 5B – 5F, 6B, 7A and 7B**). This approach underscores the value of incorporating experimentally determined structures when studying residue-level reactivity, particularly for interpreting complex biophysical features that may be underrepresented in predicted models.

Recent work studying iodoacetamide-cysteine reactivity found that simple features such as 2D sequence, pKa or even data derived from AlphaFold models were not sufficient to predict a cysteine’s relative reactivity. More accurate prediction required experimentally determined structures to start to model residue-level reactivity^56^. While AlphaFold offers transformative accessibility and continues to enable exciting applications such as predictive covalent docking, the current generation of models often struggles with accurate representation of sidechain packing and interaction geometries. As we expand our platform to profile reactive arginines across more cell types, the growing bulk of arginine reactivity data may make AlphaFold-derived structures increasingly useful at scale, particularly given the computational demands of mining the PDB. Future iterations of Alphafold, which are showing improvements in sidechain accuracy, may further enhance its utility for predicting covalent probe engagement.

Modification of functionally relevant arginines is exemplified by ninhydrin-mediated labeling of aconitase at multiple arginines necessary for the positioning of its native substrate. Previous work has demonstrated that ninhydrin inhibits aconitase catalytic activity, and the RAP data extends this hypothesis by providing a putative molecular mechanism of inhibition(**Figure 5A and 5B**). Structural analysis confirmed that R479 is critical for substrate positioning, and covalent modeling indicated that modification at this site sterically occludes substrate binding, contributing to inhibition. These findings highlight the potential of arginine modification as a powerful approach for probing enzyme function and developing covalent inhibitors targeting active-site arginines.

Beyond aconitase, the selective modification of R55 at cyclophilin A with **CA-Nin-Alk** at 10 µM further underscores the utility of ninhydrin-based probes in ligand-directed covalent targeting (**Figure 6**). The rapid development of a ∼22 μM inhibitor of cyclophilin A was achieved with the design of a single lead molecule, highlighting the efficacy of synergizing covalent chemical probes and structural biology to facilitate rapid tool compound development.

Structural comparisons with a regioisomeric probe (**iso-CA-Nin-Alk**), which mispositioned the ninhydrin warhead relative to R55, emphasized the importance of precise warhead placement in achieving selective, site-specific arginine engagement **(Extended Data Figure 8)**. The selectivity demonstrated between the regioisomeric pairs likely is assisted by subtle differences in binding affinity and probe conformation as well as proper electrophile placement. These results demonstrate how integrated computational modeling, proteomics, and structure-guided ligand design can systematically guide the development of selective covalent ligands.

This study establishes a robust framework for selective covalent arginine modification using ninhydrin-derived probes. We introduce RAP as a chemical proteomic approach to classify arginine reactivity, facilitating rational probe development and the identification of key arginines for therapeutic targeting. Future research will focus on refining probe specificity, elucidating additional structural determinants of arginine reactivity, expanding the application of ninhydrin-based probes to a broader range of therapeutically relevant targets, and leveraging the RAP platform as a competitive screening tool for the discovery of arginine-reactive small molecules. Importantly, the ability to perform competitive screening without requiring enrichment handles will simplify the evaluation of candidate molecules, enhance throughput and reduce synthetic complexity, thus accelerating the development of novel arginine-targeted therapeutics. By advancing strategies for arginine-selective covalent modification, this work paves the way for novel chemical biology tools and next-generation covalent therapeutics.

## Supporting Information

The following files are available free of charge.

Supporting Information (PDF including Extended and Supplemental Figures and Methods) Data tables (Excel)

## Author Contributions

A.K.E. conceived of the idea, synthesized molecules, designed and performed experiments, analyzed and interpreted the data, and wrote and edited the manuscript. A.L. designed and performed experiments, analyzed and interpreted the data, and wrote the manuscript. C.S.F. performed experiments, analyzed, and interpreted the data. J.L.M. designed experiments and generated figures. M.T. participated in idea conception and synthesized molecules. I.B.S. and B.W.Z. oversaw the work, designed experiments, analyzed and interpreted the data, and wrote and edited the manuscript.The manuscript was written through contributions of all authors. All authors have given approval to the final version of the manuscript. ‡These authors contributed equally.

## ACKNOWLEDGMENT

We thank members of the Seiple and Zaro labs for helpful discussions, support, and feedback. We thank James Fraser for sharing his cyclophilin A activity assay protocol and spectrophotometer. We thank the Mass Spectrometry facility at Scripps Research for providing Intact Protein analysis. We thank the Automated Synthesis Facility for providing HR-MS analysis.

## ABBREVIATIONS

Nin-Alk: Nihydrin-Alkyne
PGO: phenylglyoxal alkyne
RAP: reactive arginine profiling
CypA: cyclophilin A

## SYNOPSIS

This work describes a selective method to covalently engage arginine residues in proteins, revealing thousands of reactive sites across the proteome and new opportunities for modulating protein function.

## Extended Data

**Extended Data Figure 1.**
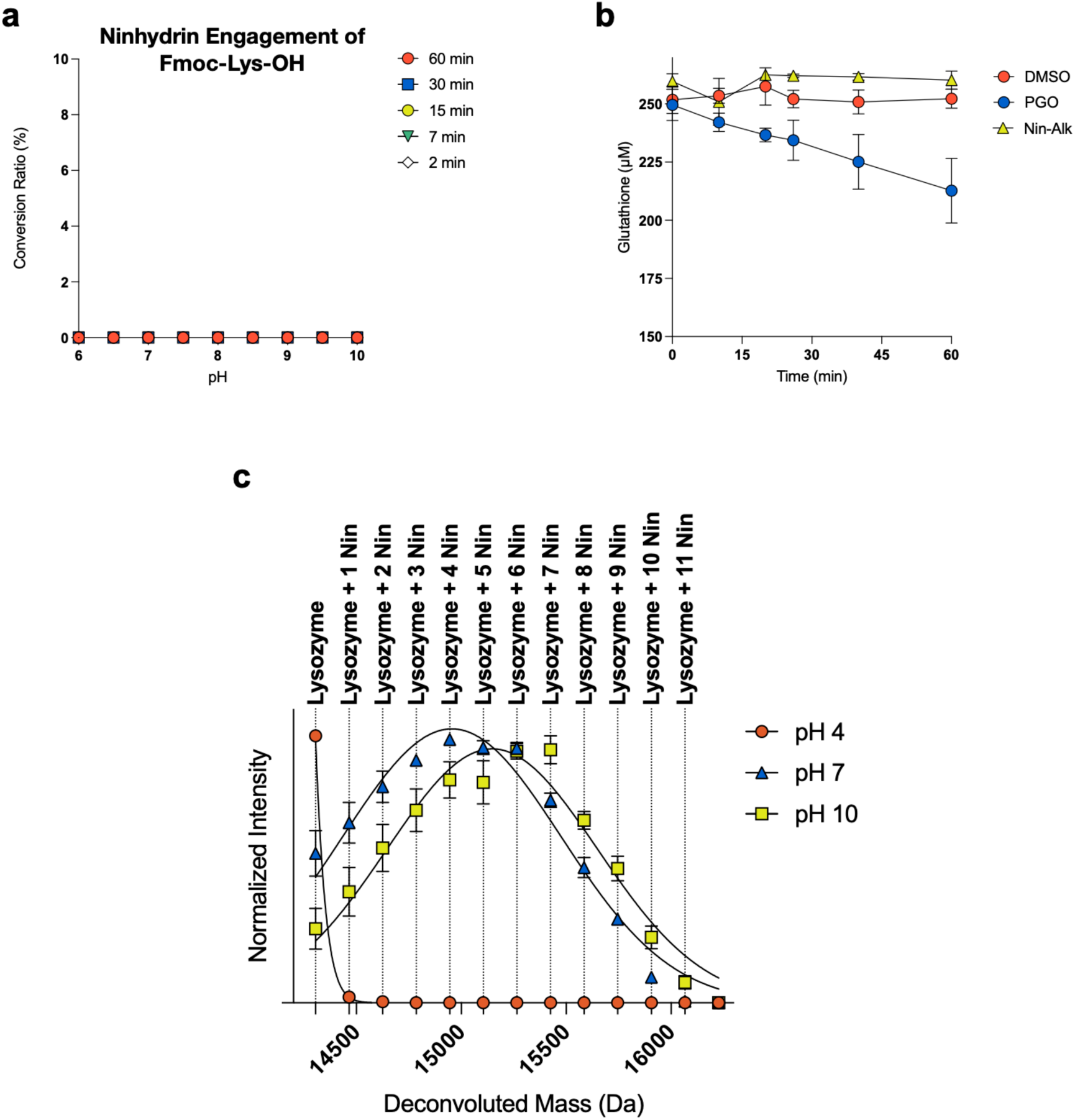
Evaluation of ninhydrin as a covalent modifier of lysine and free thiols. **A.** Kinetic analysis of pH- and time-dependent consumption of Fmoc-Lys (500 µM) by ninhydrin (15 mM, 30x) in PBS at 20 °C. Lysine consumption was quantified by LC-MS over 2 to 60 minutes. Error bars represent SD. n = 3. **B**. Glutathione reactivity assay for arginine reactive warheads. Compounds were incubated with reduced glutathione (250 μM) in assay buffer (100 mM Tris pH 9.0, 50% MeOH) at a final concentration of 500 μM. At the indicated time points (0–60 min), Ellman’s reagent was added and residual free thiols were quantified via absorbance at 412 nm. Values reflect means of triplicate measurements with individual replicates shown. DMSO control (orange circles), phenylglyoxal (blue circles), **Nin-Alk** (yellow triangles). **C.** Assessment of ninhydrin engagement of lysozyme. Lysozyme is covalently modified by ninhydrin. Intact protein mass spectrometry analysis of Lysozyme (1.5 µM) and 100 µM ninhydrin at 37 °C for 30 minutes across pH 4.0, 7.0 and 10.0. Reactions were quenched by dilution into 0.1% aqueous TFA and analyzed by LC–MS on a Agilent 6230 TOF system. N =3 Error bars represent SEM. Representative deconvoluted mass spectra in SI.

**Extended Data Figure 2.**
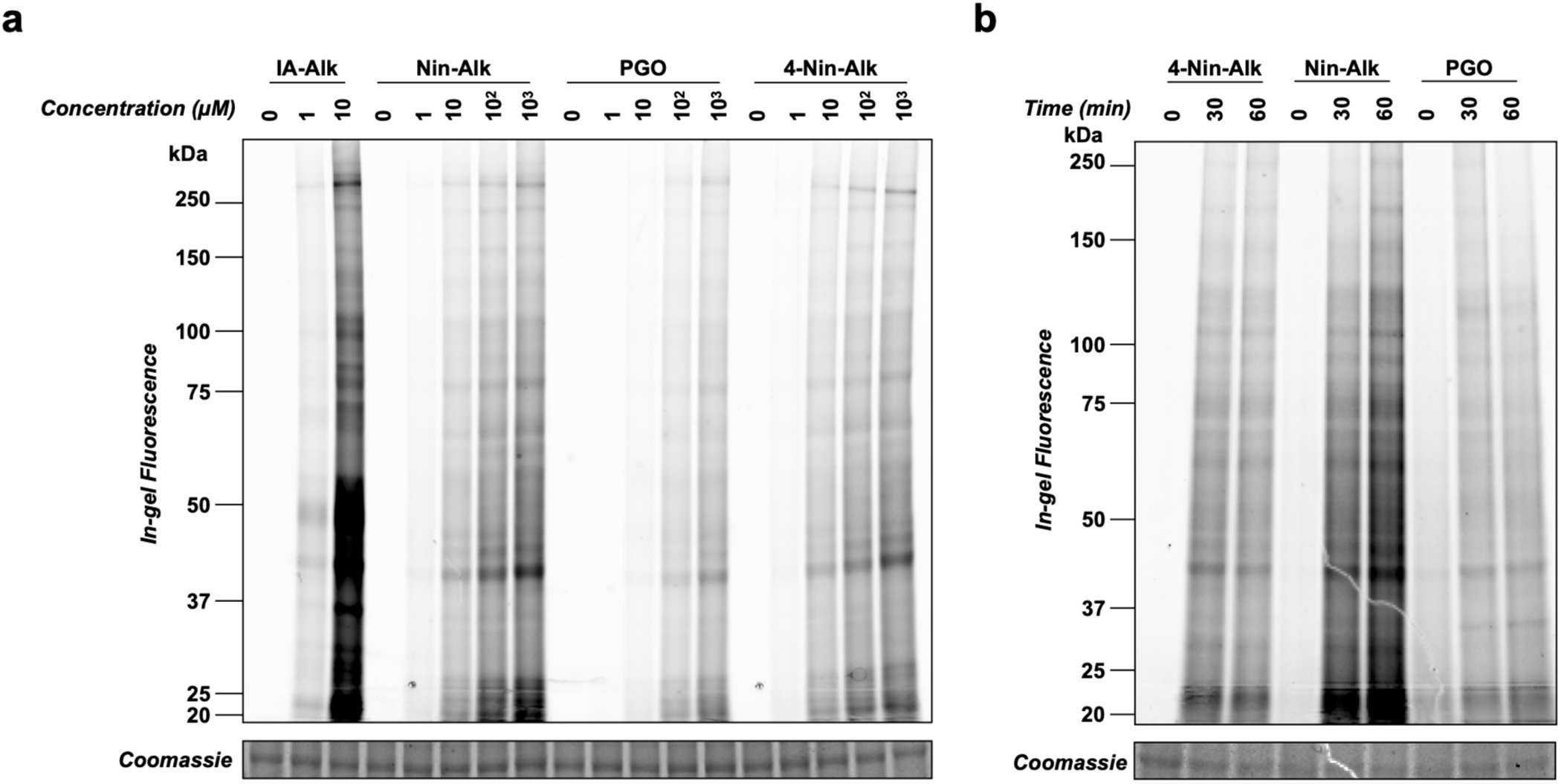
In-gel fluorescence scanning of cysteine and arginine reactive probe labeling in lysates. **a.** Qualitative, gel-based assessment of proteome-wide reactivity of alkyne-bearing covalent probes in Mino cell lysate. Soluble proteome (1.5 µg µL⁻¹) was treated with each probe at the indicated concentration for 1 h, followed by CuAAC conjugation with TAMRA-N₃, separation by SDS–PAGE, and in-gel fluorescence scanning. **b.** Qualitative time-dependent reactivity of indicated probes in live cells. Mino cells were treated with 100 µM of the indicated probe for 0, 30, or 60 min, followed by lysis, CuAAC conjugation with TAMRA-N₃, separation by SDS–PAGE, and in-gel fluorescence scanning. Uncropped gel images available in SI.

**Extended Data Figure 3.**
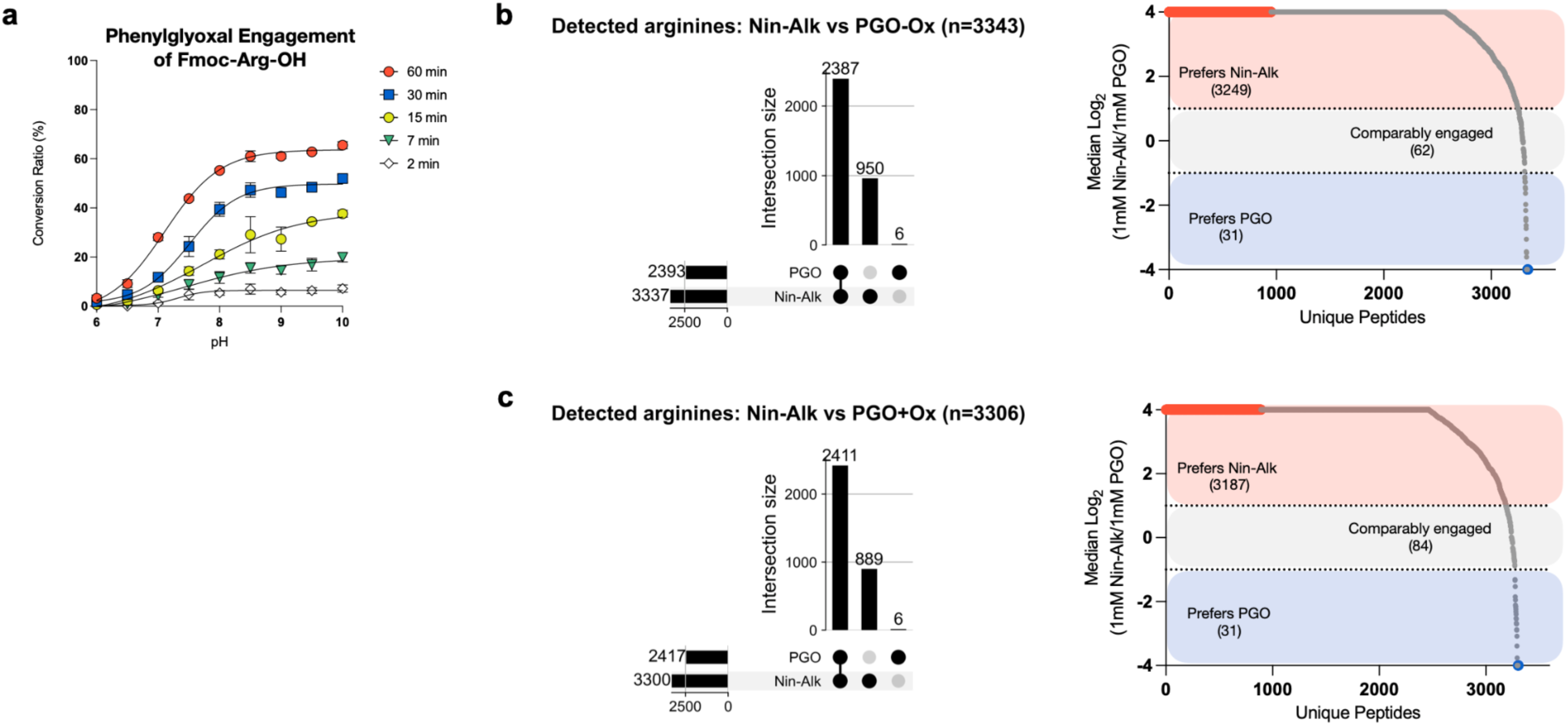
**a.** Kinetic analysis of pH- and time-dependent consumption of Fmoc-Arg (500 µM) by PGO (15 mM, 30x) in PBS at 20 °C. Arginine consumption was quantified by LC-MS over 2 to 60 minutes. Error bars represent SD. n = 3 **b.** UpSet and waterfall plots for the closed search of the non-oxidized **PGO**-engaged and **Nin-Alk** engaged arginines detected by RAP. High overlap was observed between **PGO** and **Nin-Alk**–engaged arginines (n = 2,387), with the majority preferentially engaged with a log_2_ ratio ≥ 1 by **Nin-Alk** (n = 3,249) relative to **PGO** (n = 31). **c.** Corresponding analysis for the closed search including the oxidized **PGO** species (+ox). Similar trends were observed (n = 3,306 total), with a higher preference for **Nin-Alk** (n = 3,187) relative to **PGO** (31).

**Extended Data Figure 4.**
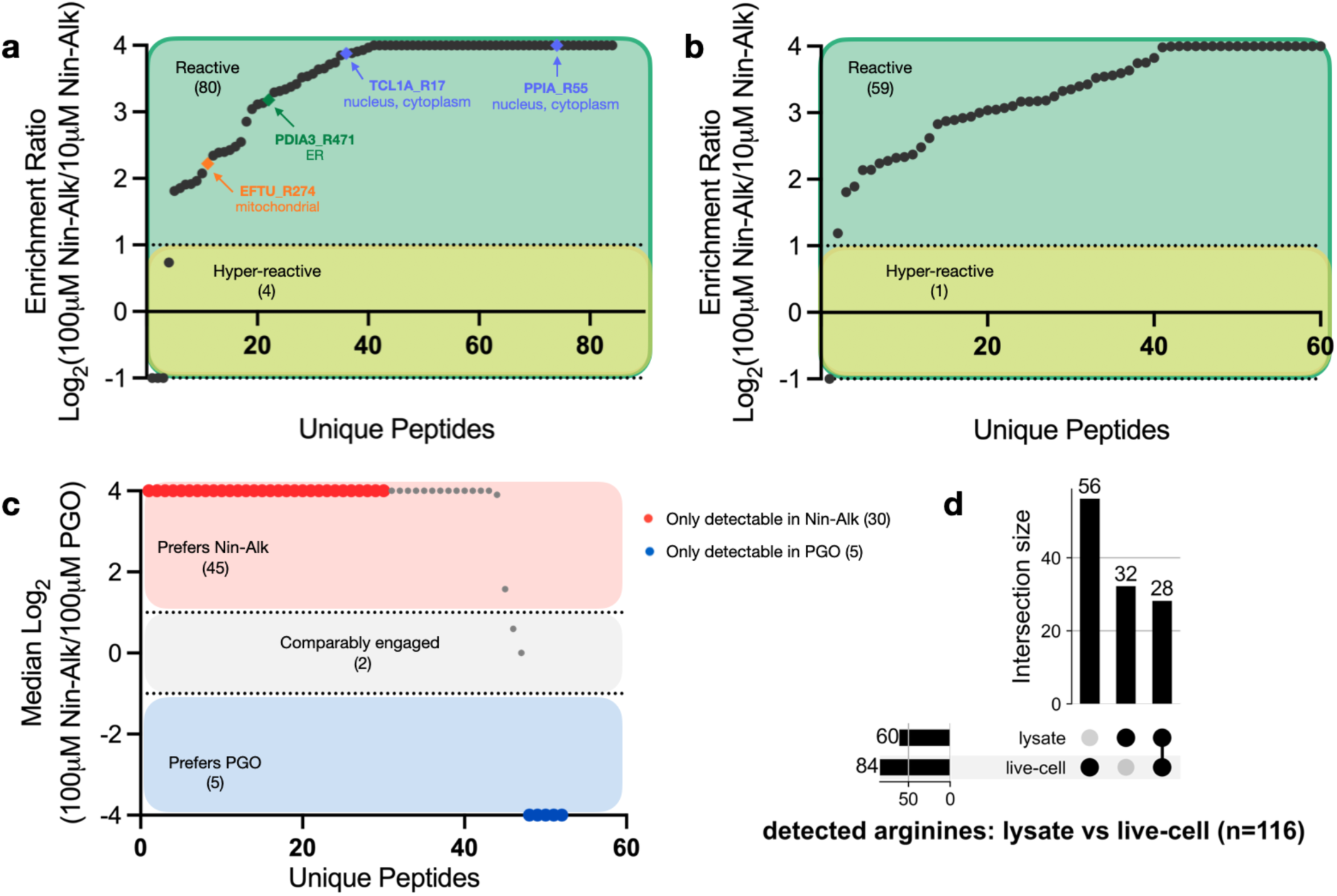
Live-cell RAP profiling in Mino cells. **a.** Waterfall plot showing the distribution of median RAP enrichment ratios (log_2_100 μM **Nin-Alk**/10 μM **Nin-Alk**) obtained from live-cell labeling in Mino cells. Arginines with log_2_ ≥ 1 were characterized as reactive (n = 80), while those with a log_2_ < 1 were designated hyper-reactive (n = 4). Representative proteins with their annotated subcellular localizations (UniProt) are highlighted, demonstrated labeling of arginines from distinct cellular compartments including the nucleus/cytoplasm (TCL1A, PPIA), endoplasmic reticulum (PDIA3), and mitochondria (EFTU). **b.** Waterfall plot showing distribution of median RAP enrichment ratios for lysate labeling of Mino cells at 100 μM and 10 μM of **Nin-Alk**. Despite the absence of cellular barriers and growth media, overall coverage remained modest (n = 60 reactive arginines), suggesting that the lower number of labeled sites observed in live-cell experiments is not solely attributable to probe permeability or intracellular accessibility. **c.** Waterfall plot comparing distribution of RAP enrichment ratios for **Nin-Alk** (100 µM) and **PGO** (100 μM) labeling in live Mino cells. Peptides with log_2_ ≥ 1 were classified as preferring **Nin-Alk** (n = 45), whereas those with log_2_ ≤ −1 preferred **PGO** (n = 5). These results mirror the trends observed in lysate (**Figure 3B**). **d.** UpSet plot showing overlap between arginine-modified peptides detected in lysate and live-cell experiments (n = 116 total). 28 peptides were common to both datasets, suggesting potential context-dependent factors driving arginine accessibility and reactivity.

**Extended Data Figure 5.**
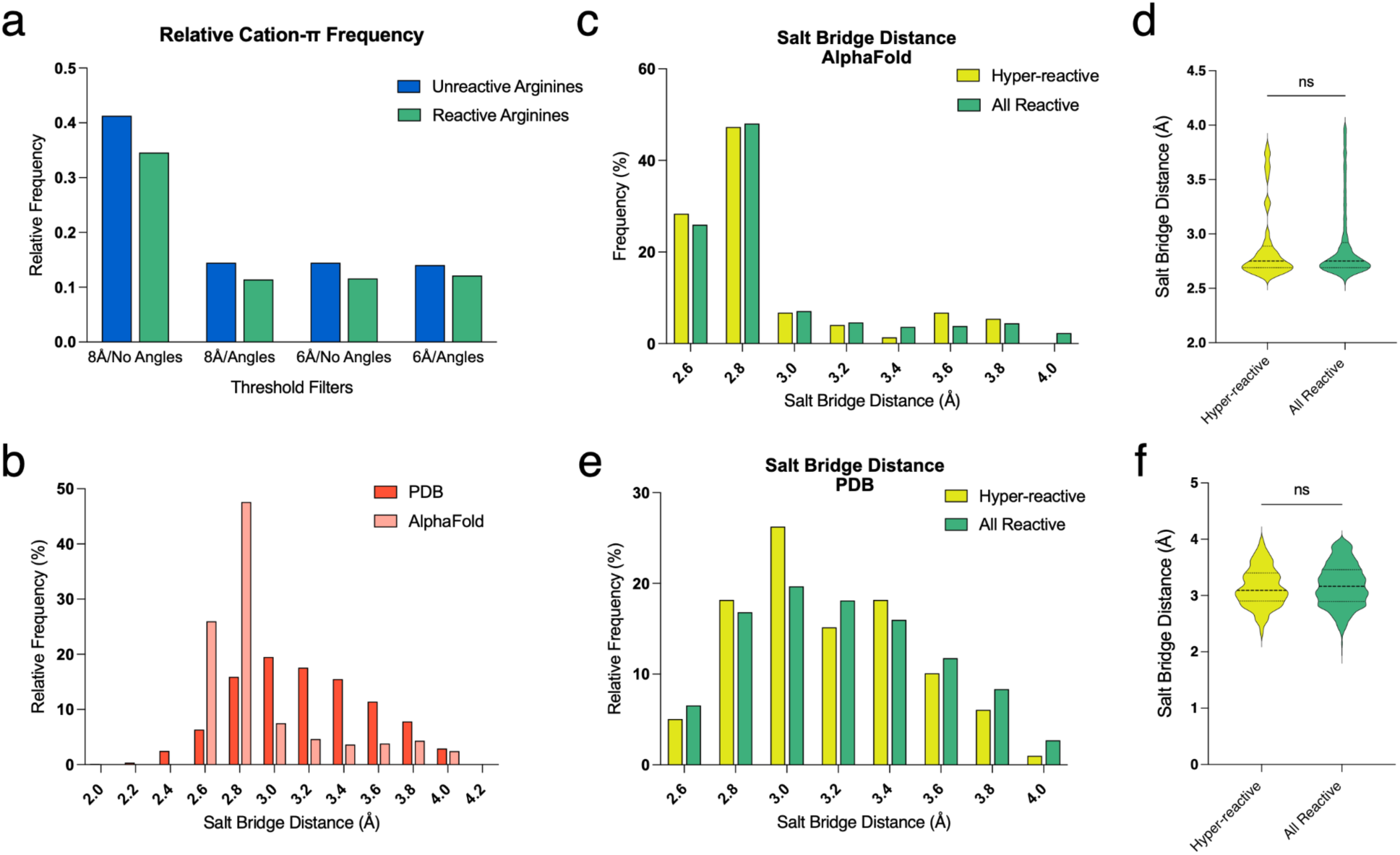
Comparative analysis of intermolecular interactions surrounding reactive arginines. **a**. Frequency of arginines engaged in cation–π interactions across reactive (green) and unreactive (blue) datasets, based on AlphaFold-predicted models and varying distance (6 Å or 8 Å) and angle filters. No significant enrichment or depletion of cation–π interactions was observed among reactive residues. **b**. Comparison of salt bridge distances between reactive arginines and acidic residues in AlphaFold (light coral, n = 1594) versus experimentally determined PDB structures (terra cotta, n = 1445), showing generally shorter predicted distances in AlphaFold models. **c**. Histogram of salt bridge distances from AlphaFold models comparing hyper-reactive (yellow, n = 74) and all reactive (green, n = 1594) arginines. **d**. Violin plot showing the distribution of salt bridge distances from AlphaFold models for hyper-reactive and all reactive arginines; no statistically significant difference observed (unpaired t-test). **e**. Histogram of salt bridge distances derived from experimental PDB structures comparing hyper-reactive (yellow, n =101) and all reactive arginines (green, n = 1445). **f**. Violin plot of salt bridge distances in PDB models; consistent with AlphaFold data, no statistically significant difference was observed between hyper-reactive and all reactive arginines.

**Extended Data Figure 6.**
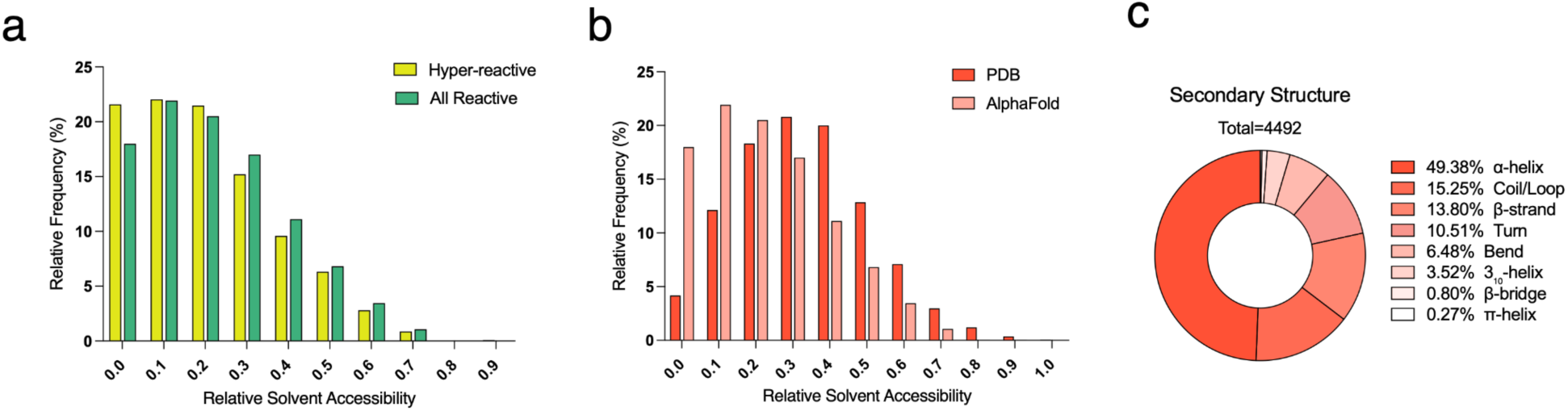
Solvent accessibility and secondary structure profiles of reactive arginines. **a.** Distribution of relative solvent accessibility (RSA) for hyper-reactive (yellow, n = 1960) versus all reactive (green, n = 4309) arginines in AlphaFold-predicted structures. Both sets show a similar solvent exposure profile. **b.** Comparison of RSA values for all reactive arginines across AlphaFold (light coral, n = 4309) and PDB (terra cotta, n = 1649) models reveals a modest shift toward higher accessibility in AlphaFold predictions. **c.** Secondary structure distribution of all reactive arginines based on DSSP annotation of AlphaFold models (n = 4492 residues). The majority localize within α-helices (49.38%), followed by coil/loop regions and β-strands.

**Extended Data Figure 7.**
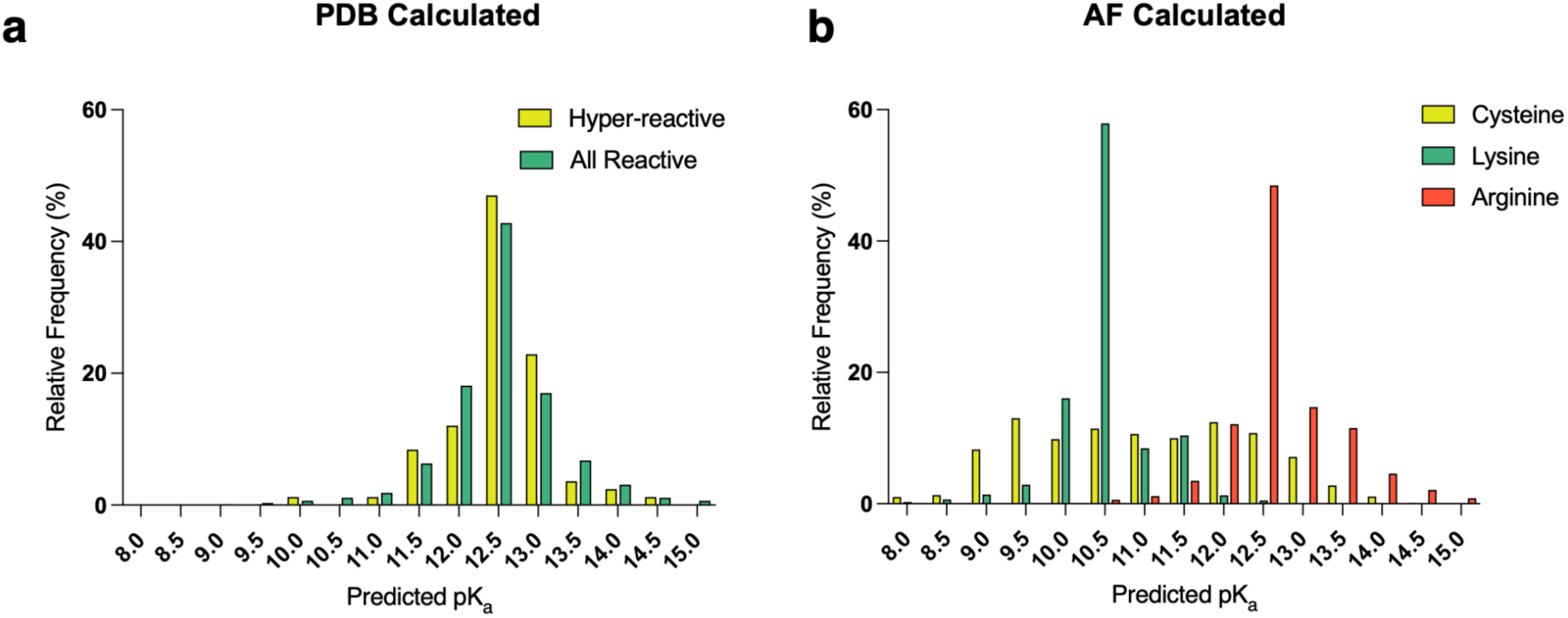
Predicted pKa distributions of reactive residues. **a.** Distribution of PROPKA-predicted pKa values for reactive arginines calculated using experimental PDB structures, comparing hyper-reactive (yellow, n = 85) and all reactive (green, n = 1570) residues. **b.** Predicted pKa distributions for reactive arginines(orange, n = 4334), cysteines(yellow, n = 3988), and lysines (green, n = 3863) using AlphaFold structures. Arginine predictions are from the current dataset, cysteine data was based on matched cell analysis with Mino cell lysate and iodoacetamide alkyne as the electrophile and lysine data are derived from previously reported reactivity profiles from Hacker et al. (2017)^4^. No significant pKa shift distinguishes hyper-reactive from reactive residues across any residue class.

**Extended Data Figure 8.**
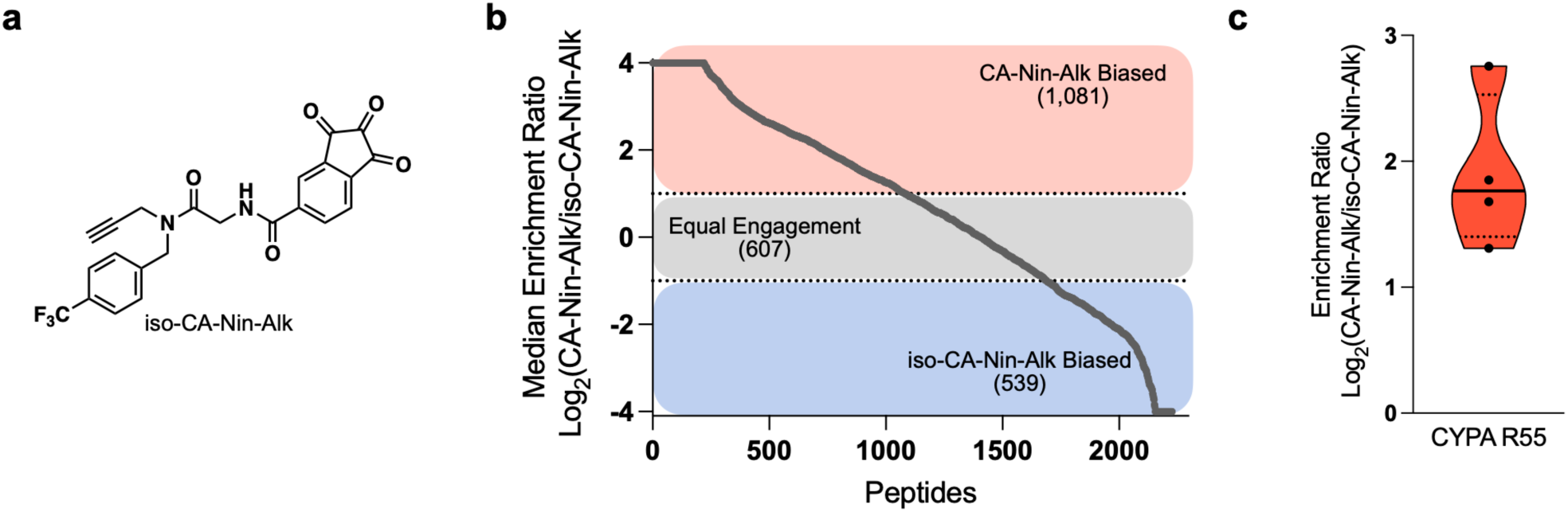
Exploration of a regioisomer of CA-Nin-Alk. **a.**Chemical structure of the tri-vector alkyne **iso-CA-Nin-Alk**, a clickable analog previously shown to bind cyclophilin A (CypA) **b.** Waterfall plot depicting the regioselective enrichment of peptides labeled by **CA-Nin-Alk** and **iso-CA-Nin-Alk** in Mino cell lysate. Peptides identified by the RAP workflow were classified based on relative enrichment at 1 mM probe concentration in biological triplicates. Each point represents the log₂ ratio of **CA-Nin-Alk** to **iso-CA-Nin-Alk** enrichment for an individual peptide. Peptides were grouped into **CA-Nin-Alk** biased (red, n = 1,081), **iso-CA-Nin-Alk** biased (blue, n = 539), and equally engaged (gray, n = 607) categories **c.** Relative enrichment of CypA R55 between **CA-Nin-Alk** and **iso-CA-Nin-Alk** as determined by RAP. n = 4 replicates.

**Extended Data Figure 9.**
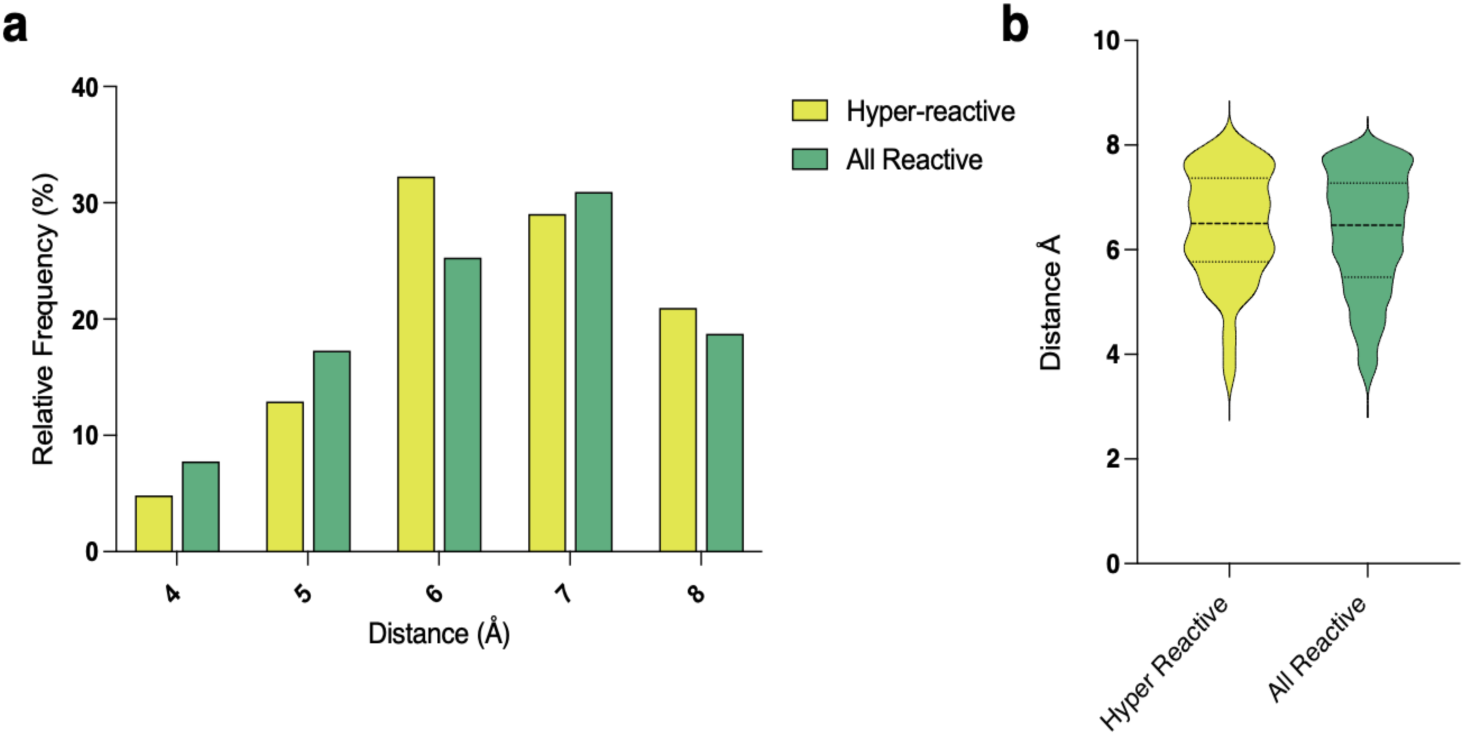
Structural comparison of cation–π interactions at hyper-reactive versus general reactive arginines. **a.** Histogram depicting the frequency distribution of cation–π interactions among hyper-reactive arginines (yellow, n = 62) compared to all reactive arginines (green, n = 905) based on structural analysis of the experimental models from the PDB. **b.** Violin plot showing the same comparison as in (**a**), highlighting the broader distribution of cation–π interactions across the two arginine populations.

## Methods

### Density Functional Theory (DFT) Calculations

Initial molecular models were prepared using Avogadro^57^. All calculations were performed using GAMESS^58^ on ChemCompute.org. Geometry optimization was performed with the PM3 basis set, restricted Hartree-Fock (RHF) molecular orbital method, and B3LYP DFT functional. Solvent effects were modeled using the water PCM solvent model on a single processor core. The optimized structures were subjected to a single-point energy calculation with the 6-311G** basis set. All output models were reviewed in Avogadro for realistic geometry, and the delta G free energy in solvent between the hydrated and dehydrated states (with an additional water molecule) for both phenylglyoxal and ninhydrin was calculated to determine the relative equilibrium constants.

### Fmoc- Arginine and Lysine Kinetics Assay

In triplicate, 190 µL of PBS buffer (pH 6.0 – 10.0, in 0.5 pH unit increments) was allotted into 27 Eppendorf tubes. A 5 µL aliquot of a 20 mM stock solution of Fmoc-Arg-OH(or Fmoc-Lys-OH) in DMSO was added to each tube, mixed thoroughly, and briefly centrifuged to ensure complete pooling of the solution. A 600 mM Ninhydrin solution (1:1 DMSO/DI water) was prepared alongside a 2 M HCl quenching solution. All tubes were pre-opened, and a timer, vortex mixer, waste container, and fresh pipette tips were arranged for efficient handling. Reactions were initiated by adding 5 µL of Ninhydrin solution to the first tube while starting the timer, followed by immediate capping and vortex mixing. This process was repeated every 10 seconds for the subsequent tubes, ensuring all reactions were initiated within 4.5 minutes. At designated time points, the reactions were quenched by sequentially adding 10 µL of 2 M HCl to each tube, following the same 10-second interval sequence as for reaction initiation. Samples were then centrifuged, and 200 µL of the supernatant was carefully transferred using gel-loading tips into pre-labeled LC-MS vials with conical inserts, ensuring minimal air bubble formation. Samples were analyzed using a Waters Acquity UPLC SQD2 detector LC-MS system on a polar gradient method (20–55% acetonitrile/water + 0.1% formic acid). The UV detection window was set to 300 nm for Fmoc absorbance, and free arginine (SM) and adduct formation were identified based on the extracted ion chromatograms. Peak integrations at 300 nm were used to calculate the conversion ratio of reactants to products.

### Cysteine Reactivity Assay

L-cysteine was prepared at 250 µM in assay buffer (100 mM Tris, pH 9.0, 50% MeOH). In triplicate, 100 µL of the cysteine solution was dispensed into each well of a clear, flat-bottom 96-well plate (Costar). Inhibitors were added as 100X stocks (1 µL per well; 500 µM final concentration) or with DMSO vehicle control, and reactions were incubated at room temperature for the indicated times (0-60min). Following incubation, Ellman’s reagent (10 µL of a 50mM stock in DMSO; 5mM final concentration) was added, and plates were incubated for 10 minutes at room temperature. Absorbance was measured at 412nm in a plate reader (Spectramax M5). Free cysteine concentrations were quantified using a standard curve, and second-order rate constants were calculated as previously described^23^.

### Intact Protein Mass Spectrometry

Reactions were performed with 1.5 µM BSA or Lysozyme in PBS, incubated with 150 µM ninhydrin at 37 °C for 30 minutes. Reactions were quenched by a dilution to 0.1% TFA (aq) and analyzed by intact protein LC/MS using an Agilent 6230 TOF LC/MS with a Dual AJS ESI ion source. Agilent 6230 TOF system equipped with an Agilent RLRP-S 1000Å 5 μm 50×2 1MM column. The mobile phase was a linear gradient of 5–90% acetonitrile/water + 0.1% formic acid over 4 minutes with a flow rate of 300 µL/min.

### Glutathione reactivity assay

Reduced glutathione (GSH) was prepared at a concentration of 250 μM in assay buffer (100 mM Tris pH 9.0, 50% MeOH as co-solvent). In triplicate, 100 μL of GSH (250 μM) in assay buffer was added to each well of a clear, flat-bottom 96-well plate (Greiner Bio-one). To each well, 1 μL of 100x inhibitor stocks in DMSO (final concentration 500 μM) or DMSO vehicle was added, and the plate was incubated at room temperature for the indicated time (0–60 min). Following incubation, Ellman’s reagent (Thermo) (10 μL, 50 mM stock in DMSO, final concentration 5 mM) was added, and the reaction was incubated at room temperature for 10 minutes. Absorbance was measured at 412 nm using a plate reader (Cytation 5, BioTek). The remaining concentration of free thiols was quantified using a glutathione standard curve.^23^

### Cell Culture

Mino cells were cultured under standard conditions (37 °C, 5% CO₂) in T175 flasks. Cells were revived from frozen stocks stored in liquid nitrogen. Cryovials containing the cells were rapidly thawed in a 37 °C water bath and immediately transferred to pre-warmed complete culture media. To remove residual DMSO, cells were centrifuged at 300 × g for 5 min, resuspended in fresh culture media, and plated in appropriate flasks. Cells were allowed to recover for 24–48 hours before the first media change and were subsequently maintained under routine culture conditions.

Cells were maintained in non-treated T175 flasks in Cytiva RPMI-1640 medium (no HEPES, stored at 4 °C) supplemented with 10% fetal bovine serum (HyClone), L-glutamine (100 µg mL^−1^), 0.05 mM β-mercaptoethanol, and penicillin-streptomycin (100 μg mL^−1^). Cells were routinely passaged every 2–3 days when they reached a density of ∼8 × 10⁵ cells per mL to maintain exponential growth. Passaging was performed by centrifuging the culture at 300 × g for 5 min, discarding the supernatant, and resuspending the cell pellet in fresh pre-warmed medium at a 1:3 dilution.

Cells were harvested after three passages to ensure robust growth and adaptation to culture conditions. Mino cells were collected from suspension culture, pelleted at 300 × g for 5 min, and washed once with ice-cold PBS. Washed cell pellets were resuspended in ice-cold Dulbecco’s phosphate-buffered saline (DPBS) and sonicated on ice (2 × 20 pulses, 20% amplitude, 1 s pulse rate) using a tip sonicator. The lysed suspension was centrifuged at 21,000 × g for 20 min at 4°C, and the supernatant was transferred to a lo-bind tube. Protein concentrations were determined using the bicinchoninic acid (BCA) assay (Thermo Scientific), and the resulting lysate was used fresh for subsequent experiments.

### Gel-Based Probe Reactivity Profiling

For screening reactivity in lysate, 50 µL of freshly prepared cell lysates (∼1.5 mg mL^−1^ total protein) were treated as indicated in PBS buffer (pH 7.4) at room temperature. Lysates were dosed with a 100X probe in DMSO or an equal volume of DMSO as a control. Following probe treatment, CuAAC click chemistry was performed by sequentially adding 2 μL TAMRA-azide (5 mM, stock in DMSO), 2 μL tris(2-carboxyethyl)phosphine hydrochloride (TCEP) (50 mM, freshly prepared in water), 6 μL tris[(1-benzyl-1-H-1,2,3-triazol-4-yl)methyl]amine (TBTA) (1.7 mM, 3:1 tert-butanol:DMSO), and 2 μL CuSO4•5H2O (50 mM in water) with vortexing between additions. The reaction was incubated at room temperature for 1 h with gentle mixing.

For live-cell screening, 10^6^ cells were plated in 3 mL of complete cell growth medium (as described above) in a single well of a 12-well culture plate (Costar). Wells were treated with either 1000X of either probe or vehicle (0.1% v/v) for 1h under standard conditions (37°C, 5% CO_2_). Immediately following labeling, cells were washed 3X with 5 mL DPBS, pelleted, and flash-frozen in LN_2_ and stored at −80°C overnight. The following day, pellets were lysed and protein concentrations were normalized to 50 µg (1 mg/mL). Click reactions were then performed under the same conditions as above.

Reactions were quenched by adding 250 μL of ice-cold methanol, followed by incubation at −20°C for at least 2 hours. Precipitated proteins were pelleted by centrifugation (10,000 × g, 10 min, 4°C), and the supernatant was discarded. Tubes were inverted and allowed to dry for 30 minutes to ensure complete evaporation of methanol. Protein pellets were resuspended in 75 μL of 4% SDS in PBS (pH 7.5) and bath sonicated (35 kHz, 10 min) to achieve complete solubilization. Following resuspension, 25 μL of 4× Laemmli loading buffer (Bio-Rad, containing 10% β-mercaptoethanol) was added, and samples were vortexed gently before boiling at 95°C for 5 minutes. After cooling to room temperature, 20 μg of protein per lane was loaded onto a Tris-Glycine SDS-PAGE gel (Bio-Rad, Criterion TGX) for separation. Fluorescently labeled proteins were visualized using a Bio-Rad ChemiDoc fluorescence scanner, and total protein staining with Coomassie was performed for normalization.

### RAP Workflow

For labeling of cell lysates, 250 µL freshly prepared cell lysate at 2 µg mL^−1^ protein concentration was treated in lo-bind tubes with either 100 µM probe or 1.0mM probe for 1 hr at 37 °C in pairs. Following probe treatment, CuAAC click chemistry was performed by sequentially adding 25 μL isoDTB tags (light tags to the 100 µM treated, heavy tags to the 1.0mM treated, (Vector Labs, CCT-1565-L and CCT-1565-H, 10 mM, stock in DMSO), 25 μL tris(2-carboxyethyl)phosphine hydrochloride (TCEP) (50 mM, freshly prepared in water), 75 μL tris[(1-benzyl-1-H-1,2,3-triazol-4-yl)methyl]amine (TBTA) (1.7 mM, 3:1 tert-butanol:DMSO), and 25 μL CuSO4•5H2O (50 mM in water) with mixing between additions. The reaction was incubated at room temperature for 1 hour with gentle mixing. Protein samples were solubilized by adding 200 µL of 10% SDS in PBS, gently mixed and individually sonicated (35 kHz, 2 min) to dissolve any precipitate formed during the click chemistry reaction. Equal amounts of samples were combined in a 2 mL lo-bind Eppendorf tube and treated with 1 µL of benzonase, followed by incubation at 37°C for 30 minutes at 800 rpm.

For labeling of live cells, 10^7^ Mino cells were concentrated to 3 mL in complete RPMI-1640 (Cytiva). Media was then supplemented with 1,000X stocks of either probe or vehicle (final concentration 0.1% v/v). Cells were kept under standard conditions (37°C, 5% CO_2_) in T25 flasks for 1 hour. Following labeling, cells were immediately washed with 3X 10 mL DPBS, concentrated, flash-frozen in LN_2_, and stored at −80°C overnight. The following morning, cell pellets were then lysed in DPBS and centrifuged (10,000 × g; 10 min) and kept on ice. The soluble fraction was separated, and the remaining precipitated membrane fraction was re-suspended in DPBS. Both fractions were quantified via BCA and concentrations of respective channels were then normalized for click reactions and solubilized under the same conditions as mentioned above.

For protein precipitation and cleanup, SP3 bead purification was employed. 200 µL Sera-Mag SpeedBeads (GE45152105050250, GE65152105050250), were combined in a 1:1 ratio, washed three times with 1 mL water, and resuspended in 250 µL of water before being added to the sample. After incubation at room temperature with shaking at 1000 rpm for 5 minutes, 1 mL of absolute ethanol was added, and the sample was incubated under the same conditions for another 5 minutes. The beads were collected on a magnetic rack, and the supernatant was removed. Beads were washed three times with 1 mL of freshly prepared 80% ethanol.

To prepare proteins for digestion, the beads were resuspended in 500 µL of fresh 2 M urea in 0.5% SDS in PBS. Proteins were reduced with 10 mM DTT at 65°C for 15 minutes and then alkylated with 20 mM iodoacetamide at 37°C for 30 minutes at 500 rpm. The proteins were re-precipitated with 1 mL ethanol and subjected to the same washing steps as before.

Proteins were resuspended in 500 µL of fresh 2 M urea in PBS, and trypsin was added at a 1:100 enzyme-to-protein (w/w) ratio for overnight digestion at 37°C with shaking at 500 rpm. Following digestion, peptides were transferred to a 15 mL falcon tube containing 10 mL acetonitrile to achieve >95% final volume ACN. Samples were incubated at room temperature with shaking at 1000 rpm for 10 minutes before peptides were collected by centrifugation at 1600 rcf for 5 minutes. The pellet was washed three times with 1 mL ACN, then transferred to a 1.5 mL lo-bind Eppendorf tube. The samples were placed on a magnetic rack and washed once more in 1 mL fresh ACN. After magnetic separation, ACN was removed and peptides were eluted twice using 100 µL of 2% DMSO in water, incubated at 37°C for 30 minutes at 1000 rpm.

For enrichment, 250 µL of high-capacity NeutrAvidin resin (Thermo Fisher, 29204) was washed three times with 10 mL of IAP buffer (50 mM MOPS pH 7.4, 10 mM sodium phosphate, and 50 mM NaCl), then resuspended in a buffer volume adjusted to achieve a final peptide concentration of 0.2 µg/µL. Peptides were incubated with the resin for 2 hours at room temperature on a rotator, then washed twice with PBS and twice with water, with centrifugation at 1600 rcf between each wash. The resin was transferred to a bio-spin column (Bio-Rad, 7326204), washed twice with 500 µL of water, and centrifuged at 800 rcf for 30 seconds.

Enriched peptides were eluted in 60 µL of 80% acetonitrile in water containing 0.1% formic acid, incubated for 10 minutes at room temperature, then centrifuged at 1600 rcf for 1 minute. This elution was repeated at 72°C for 10 minutes at 300 rpm, and a final elution was performed at room temperature, followed by centrifugation at 2600 rcf for 2 minutes. The combined eluates were dried using a SpeedVac.

Crude peptides were then resuspended in 22 µL ACN and sonicated (35 kHz, 10 min). 425 µL water and 2.3 µL trifluoroacetic acid were then added followed by another sonication. C18 resin (Thermo Fisher, 89870) was activated twice with 200 µL of 50% methanol in water, followed by three washes with 5% ACN in water containing 0.1% TFA. Resin was then placed into 2 mL lo-bind tubes for sample loading. Samples were then added onto the resin over three 150 µL additions. After complete loading, 200 µL of flowthrough was re-added to the resin twice to ensure complete binding. Samples were then washed three times as mentioned before. All centrifugation steps were performed at 300 rcf for 15 seconds. Samples were then eluted twice with 20 µL 70% ACN in water containing 0.1% formic acid after a 1 minute incubation (1600 rcf, 1 min). A third elution was performed similarly, but at 2600 rcf for 2 minutes. Eluted, desalted peptides were then dried using a SpeedVac.

Samples were resuspended in 20 µL LC-LOAD (PreOmics) followed by vortexing and sonication (35 kHz, 10 min). Samples were centrifuged (21000 rcf, 10 min) and 15 µL were transferred into MS vials (Thermo Fisher, 200 046) for analysis by mass spectrometry.

### Mass Spectrometry Data Acquisition

A nanoElute2 was attached in line to a timsTOF Pro2 equipped with a CaptiveSpray Source (Bruker, Hamburg, Germany). Chromatography was conducted at 40°C through a 25 cm reversed-phase C18 column (PepSep, 1893476) at a constant flow rate of 0.5 μL min^−1^. Mobile phase A was 98/2/0.1% water/MeCN/formic acid (v/v/v, Thermo Fisher, LS118, LS120) and phase B was MeCN with 0.1% formic acid (v/v). During a 120 min method, peptides were separated by a 3-step linear gradient (4% to 40% B over 95 min, 40% to 60% B over 10 min, 60% to 90% B over 10 min) followed by a 10 min isocratic flush at 95% before washing and a return to low organic conditions. Experiments were run as data-dependent acquisitions with ion mobility activated in PASEF mode. MS and MS/MS spectra were collected with m/z 100 to 1700 and ions with z = +1 were excluded.

### FragPipe Open Search

To comprehensively survey mass shifts present in peptides across the dataset, an Open Search was performed using MSFragger within FragPipe v22.0. The search was configured with the following parameters: precursor mass range −10 to +850 Da, fragment mass tolerance 40 amu, isotope error set to 0, and enzyme set to “trypsin” with cleavage after “KR” and not before “P” (C-terminal specificity). Enzymatic digestion was assumed, allowing up to 2 missed cleavages. Minimum and maximum peptide lengths were set to 7 and 50 residues, respectively, and peptide mass range from 500 to 5000 Da. N-terminal methionine clipping was enabled, and a minimum of 15 peaks and a 0.01 intensity ratio were required per spectrum. Fragment ions considered included *b* and *y* series, with a maximum fragment charge of 2+. Calibration and parameter optimization were disabled, and deisotoping and de-neutralloss filtering were enabled. No fixed modifications were applied, and variable modifications were included but not enforced in the initial search (used only for localization and PTM profiling downstream).PeptideProphet was enabled with the following command-line options: --nonparam --expectscore --decoyprobs --masswidth 1000.0 --clevel - 2, and results from multiple replicates were combined. PTMProphet was disabled. ProteinProphet was run with the setting --maxppmdiff 2000000. PTM-Shepherd was enabled for post-processing and mass shift profiling. Peak picking parameters were set to a smoothing factor of 2 bins, prominence ratio of 0.3, and peak picking width of 0.002 Da. Precursor mass tolerance was 0.01 Da. Fragment ion types considered for modification annotation were *b* and *y*, with a maximum fragment charge of 2+. PTM-Shepherd was run with default UniMod annotations disabled to allow unsupervised delta mass discovery, and glycan annotation was enabled but no glycan database was used. Glycan delta masses were removed before peak picking. Normalization of mass shifts was performed across PSMs (not scans). Report generation was enabled with the following options: --sequential --mapmods --prot 0.01. MS1-based quantification, TMT-Integrator, and Spectral Library generation were disabled. For downstream analysis, the global.modsummary.tsv file from PTM-Shepherd was used to extract the number of PSMs associated with each observed mass shift. These counts were plotted against the theoretical mass shift to visualize the landscape of modifications in the range of +200 to +750 Da.

### FragPipe Closed Search

Mass spectrometry data were analyzed using FragPipe v22.0. Closed Search was performed using MSFragger with the following parameters: precursor mass tolerance −50 to +50 ppm, fragment mass tolerance 50 amu, isotope error set to 0/1/2, enzyme name “stricttrypsin”, cleavage at “KR” but not before “P” (C-terminal cleavage), enzymatic cleavage with up to 2 missed cleavages, peptide length between 7 and 50 residues, and peptide mass range from 500 to 5000 Da. Fragment ion types included *b* and *y* ions, with a maximum fragment charge of 2+. N-terminal methionine clipping was enabled. The number of variable modifications per peptide was limited to 4, and the number of allowed variable modification combinations was capped at 5000. Spectra were required to have a minimum of 15 peaks and a minimum peak intensity ratio of 0.01. Mass calibration was set to “Mass Calibration and Parameter Optimization,” and deisotoping and de-neutralloss processing were enabled.

Variable modifications included oxidation of methionine (+15.9949 Da), N-terminal acetylation (+42.0106 Da), and a pair of arginine labels (+695.2942 Da and +701.3014 Da) each with a maximum of 1 occurrence. Cysteine was either unmodified or labeled with a variable modification of +57.02146 Da (up to 3 occurrences). All other modifications used in the experiment are listed in Supplementary Table 5. Fixed modifications were not applied except in specific runs evaluating cysteine, where carbamidomethylation (+57.02146 Da) was fixed on C.

Crystal-C, PTMProphet, and TMT-Integrator were disabled. PeptideProphet was also disabled. ProteinProphet was enabled and run with the following setting: --maxppmdiff 2000000. Percolator was used for peptide-spectrum match (PSM) validation with the following command-line options: --only-psms --no-terminate --post-processing-tdc, and a minimum probability threshold of 0.5.

MS1-based quantification was enabled using IonQuant with the following parameters: m/z tolerance of 10 ppm, retention time (RT) tolerance of 0.4 min, ion mobility tolerance 0.05 (although ion mobility was not used), labeling mode enabled with two arginine labels (+695.2942 Da and +701.3014 Da), re-quantification enabled, minimum frequency of 0.5, a minimum of 2 isotopes and 1 scan per feature, top 3 ions used for quantification, and no normalization applied. Match-between-runs was disabled. Label-free quantification was not used.

MSBooster, PTMShepherd, SAINTexpress, Skyline, and Spectral Library generation were disabled. Report generation was enabled with the following options: --sequential --razor --mapmods --prot 0.01, with decoys and known contaminants removed. Summary tables were generated at both peptide and protein levels.

### SASA Analysis Pipeline

Output modified peptides from Fragpipe were processed using a custom Python-based pipeline to identify modified residues based on their sequence position in UniProt. Protein identifiers were initially provided as UniProt names, which were converted to their corresponding UniProt accession numbers using the UniProt API (https://rest.uniprot.org/). Structural models were then downloaded from the European Bioinformatics Institute (EBI) AlphaFold Database (https://alphafold.ebi.ac.uk/). Each protein structure was parsed using Bio.PDB (Biopython) and the Shrake-Rupley algorithm^36^ was applied to compute solvent-accessible surface area (SASA) at the atomic level. Residue positions were adjusted by shifting input values down by one to ensure alignment with AlphaFold numbering. To ensure the reliability of structural predictions, residues with a predicted local distance difference test (pLDDT) score below 70 were excluded from the analysis. The relative solvent accessibility (RSA) was then calculated by normalizing SASA values against a reference maximum exposure of 274 Å². Computational storage was optimized by removing downloaded PDB files immediately following analysis, and results were iteratively written to an output file. This automated workflow enabled high-throughput extraction of solvent accessibility data from AlphaFold models, facilitating the structural interpretation of specific residues within predicted protein conformations.

For experimental structures, PDB accessions were retrieved from UniProt cross-references and corresponding mmCIF files were downloaded using Biopython’s MMCIFParser. Structures with ribosomal content or more than 10 chains were excluded. Each PDB structure was aligned to the UniProt sequence to correctly map the modified residue to the structural coordinates. SASA values were computed using the same Shrake-Rupley algorithm, and RSA values were calculated using the same normalization scheme. In cases where multiple structures existed for a given protein, all qualifying entries were processed independently, and the mean and standard deviation of RSA values for each residue were calculated and used in downstream analysis.

PDB Proximal Ligand Search Pipeline To assess the structural context of modified arginine residues in proteins, an in-house computational pipeline was developed to map modified arginines from the Fragpipe output onto full-length UniProt sequences, align them to resolved PDB structures, and evaluate their spatial proximity to bound ligands. The input dataset consisted of a CSV file containing UniProt accession numbers and peptide sequences with modified arginine residues denoted as R[701.3014]. Other modifications annotated in brackets, such as oxidation or carbamidomethylation, were ignored. The modified residue’s position within the peptide was extracted, and all bracketed modifications were removed to generate a cleaned peptide sequence for further processing. To determine the position of the modified arginine in the full-length protein sequence, the cleaned peptide was aligned to the UniProt sequence using Biopython’s PairwiseAligner with the BLOSUM62 substitution matrix, and affine gap penalties (−10 for opening, −0.5 for extension). The modified residue’s position in the peptide was mapped to its corresponding position in the aligned UniProt sequence, yielding the residue number in the full-length protein. The script then queried the UniProt cross-references to retrieve all associated PDB accession numbers for each protein. Structures were downloaded in the PDBx/mmCIF format using MMCIFParser from Biopython. To improve computational efficiency and ensure relevance to the analysis, structures containing ribosomal components were excluded, and entries with more than 10 chains were filtered out. For structural mapping, the UniProt sequence was aligned to the sequences of all polypeptide chains in each PDB structure, allowing for potential discrepancies in residue numbering between sequence databases and structural models. The modified residue’s position in the PDB structure was identified based on this alignment, ensuring that subsequent analyses were performed on the correct structural residue. To assess the molecular environment of modified arginines, the spatial relationship between the CZ atom of the modified residue and bound ligands was analyzed. NeighborSearch from Biopython^59^ was used to determine whether any ligand atoms were within 6.0 Å of the arginine CZ atom. Water molecules (residue name ‘HOH’) were excluded to focus the analysis on small molecules and cofactors. If a ligand met the distance threshold, its residue name and chain position were recorded. To ensure robustness in handling large datasets, results were written incrementally to a CSV file containing the UniProt ID, PDB ID, chain identifier, mapped residue position in the PDB, and the nearest ligand within the specified distance.

### AlphaMissense Pathogenicity Annotation

To evaluate the potential functional relevance of modified arginines, AlphaMissense pathogenicity scores were incorporated into the dataset. UniProt accession numbers and residue positions for all modified arginines were used to generate a queryable index. A custom Python script was developed to query the AlphaMissense API (https://alphamissense.hegelab.org/hotspotapi) for each protein and residue pair using concurrent requests with up to 10 worker threads to accelerate the process. Isoform-specific identifiers (e.g., P12345-2) were excluded to ensure consistency with canonical AlphaMissense annotations. For each query, the API returned a predicted mean pathogenicity score and a classification (e.g., ‘likely pathogenic’, ‘likely benign’). If a classification was not explicitly returned, a custom threshold-based scheme was applied: scores ≤ 0.34 were labeled ‘likely benign’, scores ≥ 0.564 were labeled ‘likely pathogenic’, and intermediate values were designated ‘ambiguous’. All results were written to a CSV file including the protein ID, residue position, score, and classification.

### Secondary Structure Assignment via DSSP

To assess local structural context of modified residues, DSSP-based secondary structure annotations were computed for both AlphaFold-predicted and experimentally resolved PDB structures. For AlphaFold models, UniProt IDs were first mapped to primary accessions using the UniProt API. Structural models were downloaded from the AlphaFold Protein Structure Database. For each protein, the corresponding PDB file was parsed using Biopython’s PDBParser, and DSSP secondary structure assignments were computed using Bio.PDB.DSSP. Modified residue positions were matched to AlphaFold residue indices (adjusted for 0-based numbering), and residues with a predicted local distance difference test (pLDDT) score below 75 were excluded. For residues passing the pLDDT filter, a ±4 residue sliding window was used to collect local secondary structure context. Both the DSSP code for the modified residue and its surrounding window were recorded.

For experimental structures, UniProt cross-references were used to identify relevant PDB accessions. Structures were downloaded in mmCIF format and parsed using Biopython’s MMCIFParser. Resolution and structural complexity filters were applied: only structures with ≤10 chains and ≤3.0 Å resolution were retained. The best-aligned chain was selected by global pairwise sequence alignment using Biopython’s PairwiseAligner, and modified residues were mapped from UniProt positions to structure-specific indices. DSSP was run on each selected PDB model to obtain secondary structure codes. For each modified residue, the DSSP code and ±3 residue sequence context were recorded. Results were aggregated across all qualifying structures for each protein, and the most frequently observed DSSP code (majority vote) was assigned as the representative secondary structure for that residue. Confidence scores were calculated as the frequency of the major assignment relative to total observations.

### Motif Enrichment Analysis via pLogo

To investigate sequence preferences surrounding modified arginines, motif enrichment analysis was performed using pLogo (https://plogo.uconn.edu). Foreground sequences were generated from a list of experimentally identified arginine modification sites, with UniProt accession numbers and residue positions extracted from FragPipe output. A custom Python script queried the UniProt API to retrieve the full protein sequences and extracted a 21-residue window (±10 amino acids) centered on each modified arginine. If a full window could not be extracted due to proximity to sequence termini, the missing positions were padded with ‘X’ characters to maintain fixed length. Background sequences were generated separately from the full UniProt human proteome FASTA (including isoforms). For each protein sequence, a sliding window approach was used to extract 21-residue sequences centered on every amino acid position, again padding with ‘X’ as needed. This resulted in an unfiltered and comprehensive background set. Due to file size limitations on the pLogo web interface, the background set was downsampled to 1,000,000 sequences using Python’s random.sample() method. Foreground and background sets were submitted to the pLogo web interface, specifying arginine (R) as the central anchor residue. A binomial statistical model with Bonferroni correction was used to evaluate motif significance. Enrichment scores and residue logos were downloaded and replotted using custom scripts to standardize styling across figures.

### Salt Bridge Identification in AlphaFold Structures

To identify potential salt bridge interactions between modified arginines and acidic residues, AlphaFold-predicted structural models were analyzed using a custom Python pipeline. UniProt IDs and modified residue positions were extracted from a CSV input file. Isoform identifiers were excluded. For each canonical UniProt ID, the AlphaFold API was queried to retrieve the corresponding structure metadata and download the associated PDB file. Models were stored locally and parsed using Biopython’s PDBParser. Each arginine residue was scanned for proximity to nearby aspartate (ASP) or glutamate (GLU) side chain atoms (OD1/OD2 or OE1/OE2, respectively). Only models with pLDDT scores ≥ 70 for both arginine and acidic residues were considered. Salt bridges were defined by the presence of at least one NE, NH1, or NH2 atom from the modified arginine within 4.0 Å of the target carboxylate atom. When multiple atoms satisfied the distance criterion, the closest atom pair was selected. Electrostatic interaction energies were estimated using a simplified Coulombic potential:, assuming unit charges, a dielectric constant of 20.0, and interatomic distance in Å. For each interaction, the chain ID, residue names and numbers, interacting atoms, interatomic distance, pLDDT scores, and computed Coulombic energy were recorded. Following analysis, downloaded PDB files were removed to conserve disk space. Final results were compiled into a CSV file for downstream analysis and visualization.

### Cation–π Interaction Identification in AlphaFold Structures

To identify potential cation–π interactions involving modified arginines, AlphaFold-predicted structures were evaluated using a geometry-based Python workflow. For each UniProt ID and arginine position, the corresponding AlphaFold model was downloaded from the EBI database. PDB files were parsed with Biopython’s PDBParser, and residues were scanned for interactions between arginine CZ atoms and aromatic residues (PHE, TYR, TRP, HIS). For each aromatic residue, centroid coordinates were computed from annotated ring atoms (e.g., CZ, CE1, CE2, etc.), and the Euclidean distance to the arginine CZ atom was calculated. Interactions within 6.0 Å were considered candidates. If angular filtering was enabled, the orientation between the CZ–centroid vector and the ring plane normal was used to classify geometry as parallel (face-to-face) or T-shaped (edge-to-face), with thresholds of ≤30° and 90°±20°, respectively. Models with pLDDT scores below 70 for either the arginine or aromatic residue were excluded. For each identified interaction, the protein ID, chain IDs, residue positions and types, distance, angle, geometry classification, and pLDDT scores were recorded. Only one interaction was logged per arginine to avoid multiple counting. Results were written to two CSV outputs: a detailed interaction table and a simplified summary indicating whether each modified arginine participates in a cation–π interaction.

### Salt Bridge and Cation–π Interaction Analysis in Experimental PDB Structures

To validate and expand upon AlphaFold-based interaction predictions, salt bridges and cation–π interactions were also identified using experimentally resolved PDB structures. For each UniProt ID and modified residue position, all associated PDB structures were retrieved via UniProt cross-references. Structures were excluded if they represented ribosomal proteins, large macromolecular complexes (taxonomy count > 15), NMR structures, or models containing more than 10 chains. PDBx/mmCIF files were downloaded and parsed using Biopython’s MMCIFParser. For each candidate arginine, neighboring residues were identified using NeighborSearch, and interactions were classified as salt bridges or cation–π based on geometric and electrostatic criteria. Salt bridges were defined as any acidic residue (ASP, GLU) within 4.0 Å of NE, NH1, or NH2 atoms of the arginine. Cation–π interactions were defined as cases where the arginine CZ atom was within 8.0 Å of the centroid of an aromatic side chain (PHE, TYR, TRP, HIS). When enabled, angular filtering was used to categorize interaction geometries as face-to-face (≤30°) or T-shaped (90° ± 20°). All interactions were annotated with residue identity, chain location, distance, geometry (if applicable), whether the partner residue was part of the same polypeptide chain, and optionally, an estimated electrostatic energy. Final results were compiled into a single CSV table containing all identified intermolecular interactions from experimental PDB models.

### pKa Estimation for Arginines Using Experimental PDB Structures

To estimate side-chain protonation states of modified arginines, PROPKA 3.0 was used to calculate predicted pKa values from experimental structures in the Protein Data Bank (PDB). Modified arginines were identified by UniProt accession and residue position. Isoforms and ribosomal proteins were excluded. Associated PDB structures were retrieved via UniProt cross-references and filtered to exclude NMR structures, entries with >10 polymer entities, or structures with >10 chains. Structures were downloaded in mmCIF format and parsed using Biopython’s MMCIFParser. Each structure was aligned to the canonical UniProt sequence using PairwiseAligner to identify the best-matching chain and map UniProt residue indices to structural coordinates. Only arginines with a resolved CZ atom and a B-factor ≥ 30.0 were considered. Structures were converted to PDB format using Biopython’s PDBIO to ensure compatibility with PROPKA. The PROPKA executable was run via subprocess call, and predicted pKa values were parsed from the resulting .pka files. For each modified arginine, the output included UniProt ID, PDB ID, chain ID, UniProt and author residue numbers, and pKa value. Structures and temporary output files were deleted after use to conserve disk space.

### pKa Estimation for Arginine, Lysine or Cysteine from AlphaFold Models

To complement structure-based pKa estimation, PROPKA was also applied to AlphaFold-predicted structures. AlphaFold models were downloaded in PDB format for each UniProt accession. Only residues with pLDDT confidence scores >70 were considered for analysis. For each AlphaFold structure, PROPKA 3.0 was executed using the PDB file as input. Predicted pKa values were extracted using a custom parser that accommodated multiple possible output formats. Residue-level pKa values were reported for each modified arginine, cysteine or lysine, alongside the corresponding pLDDT score. Proteins for which PROPKA failed to produce valid output were excluded. Temporary AlphaFold PDB and PROPKA output files were deleted after processing.

### Intrinsic Disorder Prediction via IUPred2A

To evaluate the intrinsic disorder propensity of modified arginine residues, disorder scores were retrieved using the IUPred2A web API (https://iupred2a.elte.hu). A CSV file containing UniProt IDs and modified residue positions served as input. For each protein, the full-length disorder profile was fetched in JSON format and parsed to extract per-residue IUPred2 scores. Modified arginine positions were indexed using 1-based UniProt residue numbering. Prior to extraction, each index was converted to 0-based to match Python list conventions. If a position fell within the length of the retrieved disorder profile, the corresponding IUPred2 score was recorded. Otherwise, ‘NA’ was assigned. Each entry in the output file included the UniProt ID, residue position, and corresponding disorder score. Results were written iteratively to a CSV file for downstream integration with structural and reactivity data.

### Molecular Docking and Covalent Adduct Modeling

Molecular docking was performed using the Molecular Operating Environment (MOE2024). The crystal structures of aconitase (PDB: 1B0J) and cyclophilin A (PDB: 6GJI) were obtained, with all crystallographic ions removed. Each structure was solvated in a 10 Å water sphere to mimic physiological conditions before preparation using MOE’s QuickPrep feature, which optimized hydrogen placement, assigned protonation states, and performed an initial energy minimization. For aconitase, the docking site was defined based on the known binding site of isocitrate, and the dehydrate of ninhydrin was docked using MOE’s docking protocol. For cyclophilin A, the ligand CA-Nin-Alk was docked using MOE’s template-based docking to ensure proper positioning. In both cases, the top-scoring pose was selected and loaded into the MOE Window, where a covalent adduct was modeled. Energy minimization was performed using the Amber10:EHT force field with gradient-based optimization until the root mean square gradient reached 0.05 kcal/(mol·Å). Visualization and figure preparation were performed using ChimeraX 1.6.1^60^.

### Cyclophilin A Inhibition Assay

To measure CA-Nin-Alk inhibition of cyclophilin A (CypA) activity, a peptidyl-proline isomerase (PPIase) assay was utilized following a previously established protocol^54^. Recombinant human CypA (R&D Systems) was diluted to a concentration of 1 µM in a reaction buffer of 20 mM HEPES, 100 mM NaCl, and 0.5 mM TCEP (pH 7.5) and kept on ice. Chymotrypsin was diluted to 60 mg/mL in 2 mM CaCl_2_, 1 mM HCl in H_2_O. To test inhibition, the inhibitor, CA-Nin-Alk, was diluted in DMSO to generate 100X stock solutions (25 µM, 10 µM). These were added to CypA solutions (final DMSO = 1% v/v) and incubated for 1 hour at room temperature. The substrate, suc-AAPF-pNA (Thomas Scientific), was freshly prepared at 10 mM in 470 mM LiCl/TFE. The spectrophotometer (Cary UV-Vis Spectrophotometer) was cooled to 4°C before use.

For uncatalyzed reactions, 980 µL of reaction buffer and 10 µL of chymotrypsin were mixed in a cuvette and placed in the spectrophotometer. The reaction was initiated with 10 µL substrate and A390 was monitored every 0.1s for 5 minutes. For CypA catalyzed reactions, 970 µL of the reaction buffer, 10 µL of chymotrypsin, and 10 µL of CypA solution (pre-incubated ± inhibitor) were combined prior to substrate addition and identical data collection. All assays were performed in triplicate. To ensure calculations depended solely on the first-order reaction of isomerization and not residual trans-substrate in the solution, the absorbance traces, excluding the first 10s, were fit to a single-exponential function to obtain the rate constants for uncatalyzed (*k_uncat_*) or CypA-catalyzed (*k_obs_*) isomerization. *k_cat_/K_m_* values were then calculated via the following equation:

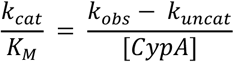

using a final enzyme concentration of 1*10^−8^M.

### IC_50_ calculations

Apparent IC_50_ values were estimated using a two-point linear interpolation. Percent inhibition of CypA catalytic efficiency (*k_cat_/K_m_*) was measured at two compound concentrations (25 µM, 10 µM). A linear regression of inhibition (*y*) vs compound concentration (*x*) was used to approximate the relationship (*y = mx + b*), and the concentration corresponding to 50% inhibition was calculated as:

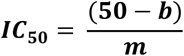

### Code Availability Statement

All code is available on the ZaroLab GitHub. Code is available for academic use, all other non-academic usage is restricted to those with permission from the corresponding author.

### Statement of Data Availability

All raw mass spectrometry datasets are available on MASSive under the ID: MSV000097646. Other raw data and processed data essential to this work are provided in the Supplementary Information and Source Data file. Source data are provided with this paper via 10.5281/zenodo.15579687.

## Uncropped Gels

Uncropped gel images from Figures 2B and Extended Data Figure 2A.

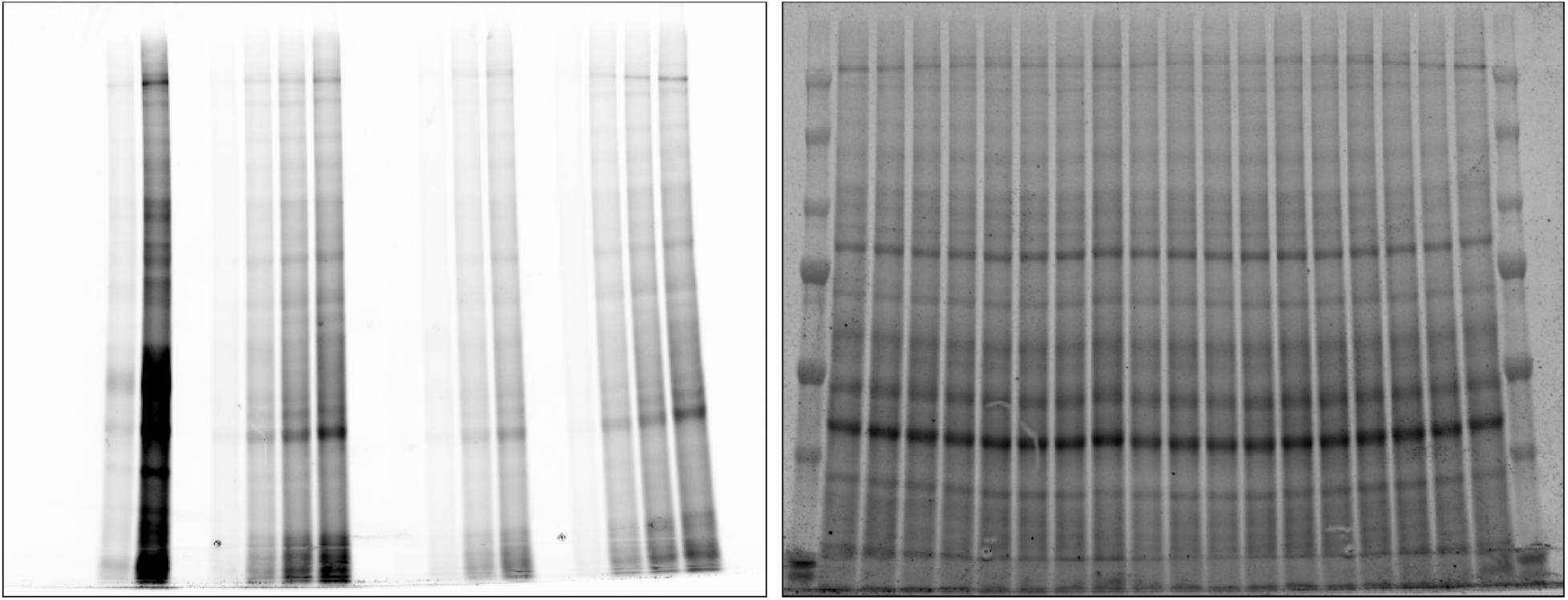

Qualitative, gel-based assessment of proteome-wide reactivity of alkyne-bearing ninhydrin probes in Mino cell lysate. Soluble proteome (1.5 µg µL⁻¹) was treated with each probe at increasing concentrations (1, 10, or 100 µM) for 1 h, followed by CuAAC conjugation with TAMRA-N₃, separation by SDS–PAGE, and in-gel fluorescence scanning.

Uncropped gel images from Figure 2C.

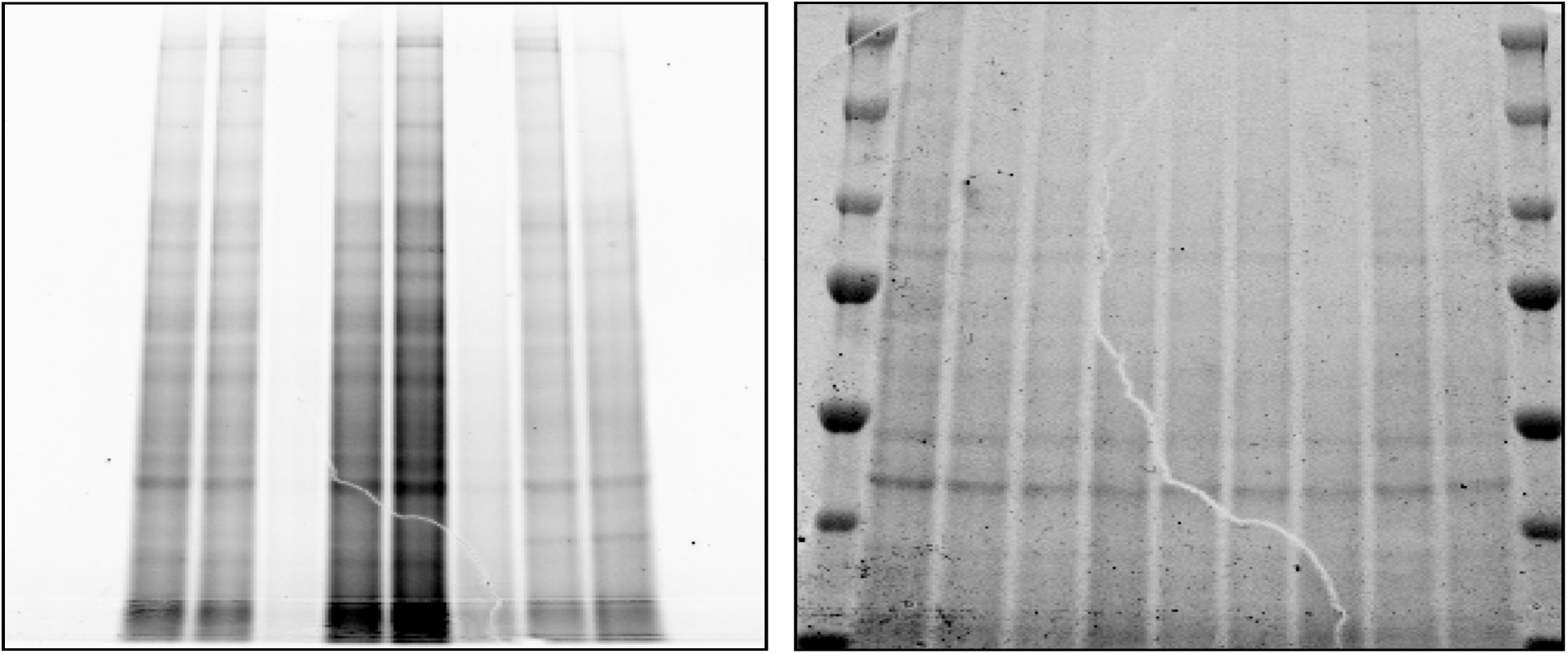

Qualitative time-dependent reactivity of 4-Nin-Alk, Nin-Alk, and PGO in live cells. Mino cells were treated with 100 µM of indicated probe for 0, 30, or 60 min followed by lysis, CuAAC conjugation with TAMRA-N₃, separation by SDS–PAGE, and in-gel fluorescence scanning.

### CypA Kinetics Data

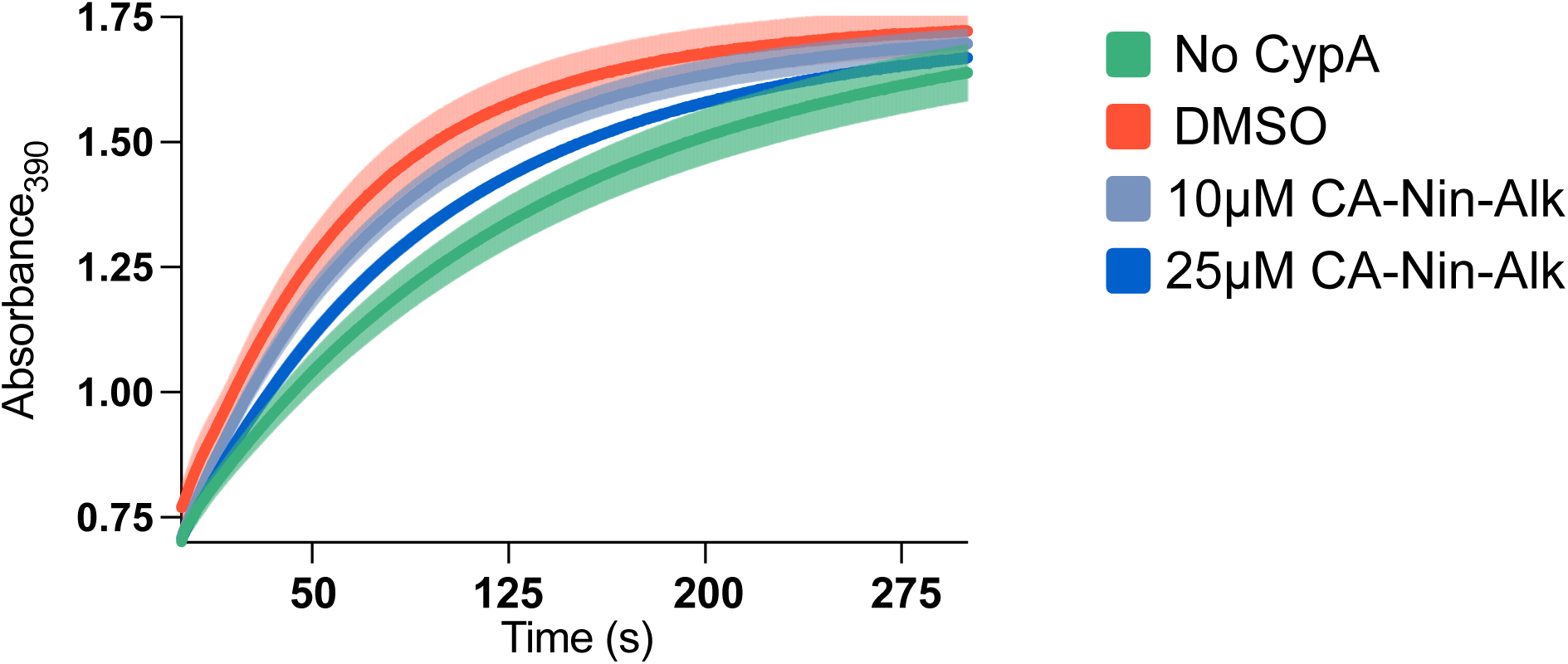

Kinetics data for the Cyclophilin A (CypA) PPIase assay in Figure 6G. Absorbance was measured for the chromogenic substrate suc-AAPF-pNA in the presence or absence of CypA and our inhibitor CA-Nin-Alk at indicated concentrations. Experimental details are described in the methods section. Individual measurements are included in Table 16. Each datapoint on the graph is the average of the 3 replicates and the error bars represent the standard error to the mean.

### Representative Traces

Representative Fmoc-AA-OH consumption assay HPLC traces.

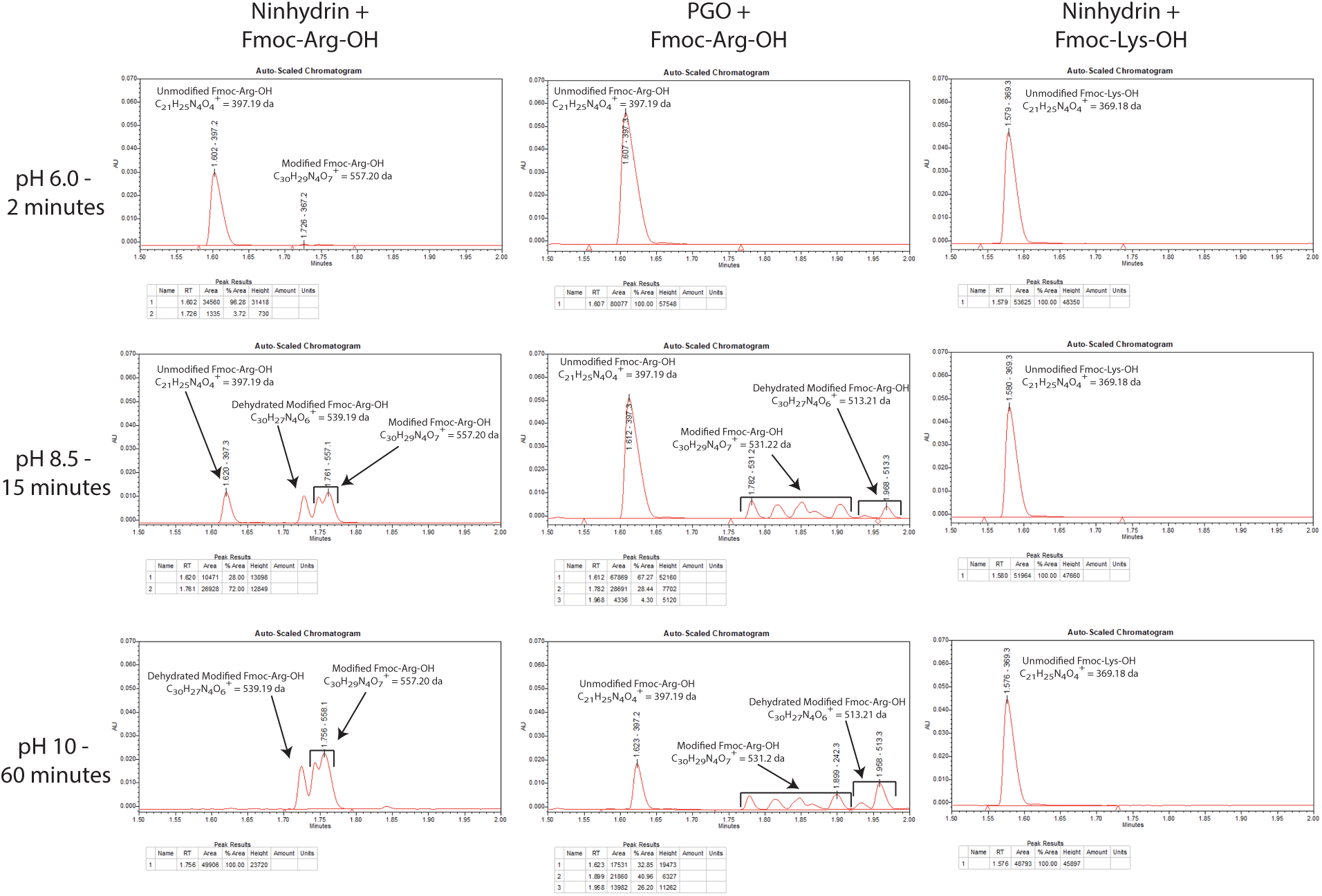

Representative HPLC traces for Fmoc-AA-OH consumption assay in Figure 1B, Figure 3C, Extended Data Figure 1A and Extended Data 3A. Kinetic analysis of pH- and time-dependent consumption of Fmoc-Arg or Fmoc-Lys (500 µM) by ninhydrin or PGO (15 mM, 30x) in PBS at 20 °C. Fmoc-AA consumption was quantified by LC-MS HPLC traces at 300nm wavelength for samples quenched between 2 to 60 minutes. Error bars represent SD. Each trace represents an individual replicate.

Representative Intact Protein Mass Spectrometry Deconvoluted Spectra.

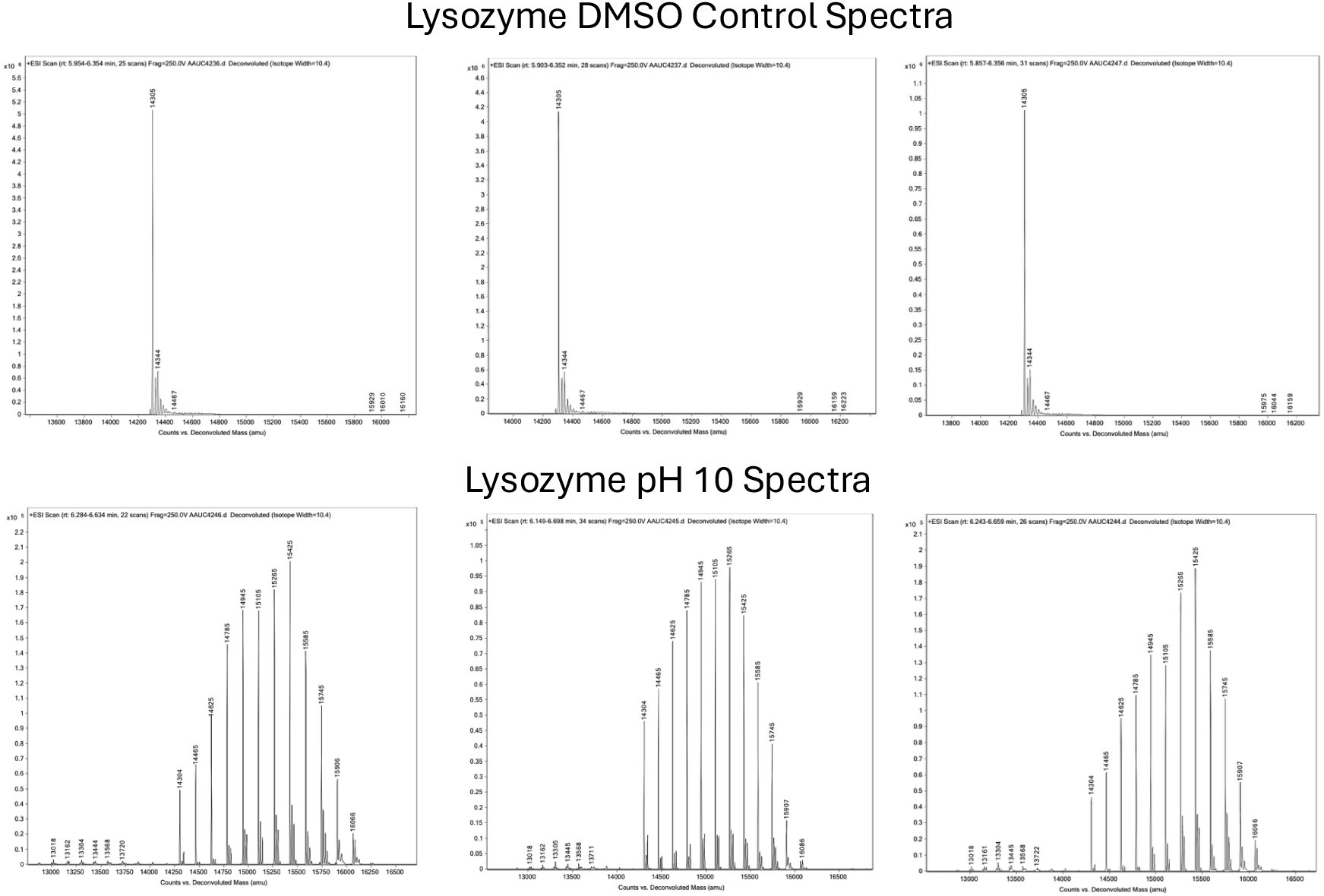

Representative intact protein mass spectrometry deconvoluted mass spectra as acquired for Figures 1D and Extended Data Figure 3. Intact protein mass spectrometry analysis of Lysozyme (1.5 µM) and 100 µM ninhydrin at 37 °C for 30 minutes across pH 4.0, 7.0 and 10.0. Reactions were quenched by dilution into 0.1% aqueous TFA and analyzed by LC–MS on an Agilent 6230 TOF system. Each spectra represents an individual replicate

### Synthesis of Probes

**General Experimental Procedures:** All reactions were performed in flame- or oven-dried glassware fitted with rubber septa under a positive pressure of nitrogen or argon, unless otherwise noted. All reaction mixtures were stirred throughout the course of each procedure using Teflon-coated magnetic stir bars. Air- and moisture-sensitive liquids were transferred via syringe or stainless-steel cannula. Solutions were concentrated by rotary evaporation below 35 °C. Analytical thin-layer chromatography (TLC) was performed using glass plates pre-coated with silica gel (0.25-mm, 60-Å pore size, 230−400 mesh, SILICYCLE INC) impregnated with a fluorescent indicator (254 nm). TLC plates were visualized by exposure to ultraviolet light (UV), and then were stained by submersion in a basic aqueous solution of potassium permanganate or with an acidic ethanolic solution of ninhydrin, followed by brief heating.

**Materials:** Dichloromethane (DCM), dimethylformamide (DMF), tetrahydrofuran (THF), ethyl ether, and acetonitrile to be used in anhydrous reaction mixtures were dried by passage through activated alumina columns immediately prior to use. Hexanes used were ≥85% n-hexane. Other commercial solvents and reagents were used as received, unless otherwise noted.

**Instrumentation:** Unless otherwise noted, proton nuclear magnetic resonance (^1^H NMR) spectra and carbon nuclear magnetic resonance (^13^C NMR) spectra were recorded on a JEOL JNM-ECZ400R 400 MHz NMR equipped with 5 mm H/F/X Royal Probe or a Bruker AVIII HD 600 MHz NMR equipped with 5 mm CPDCH CryoProbe at 23 °C. Proton chemical shifts are expressed in parts per million (ppm, δ scale) and are referenced to residual protium in the NMR solvent (CDCl_3_: δ 7.26, CD_3_CN: δ 1.94). Carbon chemical shifts are expressed in parts per million (ppm, δ scale) and are referenced to the carbon resonance of the NMR solvent (CDCl_3_: δ 77.16, CD_3_CN: δ 118.26). Data are represented as follows: chemical shift, multiplicity (s = singlet, d = doublet, t = triplet, q = quartet, dd = doublet of doublets, dt = doublet of triplets, sxt = sextet, m = multiplet, br = broad, app = apparent), integration, and coupling constant (J) in hertz (Hz). Mass measurements for high-resolution mass spec (HRMS) were performed on a Waters Xevo G2-XS TOF calibrated against sodium formate clusters and using a LeuEnk lockmass. Expected monoisotopic masses were calculated using MassLynx 4.1 and the *m/z* values for calibrant and lockmass were MassLynx-default values.

### Compound SI-1

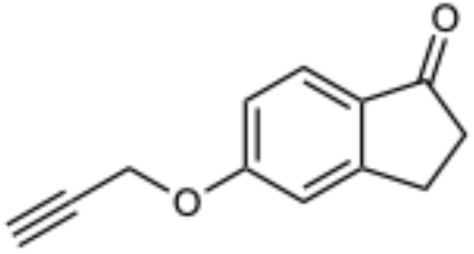

DMF (13.5 mL) was added to a round-bottom flask containing K₂CO₃ (1.87 g, 13.50 mmol, 2.00 equiv) and 5-hydroxy-2,3-dihydro-1H-inden-1-one (1.00 g, 6.75 mmol, 1 equivl). Propargyl bromide (80 wt% in toluene, 829 µL, 1.10 g, 7.42 mmol, 1.10 equiv) was added dropwise over 1 minute. The reaction vessel was heated in a 60 °C oil bath for 16 h. After cooling to ambient temperature, the mixture was diluted with DCM (30 mL) and washed with DI water (3 × 30 mL) followed by brine (2 × 30 mL). The organic layer was dried over Na₂SO₄, filtered, and the filtrate was concentrated under reduced pressure to afford **SI-1** as a brown solid, which was used in subsequent steps without further purification (1.22 g, 97% yield).

**TLC** (hexanes:ethyl acetate = 8:2): R*_f_* = 0.25 (UV)

**¹H NMR** (400 MHz, CDCl₃) δ: 7.71 (d, *J* = 8.4 Hz, 1H), 7.00 (s, 1H), 6.97 (d, *J* = 8.7 Hz, 1H), 4.77 (t, *J* = 2.1 Hz, 2H), 3.11 (t, *J* = 5.4 Hz, 2H), 2.69 (t, *J* = 5.9 Hz, 2H), 2.57 (s, 1H).

**¹³C NMR** (101 MHz, CDCl₃) δ: 205.41, 163.06, 158.06, 131.22, 125.50, 115.88, 111.00, 77.74, 76.35, 56.05, 36.53, 25.99.

**HRMS (ESI⁺):** *m/z* calcd for C₁₂H₁₁O₂⁺ ([M+H]⁺): 187.0759; found: 187.0756.

### Nin-Alk

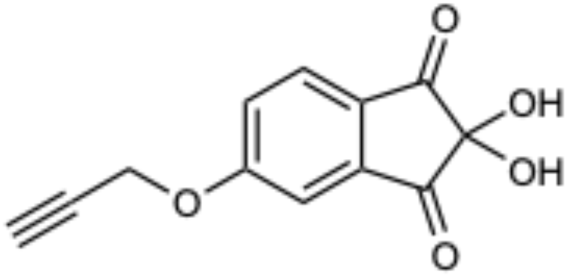

Selenium dioxide (745 mg, 6.71 mmol, 2.5 equiv) was added to a solution of **SI-1** (500 mg, 2.69 mmol, 1 equiv) in 1,4-dioxane (3.25 mL) and DI water (325 μL). The reaction mixture was heated to reflux in a 96 °C oil bath for 4 hours. After cooling to ambient temperature, the mixture was diluted with ethyl acetate (15 mL) and washed successively with DI water (2 × 10 mL), 1 M aqueous HCl (2 × 10 mL). The organic layer was washed with brine (1 × 10 mL), dried over Na₂SO₄, filtered, and the filtrate was concentrated under reduced pressure. The crude residue was purified by flash chromatography (30–50% acetone in hexanes) to afford a red solid which was still contaminated with what may be some selenium byproducts. Final purification was performed by reverse-phase HPLC (2–25% MeCN with 0.1% TFA in milliQ water with 0.1% TFA), and lyophilization of the collected fractions yielded the product as a white powder (561 mg, 90% yield).

**TLC** (hexanes:acetone = 1:1): R*_f_* = 0.34 (UV)

**¹H NMR** (400 MHz, 10:1 CD_3_CN + D_2_O) δ: 7.91 (d, J = 8.6 Hz, 1H), 7.47 (dd, J = 8.6, 2.5 Hz, 1H), 7.40 (d, J = 2.4 Hz, 1H), 4.88 (d, J = 2.4 Hz, 2H), 2.95 (t, J = 2.4 Hz, 1H)

**¹³C NMR** (151 MHz, 10:1 CD_3_CN + D_2_O) δ: 196.80, 195.28, 165.07, 142.06, 133.47, 126.88, 126.43, 107.55, 88.21, 77.88, 77.73, 57.22

**HRMS (ESI⁺):** *m/z* calcd for C₁₂H₉O₅⁺ ([M+H]⁺): 233.0450; found: 233.0448.

### Compound SI-2

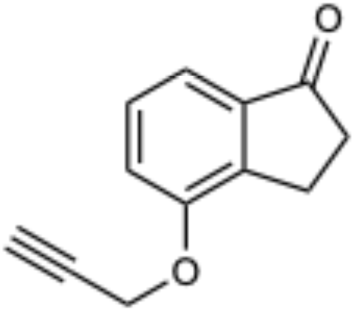

To a round-bottom flask containing K₂CO₃ (1.87 g, 13.50 mmol, 2.00 equiv) and 4-hydroxy-2,3-dihydro-1H-inden-1-one (1.00 g, 6.75 mmol, 1 equiv) was added anhydrous DMF (13.5 mL). Propargyl bromide (80 wt% in toluene, 829 µL, 1.10 g, 7.42 mmol, 1.10 equiv) was added dropwise over the course of 1 minute. The vessel was heated in a 60 °C oil bath for 16 h. After cooling to ambient temperature, the mixture was diluted with DCM (30 mL) and washed with DI water (3 × 30 mL) followed by brine (2 × 30 mL). The organic layer was dried over Na₂SO₄, filtered, and the filtrate was concentrated under reduced pressure to afford **SI-2** as a tan solid, which was used in subsequent steps without further purification (1.18 g, 94% yield).

**TLC** (hexanes:ethyl acetate = 8:2): R*_f_* = 0.40 (UV)

**¹H NMR** (400 MHz, CDCl₃) δ: 7.40 (d, *J* = 7.6 Hz, 1H), 7.36 (t, *J* = 7.6 Hz, 1H), 7.17 (d, *J* = 7.7 Hz, 1H), 4.80 (s, 2H), 3.07 (t, *J* = 6.0 Hz, 2H), 2.69 (t, *J* = 6.0 Hz, 2H), 2.55 (s, 1H)

**¹³C NMR** (100 MHz, CDCl₃) δ: 207.23, 155.10, 144.57, 139.04, 128.86, 116.50, 116.45, 78.18, 76.22, 56.04, 36.26, 22.70.

**HRMS (ESI⁺):** *m/z* calcd for C₁₂H₁₁O₂⁺ ([M+H]⁺): 187.0759; found: 187.0757

### 4-Nin-Alk

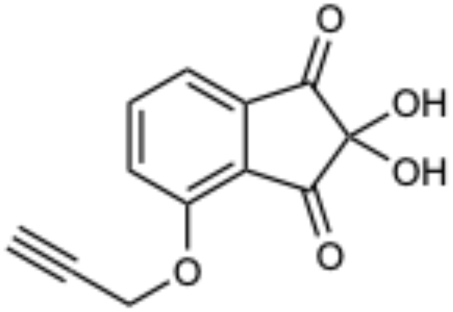

To a solution of **SI-2** (500 mg, 2.69 mmol, 1 equiv) in 1,4-dioxane (3.25 mL) and DI water (325 μL) was added selenium dioxide (745 mg, 6.71 mmol, 2.50 equiv). The reaction mixture was heated to reflux in a 96 °C oil bath for 4 hours. After cooling to ambient temperature, the mixture was diluted with ethyl acetate (15 mL) and washed successively with DI water (2 × 10 mL) and 1 M aqueous HCl (2 × 10 mL). The organic layer was washed with brine (1 × 10 mL), dried over Na₂SO₄, filtered, and the filtrate was concentrated under reduced pressure. The crude residue was purified by flash chromatography (30–50% acetone in hexanes) to yield a red solid. Final purification was performed by reverse-phase HPLC (2–25% MeCN with 0.1% TFA in milliQ water with 0.1% TFA) to remove selenic acid byproducts yielding a white powder after lyophilization (532 mg, 2.29 mmol, 85% yield).

**TLC** (hexanes:acetone = 1:1): R*_f_* = 0.36 (UV)

**¹H NMR** (400 MHz, 10:1 CD_3_CN + D_2_O) δ: 7.93 (dd, J = 8.3, 7.6 Hz, 1H), 7.58 (d, J = 7.0 Hz, 1H), 7.55 (d, J = 7.9 Hz, 1H), 4.96 (d, J = 2.4 Hz, 2H), 2.97 (t, J = 2.4 Hz, 1H)

**¹³C NMR** (100 MHz, 10:1 CD_3_CN + D_2_O) δ: 196.78, 194.41, 156.65, 141.17, 139.49, 127.15, 121.36, 116.92, 87.68, 78.00, 77.95, 57.26

**HRMS (ESI⁺):** *m/z* calcd for C₁₂H₉O₅⁺ ([M+H]⁺): 233.0450; found: 233.0446

### Compound SI-3

A solution of 4-(trifluoromethyl)benzaldehyde (1.74 g, 10.0 mmol, 1 equiv), propargylamine (826 mg, 15.0 mmol, 2.50 equiv), sodium triacetoxyborohydride (5.30 g, 25.0 mmol, 2.50 equiv), and glacial acetic acid (600 mg, 10.0 mmol, 1 equiv) in DCM (95 mL) was stirred at ambient temperature for 2.5 h. The reaction mixture was diluted with DCM (40 mL) and saturated aqueous NaHCO₃ (40 mL) was added, and the layers were mixed vigorously and then separated. The aqueous layer was extracted with DCM (2 × 20 mL), and the combined organic layers were concentrated under reduced pressure. The residue was dissolved in ether (50 mL) and extracted with 1.0 M aqueous HCl (3 × 30 mL). The combined aqueous extracts were basified to pH ∼11–12 (as judged by pH strips) with 1.0 M aqueous NaOH and extracted with ether (4 × 20 mL). The combined organic layers were dried over Na₂SO₄, filtered, and the filtrate was concentrated. The crude residue was purified by flash chromatography (silica gel, gradient from 10-25% ethyl acetate in hexanes) afforded N-(4-(trifluoromethyl)benzyl)prop-2-yn-1-amine (2.05 g, 96% yield) as a colorless oil.

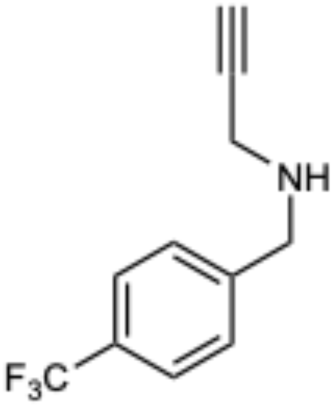

**TLC** (hexanes:ethyl acetate = 4:1): R*_f_* = 0.32 (ninhydrin)

**^1^H NMR** (400 MHz, CDCl₃) δ: 7.59 (d, J = 7.9 Hz, 2H), 7.48 (d, J = 7.9 Hz, 2H), 3.95 (s, 2H), 3.43 (s, 2H), 2.27 (s, 1H)

**^13^C NMR** (101 MHz, CDCl₃) δ: 143.58(d, J = 32.57), 129.73, 128.67, 125.42(q, J= 3.8 Hz), 124.23 (q, J = 271 Hz) 81.78, 71.94, 51.68, 37.41

**^19^F NMR** (376 MHz, CDCl_3_) δ: −62.31

**HRMS (ESI⁺):** *m/z* calcd for C_11_H_11_F_3_N⁺ ([M+H]⁺): 214.0884; found: 214.0842

### Compound SI-4

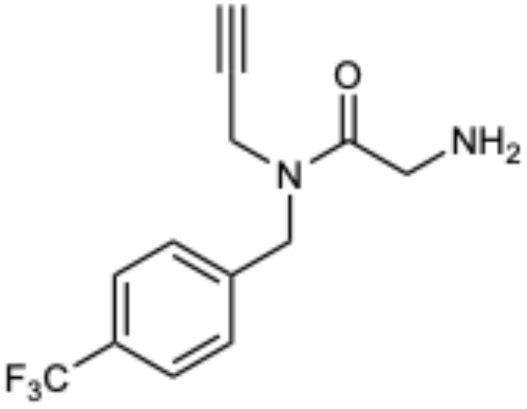

To a solution of **SI-3** (365 mg, 1.71 mmol, 1 equiv) in DMF (8.56 mL) was added DIPEA (664 mg, 895 µL, 5.14 mmol, 3.00 equiv), followed by (tert-butoxycarbonyl)glycine (330 mg, 1.88 mmol, 1.10 equiv) and PyAOP (976 mg, 2.57 mmol, 1.50 equiv). The mixture was briefly sonicated and stirred at ambient temperature for 1 hour. The reaction was diluted with ethyl acetate (50 mL) and sequentially washed with 1.0 M aqueous HCl (2 × 15 mL), DI water (2 × 15 mL), saturated aqueous NaHCO₃ (2 × 15 mL), and brine (2 × 15 mL). The organic layer was dried over Na₂SO₄, filtered, and the filtrate was concentrated under reduced pressure to afford the Boc-protected intermediate as a white solid. This material was then dissolved in 4 M aqueous HCl in dioxane and stirred for 3 hours at ambient temperature to yield the HCl salt of the product as a white solid after diluting in toluene and concentrating under reduced pressure. The HCl salt was freebased with 2 mL aqueous saturated NaHCO₃ and extracted with 20 mL ethyl acetate which was subsequently dried over Na₂SO₄, filtered, and the filtrate was concentrated under reduced pressure to yield pure **SI-4** (392.7 mg, 85% yield) as a white foam which was used in subsequent steps without further purification.

**TLC** (hexanes:ethyl acetate = 1:1): R*_f_* = 0.35 (ninhydrin)

**¹H NMR** (400 MHz, CDCl₃) δ: 7.54 (d, *J* = 8.0 Hz, 1H), 7.49 (d, *J* = 8.0 Hz, 1H), 7.42 (d, *J* = 8.0 Hz, 1H), 7.33 (d, *J* = 8.0 Hz, 1H), 4.60 (s, 2H, Ha, major rotamer), 4.62 (s, 2H, Ha′, minor rotamer), 4.29 (s, 2H, Hb, major rotamer), 4.35 (s, 2H, Hb′, minor rotamer), 4.00 (s, 2H, Hc, major rotamer), 3.96 (s, 2H, Hc′, minor rotamer), 2.26 (s, 1H, Hd, major rotamer), 2.14 (s, 1H, Hd′, minor rotamer), 1.91 (b, 3H, He, both rotamers overlapping). Signals marked with Ha/Ha′–Hd/Hd′ correspond to rotameric pairs arising from restricted rotation about the amide bond. Rotamers are observed in a ∼55:45 ratio (major:minor).

**¹³C NMR** (101 MHz, CDCl₃) δ173.25, 140.71, 139.54, 130.31, 129.97, 128.65, 127.07, 126.16(q, J = 3.8 Hz), 125.76 (q, J = 3.8 Hz), 124.14(q, J = 271 Hz),78.23, 77.35, 73.54, 72.82, 48.85, 43.51, 35.78, 35.11

**^19^F NMR** (376 MHz, CDCl_3_) δ: −62.58Hz (d, J = 36.0 Hz, 3F)

**HRMS (ESI⁺):** *m/z* calcd for C_13_H_14_F_3_N_2_O_3_⁺ ([M+H]⁺): 271.1058; found: 271.1050

### Compound SI-5

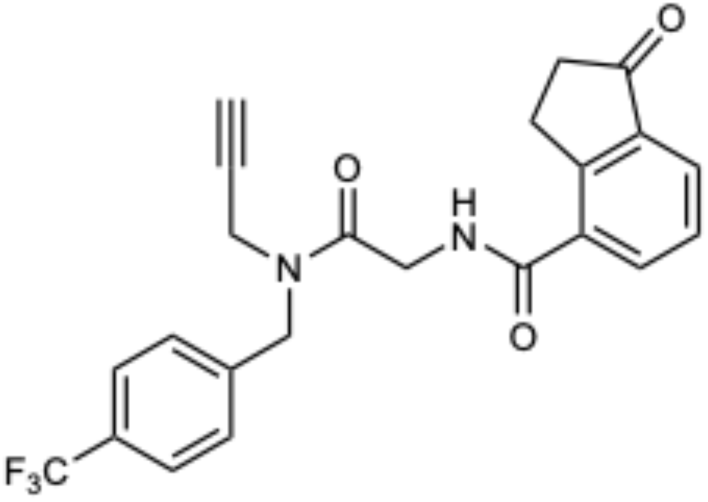

To a solution of **SI-4** (100 mg, 370 µmol, 1 equiv) in DMF was added DIPEA (191 mg, 270 µL, 1.48 mmol, 4.00 equiv), followed by 1-oxo-2,3-dihydro-1H-indene-4-carboxylic acid (72 mg, 407 µmol, 1.10 equiv) and HATU (211 mg, 555 µmol, 1.50 equiv). The reaction was stirred for 14 hours at ambient temperature. The reaction was diluted with ethyl acetate (50 mL) and washed with 1.0 M aqueous HCl (2 × 15 mL), DI water (2 × 15 mL), saturated aqueous NaHCO₃ (2 × 15 mL), and brine (2 × 15 mL). The organic layer was dried over Na₂SO₄, filtered, and the filtrate was concentrated under reduced pressure. Purification by flash chromatography (silica gel, gradient 40–60% ethyl acetate in hexanes) afforded **SI-5** as a white foam (137.7 mg, 87% yield) as a complex mixture of rotamers.

**TLC** (hexanes:ethyl acetate = 2:3): R*_f_* = 0.31 (UV)

**^1^H NMR** (600 MHz, CDCl₃) δ: 7.96 (m, 1H), 7.89 (m, 1H), 7.65 (d, J = 8.0 Hz, 1H), 7.60 (d, J = 7.9 Hz, 1H), 7.48 (t, J = 7.6 Hz, 1H), 7.42 – 7.34 (m, 2H), 7.24 (br, 1H), 4.80 (s, 2H, major rotamer), 4.76 (s, 2H, minor rotamer) 4.47 (d, J = 4.0 Hz, 2H major rotamer), 4.35 (d, J = 4.1 Hz, 1H), 4.28 (s, 1H), 4.02 (s, 1H), 3.49 – 3.42 (m, 2H), 2.72 (q, J = 5.4 Hz, 2H), 2.38 (t, J = 2.5 Hz, 1H major rotamer), 2.29 (t, J = 2.5 Hz, 1H major rotamer)

**¹³C NMR** (101 MHz, CDCl₃) δ: 206.53, 168.55, 168.35, 166.81, 154.20, 139.98, 139.02, 138.45, 133.23, 132.53, 132.43,130.54 (q, J = 32.74) 128.66, 127.90, 127.22, 126.80, 126.36 (q, J = 3.7 Hz), 125.94 (q, J = 3.7 Hz),124.06 (q, J = 247.40Hz) 77.55, 76.76, 74.35, 73.36, 49.17, 49.01, 42.04, 41.98, 36.28, 36.19, 35.18, 26.38

**^19^F NMR** (376 MHz, CDCl_3_) δ: −62.50, −62.55

**HRMS (ESI⁺):** *m/z* calcd for C_23_H_20_F_3_N_2_O_3_⁺ ([M+H]⁺): 429.1426; found: 429.1437

### Compound SI-6

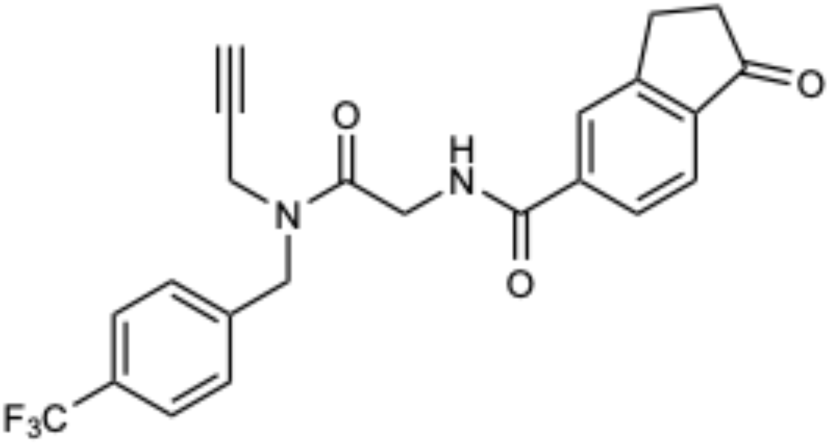

To a solution of **SI-4** (60 mg, 222 µmol, 1 equiv) in DMF was added DIPEA (57 mg, 77 µL, 444 µmol, 4.00 equiv), followed by 1-oxo-2,3-dihydro-1H-indene-5-carboxylic acid (43 mg, 244 µmol, 1.10 equiv) and HATU (127 mg, 333 µmol, 1.50 equiv). The reaction mixture was stirred at ambient temperature for 14 hours. The reaction mixture was diluted with ethyl acetate (50 mL) and sequentially washed with 1.0 M aqueous HCl (2 × 15 mL), DI water (2 × 15 mL), saturated aqueous NaHCO₃ (2 × 15 mL), and brine (2 × 15 mL). The organic layer was dried over Na₂SO₄, filtered, and the filtrate was concentrated under reduced pressure. The crude product was purified by flash column chromatography (silica gel, gradient 40–60% ethyl acetate in hexanes) to afford the product as a white foam (87.6 mg, 92% yield) as a complex mixture of rotamers.

**TLC** (hexanes:ethyl acetate = 2:3): R*_f_* = 0.29 (UV)

**^1^H NMR** (600 MHz, CDCl₃) δ: 8.12 (m, 1H), 7.85 (m, 1H), 7.59 (m, 3H), 7.41 (m, 3H), 4.78 (s, 2H, major rotamer), 4.76 (s, 2H, minor rotamer), 4.45 (t, J = 4.3 Hz, 2H, major rotamer), 4.33 (t, J = 4.1 Hz, 2H, minor rotamer), 4.26 (t, J = 2.6 Hz, 2H, minor rotamer), 4.02 (t, J = 3.1 Hz, 1H, major rotamer), 3.22 – 3.13 (m, 2H), 2.77 – 2.67(m, 2H), 2.40 – 2.33 (m, 1H, minor rotamer), 2.31 – 2.24 (m, 1H, major rotamer)

**¹³C NMR** (150 MHz, CDCl₃) δ: 206.42, 168.61, 168.43, 166.66, 166.37, 158.54, 155.38, 140.11, 139.55, 139.40, 139.16, 137.38, 133.81, 133.31, 130.28, 128.67, 127.22, 126.32 (q, *J* = 3.7 Hz), 125.89 (q, *J* = 3.2 Hz), 124.08, 123.94 (q, J = 279 Hz), 122.16, 77.67, 77.58, 76.79, 74.33, 74.24, 73.31, 73.25, 49.13, 48.97, 42.13, 42.06, 36.58, 36.19, 35.08, 26.04, 25.91

**^19^F NMR** (376 MHz, CDCl_3_) δ: −62.48(major rotamer), −62.54 (minor rotamer)

**HRMS (ESI⁺):** *m/z* calcd for C_23_H_20_F_3_N_2_O_3_⁺ ([M+H]⁺): 429.1426; found: 429.1438

### iso-CA-Nin-Alk

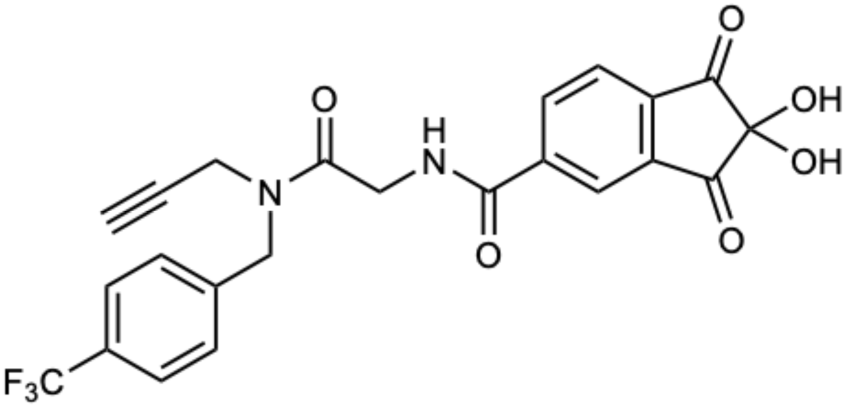

Selenium dioxide (79 mg, 716 µmol, 2.1 equiv) was added to a solution of **SI-6** (146 mg, 341 µmol, 1 equiv) in 1,4-dioxane (1.5 mL) and DI water (150 μL). The reaction mixture was heated to reflux in a 96 °C oil bath for 4 hours. After cooling to ambient temperature, the mixture was diluted with ethyl acetate (15 mL) and washed successively with DI water (2 × 10 mL), 1 M aqueous HCl (2 × 10 mL). The organic layer was washed with brine (1 × 10 mL), dried over Na₂SO₄, filtered, and the filtrate was concentrated under reduced pressure. The crude mixture was purified by reverse-phase HPLC (25-50% MeCN with 0.1% TFA in milliQ water with 0.1% TFA), and lyophilization of the collected fractions yielded the product as a white powder (18 mg, 11% yield) as a complex mixture of rotamers.

**TLC** (hexanes:acetone = 1:2): R*_f_* = 0.39 (UV)

**¹H NMR** (600 MHz, 10:1 CD_3_CN + D_2_O) δ 7.98 (2H, m), 7.67 (3H, m), 7.49 (3H, m), 4.75 (2H, m), 4.29 (2H, m), 4.17 (2H, m), 2.60 (1H, s)

**¹³C NMR** (151 MHz, 10:1 CD_3_CN + D_2_O) δ: 169.78, 169.64, 169.19, 168.83, 166.68, 153.87, 142.62, 141.88, 136.91, 136.70, 133.61, 131.39, 130.95, 130.53, 129.58, 129.21, 128.41, 126.43, 126.13, 124.30, 123.56, 79.43, 78.86, 74.53, 73.36, 50.20, 49.94, 42.32, 42.16, 37.46, 35.89

**^19^F NMR** (376 MHz, CD_3_CN + 10% D_2_O) δ: −63.68(major rotamer), −63.71 (minor rotamer), −76.55 (reference TFA)

**HRMS (ESI⁺):** *m/z* calcd for C_23_H_18_F_3_N_2_O_6_⁺ ([M+H]⁺): 475.1117; found: 475.1113.

### CA-Nin-Alk

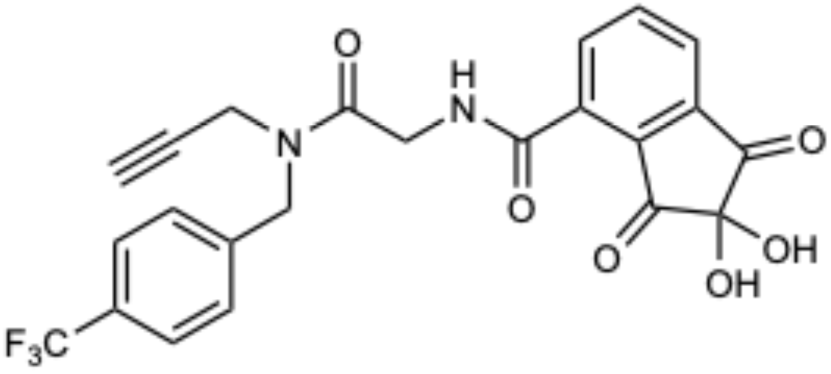

Selenium dioxide (140 mg, 1.3 mmol, 2.2 equiv) was added to a solution of **SI-5** (245 mg, 590 µmol, 1 equiv) in 1,4-dioxane (2.6 mL) and DI water (260 μL). The reaction mixture was heated to reflux in an oil bath at 96 °C for 4 hours. After cooling to ambient temperature, the mixture was diluted with ethyl acetate (15 mL) and washed successively with DI water (2 × 10 mL), 1 M aqueous HCl (2 × 10 mL). The organic layer was washed with brine (1 × 10 mL), dried over Na₂SO₄, filtered, and the filtrate was concentrated under reduced pressure. The crude mixture was purified by reverse-phase HPLC (25-50% MeCN with 0.1% TFA in milliQ water with 0.1% TFA), and lyophilization of the collected fractions yielded the product as a white powder (26 mg, 10 % yield) as a complex mixture of rotamers.

**TLC** (hexanes:acetone = 1:2): R*_f_* = 0.42 (UV)

**¹H NMR** (600 MHz, 10:1 CD_3_CN + D_2_O) δ: δ 8.05 (2H, m), 7.67 (4H, m), 7.51 (2H, m), 4.78 (2H, m), 4.40 (2H, m), 4.19 (2H, m), 2.68 (1H, m), 2.55 (2H, s)

**¹³C NMR** (151 MHz, 10:1 CD_3_CN + D_2_O₃) δ: δ 169.38, 169.27, 166.87, 165.13, 162.89, 149.24, 146.14, 140.70, 138.49, 137.80, 135.73, 135.59, 134.43, 131.75, 130.21, 129.79, 129.72, 129.50, 128.98, 128.21, 126.85, 126.22, 125.92, 124.10, 79.12, 74.43, 73.23, 50.12, 49.89, 49.74, 42.65, 42.52, 40.34, 37.31, 35.76

**^19^F NMR** (376 MHz, CD_3_CN + 10% D_2_O) δ: −63.71(major rotamer), −63.74 (minor rotamer), −76.55 (reference TFA)

**HRMS (ESI⁺):** *m/z* calcd for C_23_H_18_F_3_N_2_O_6_⁺ ([M+H]⁺): 475.1117; found: 475.1122

## Spectral Data

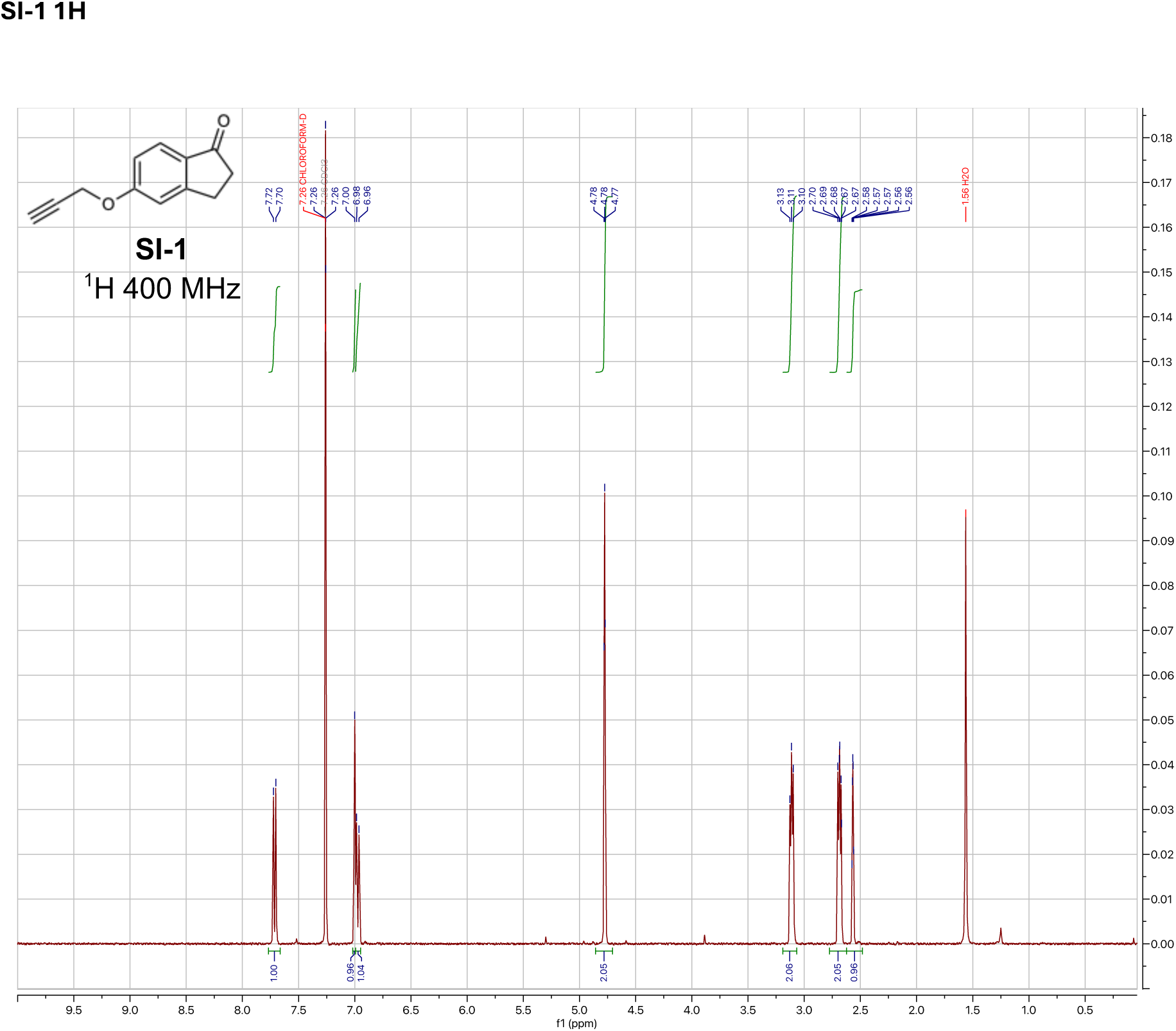

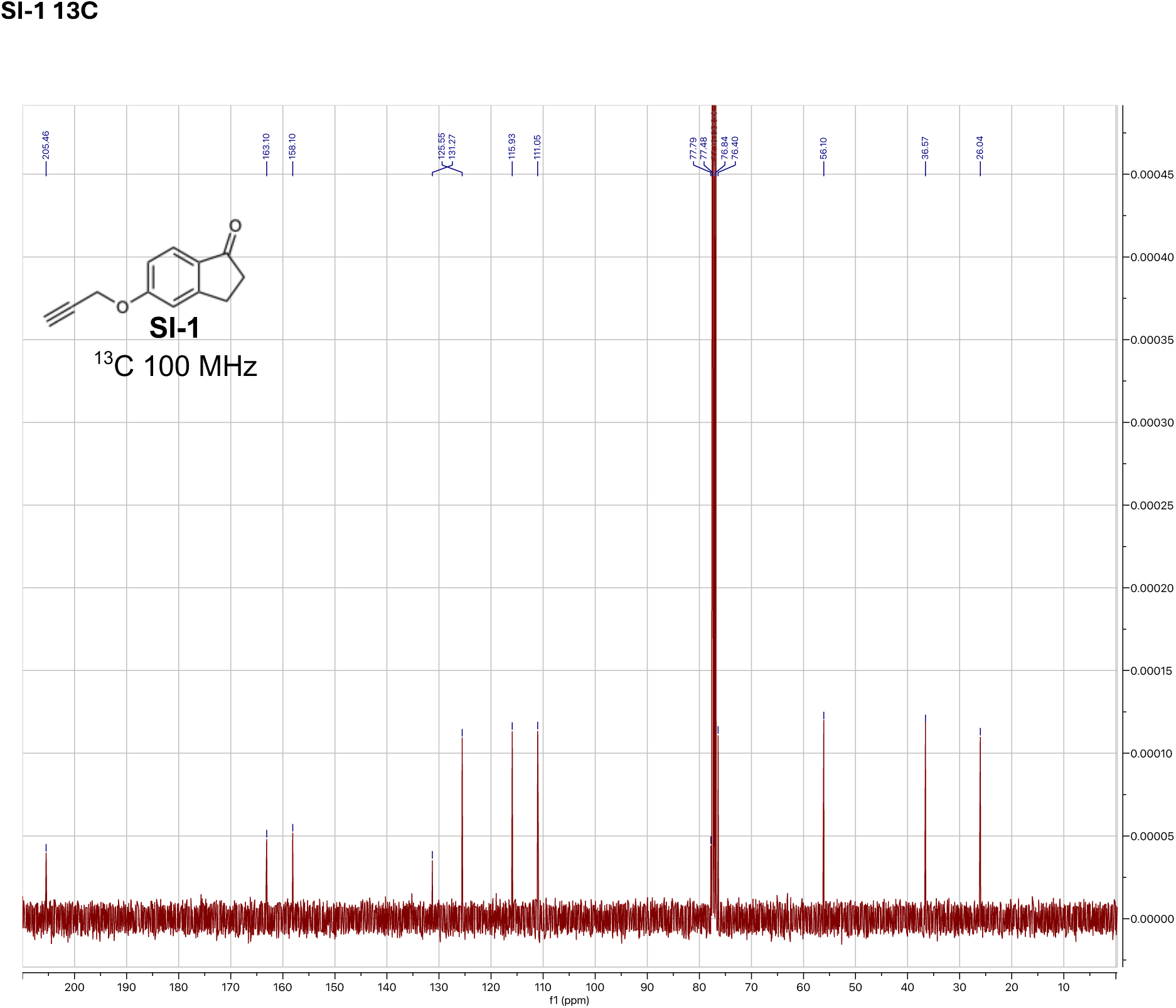

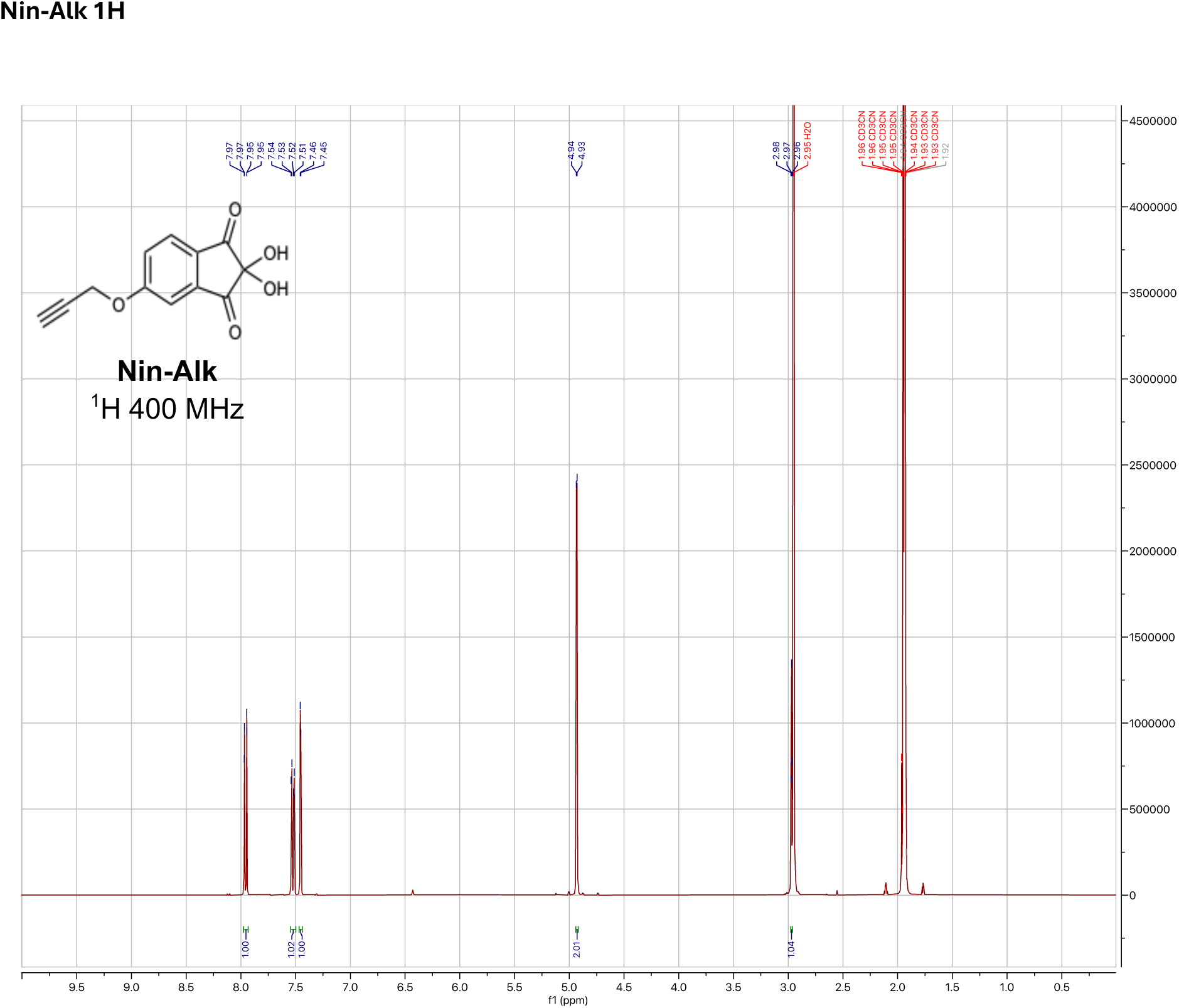

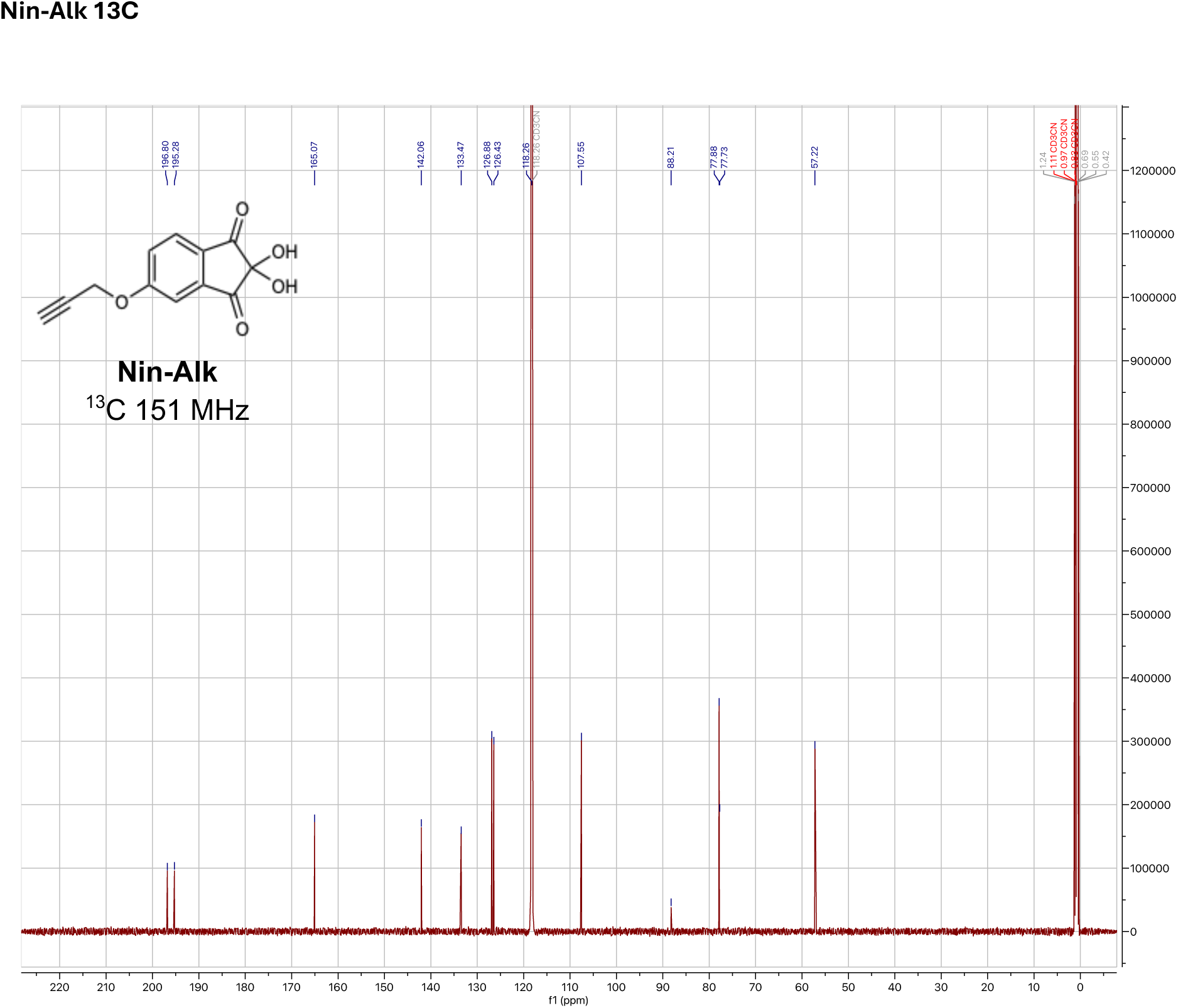

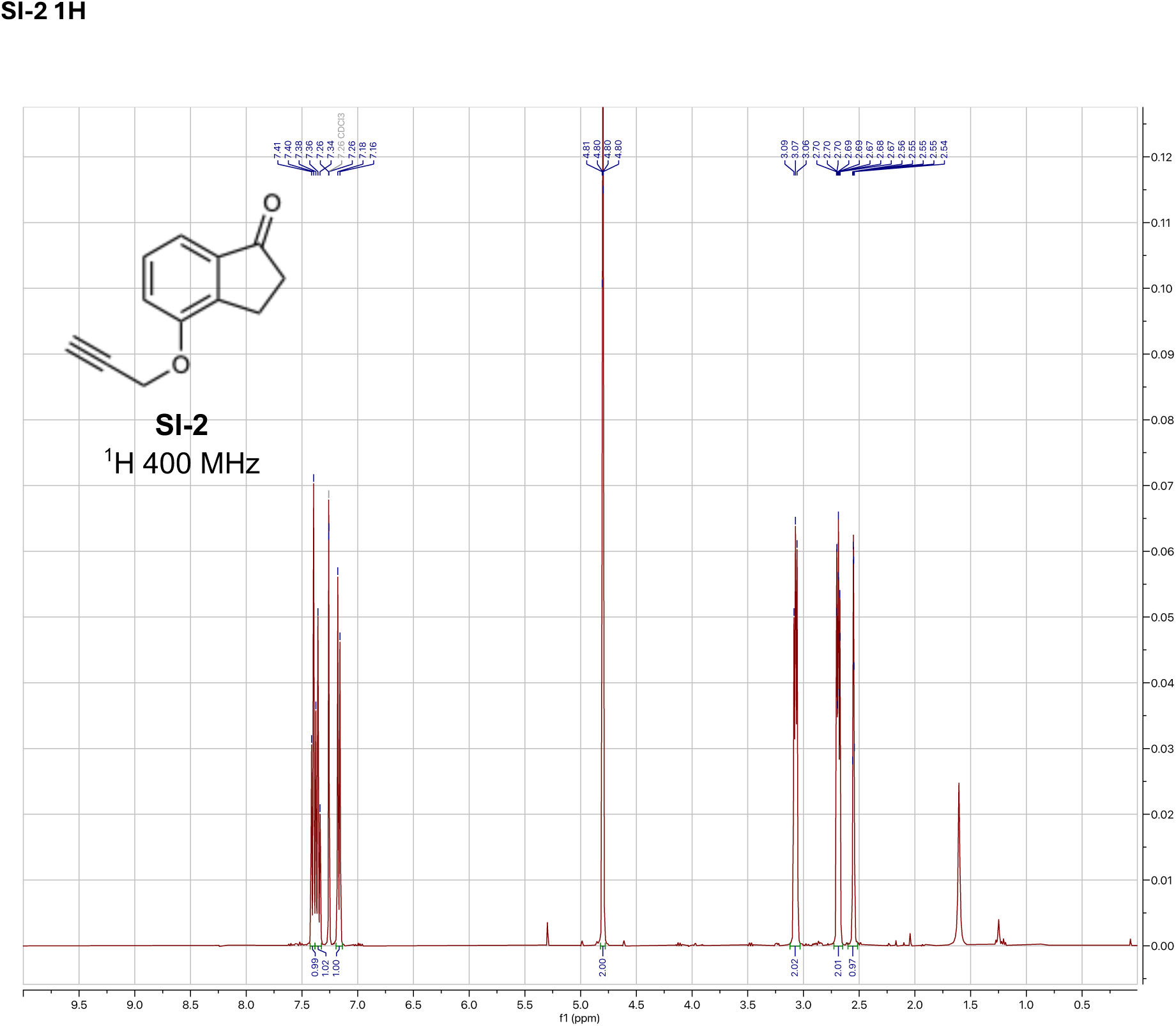

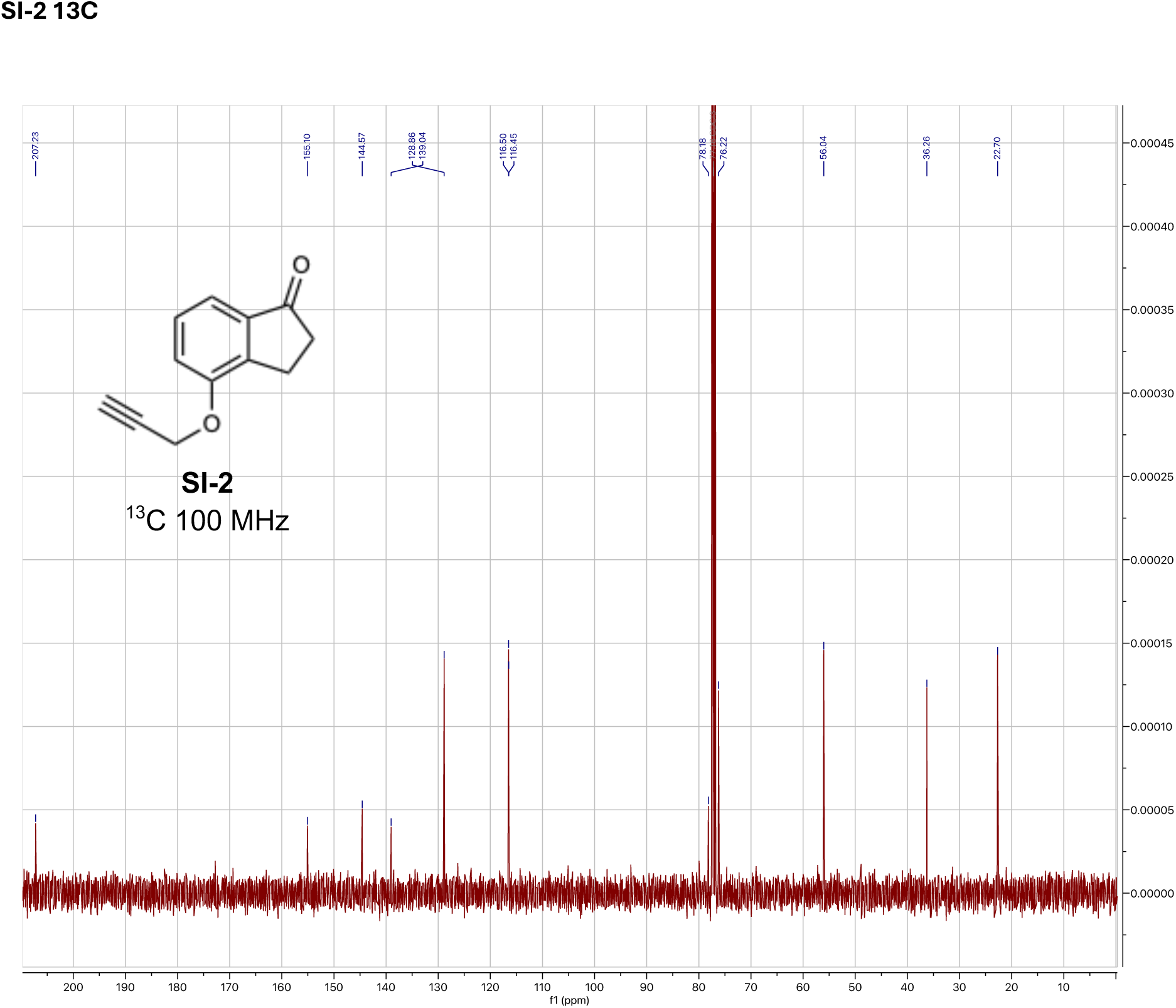

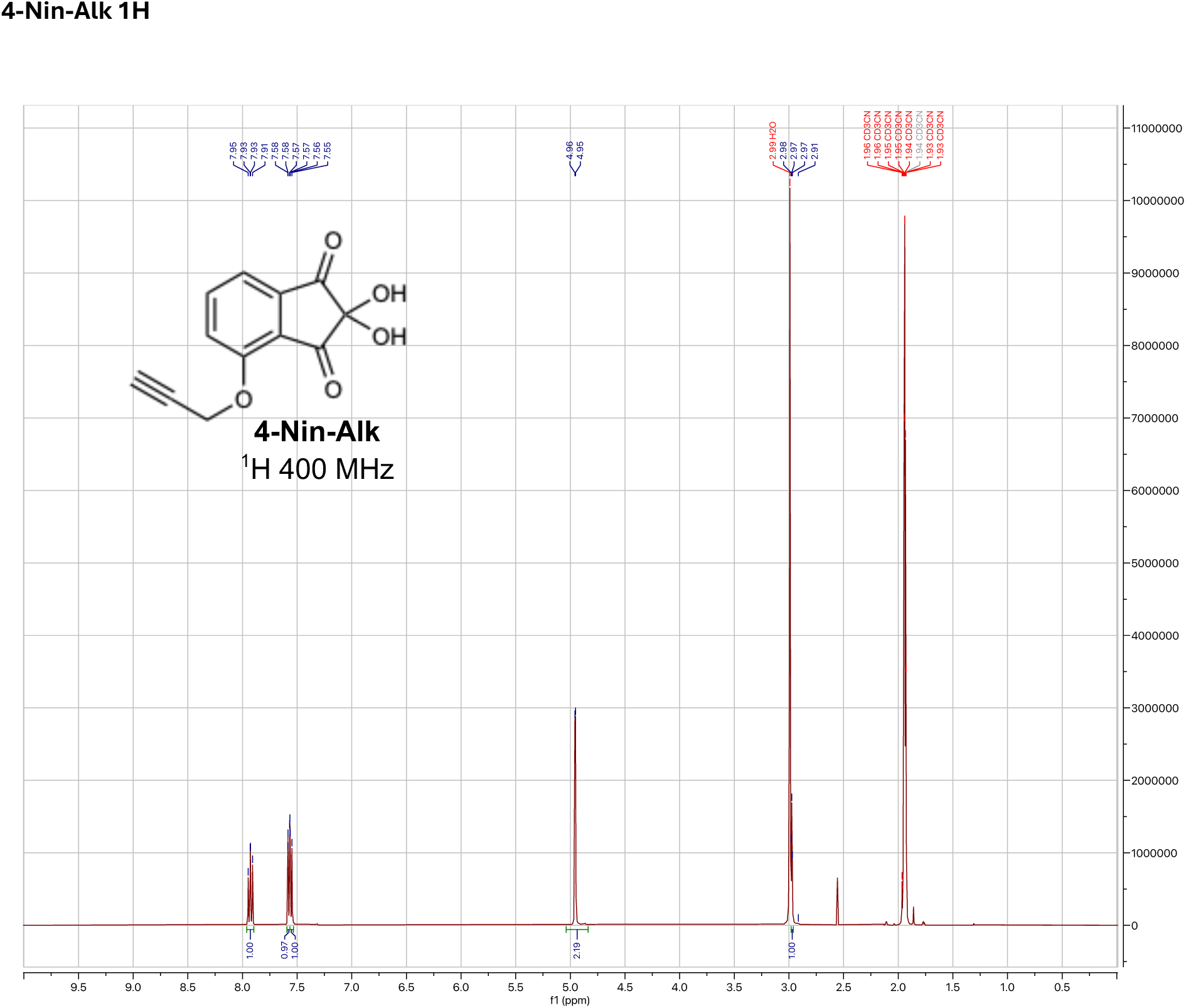

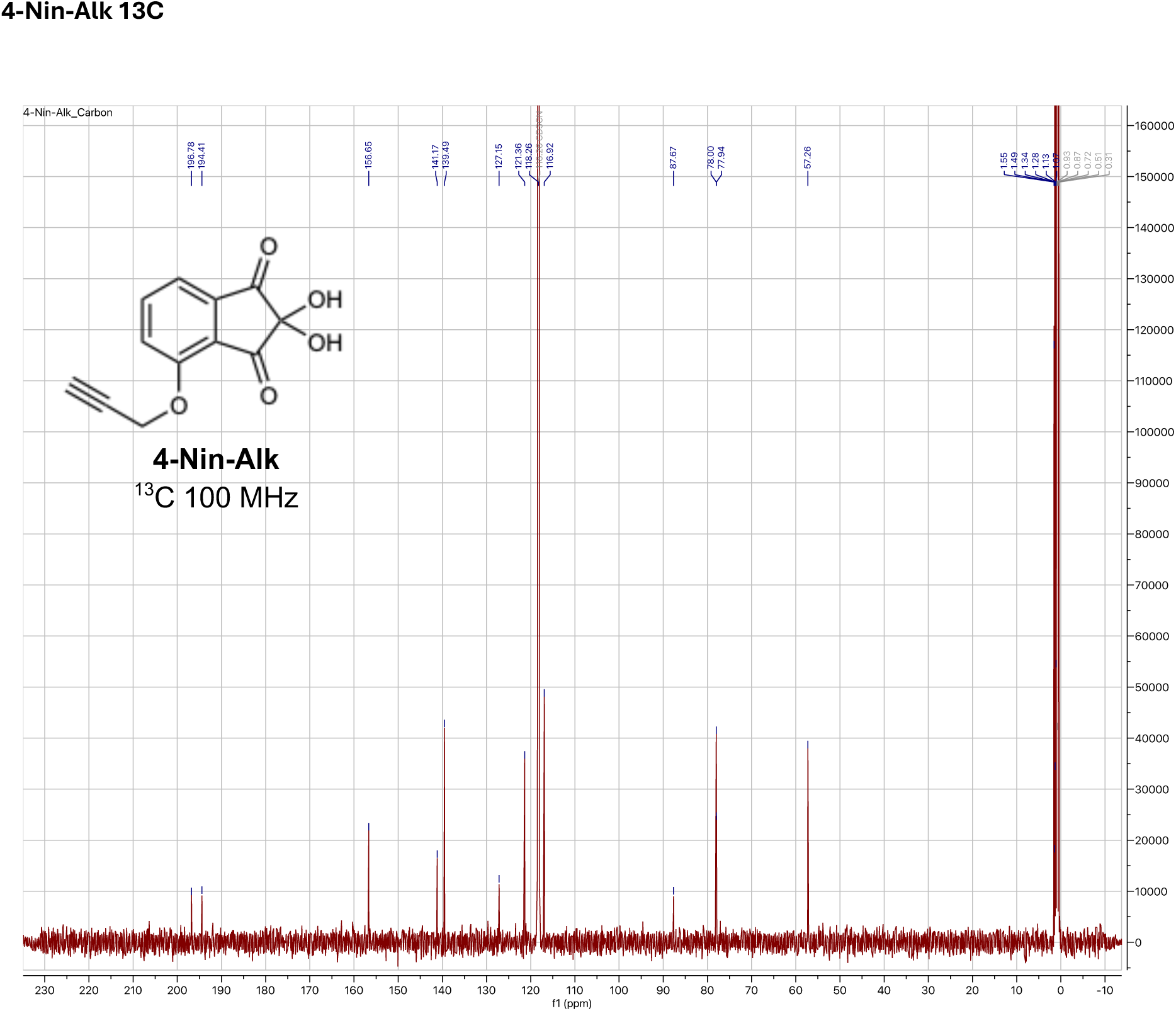

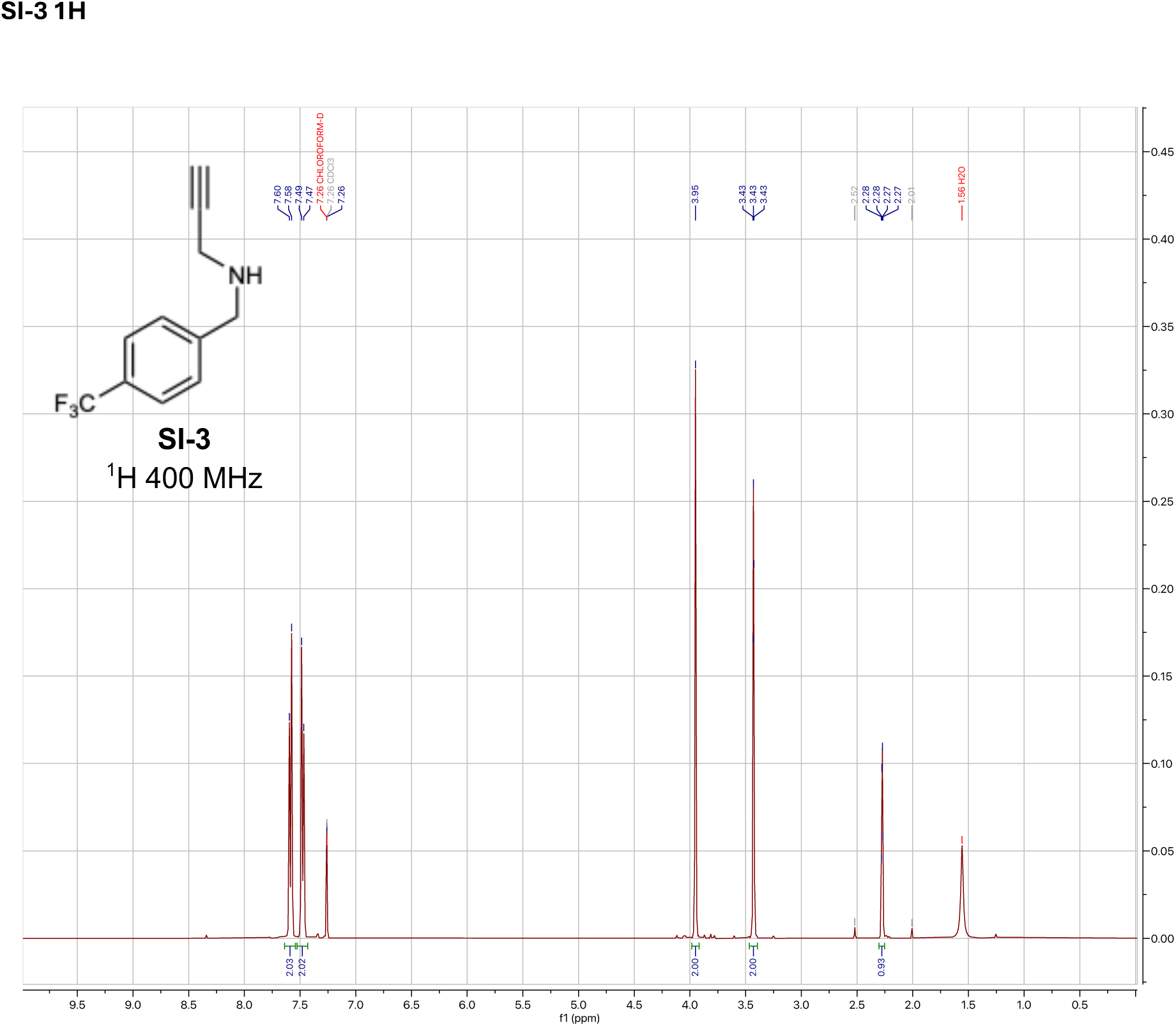

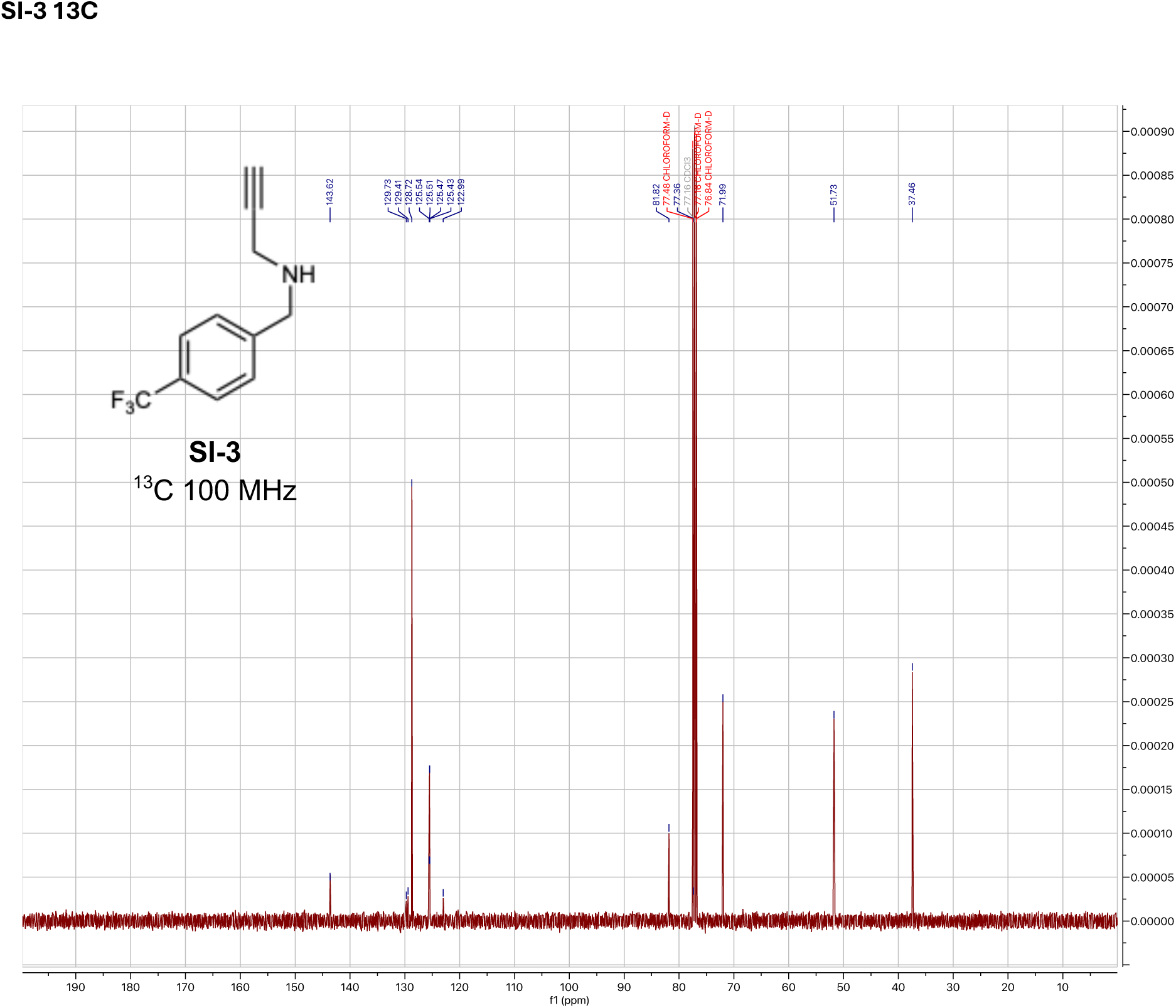

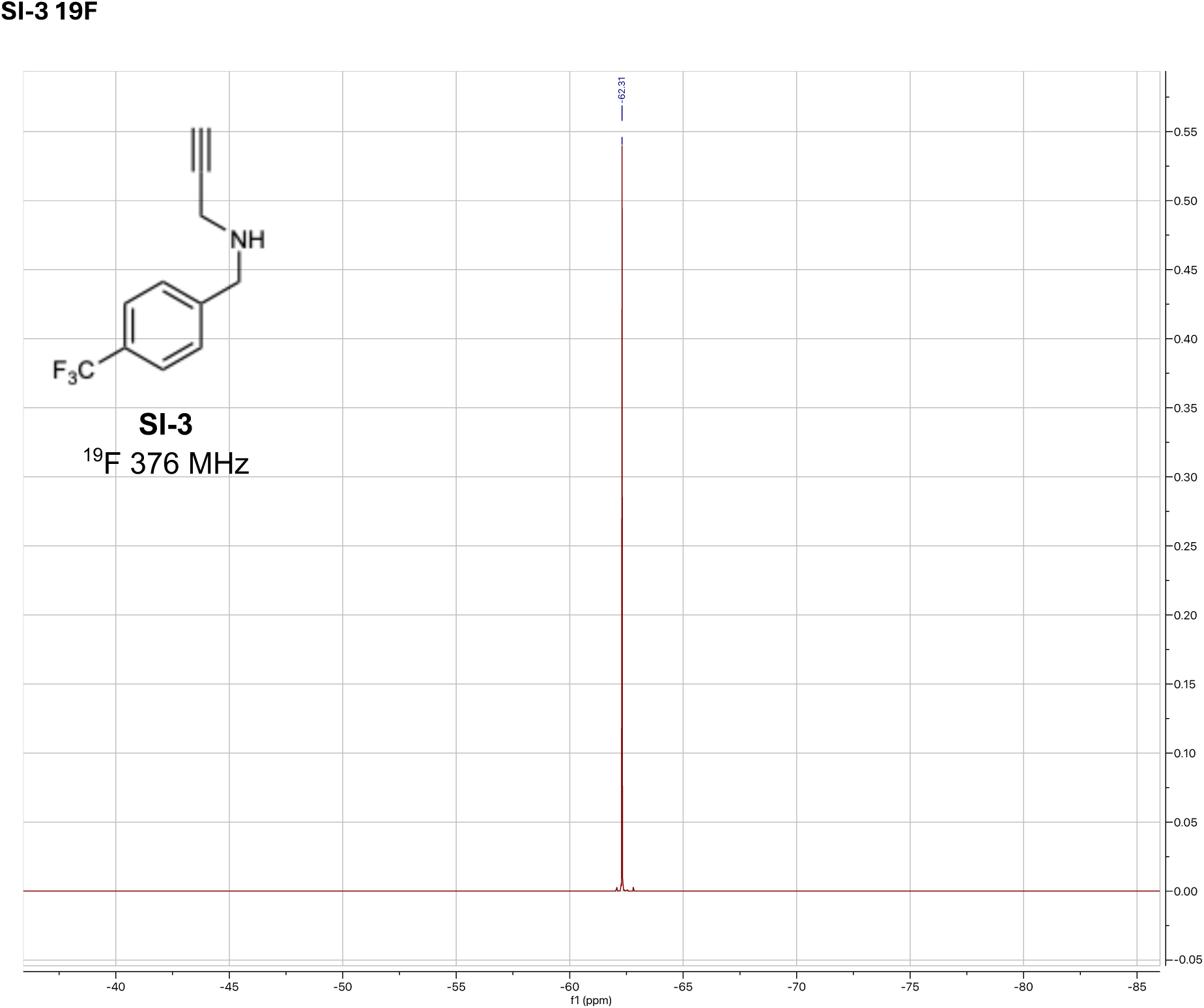

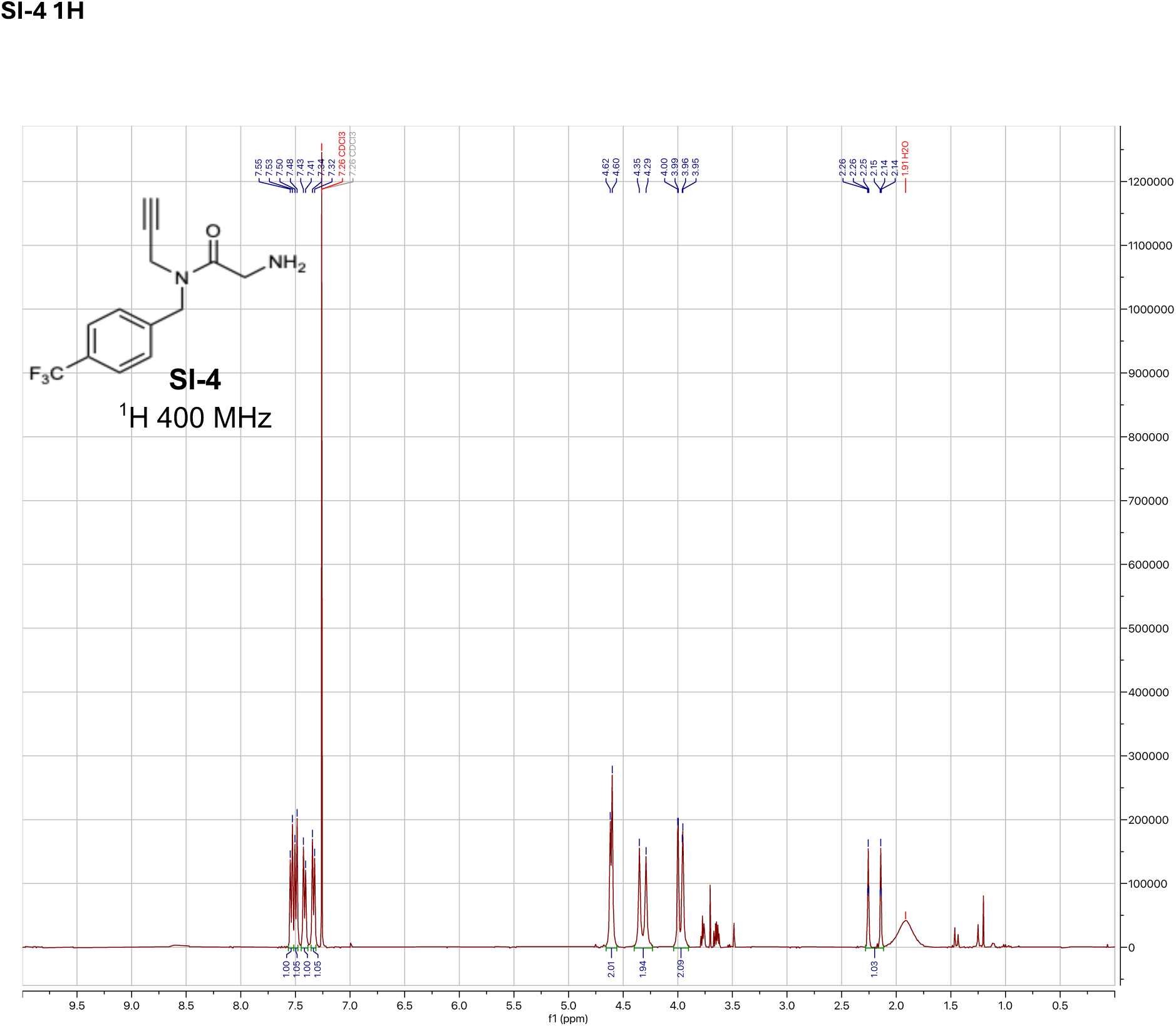

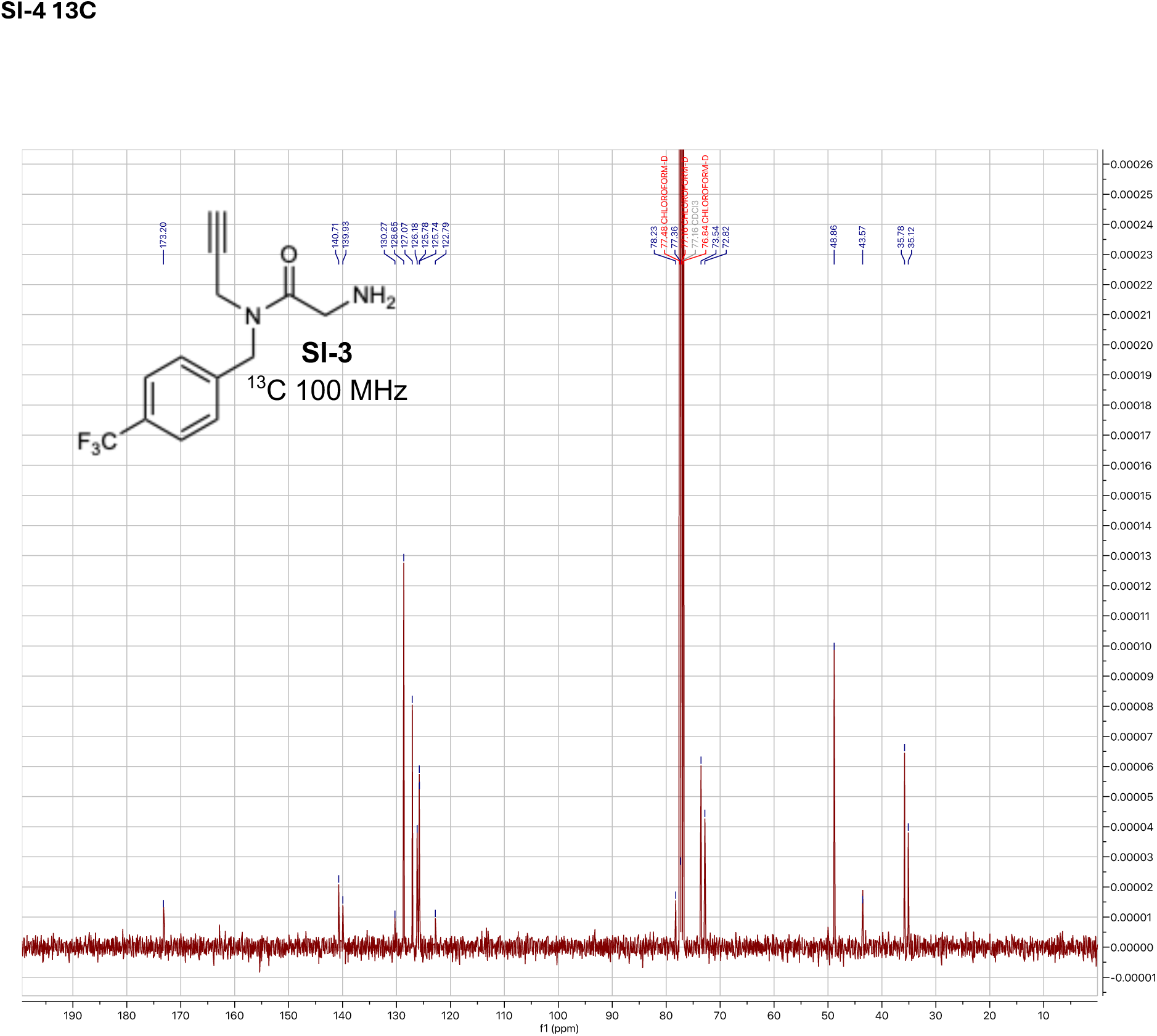

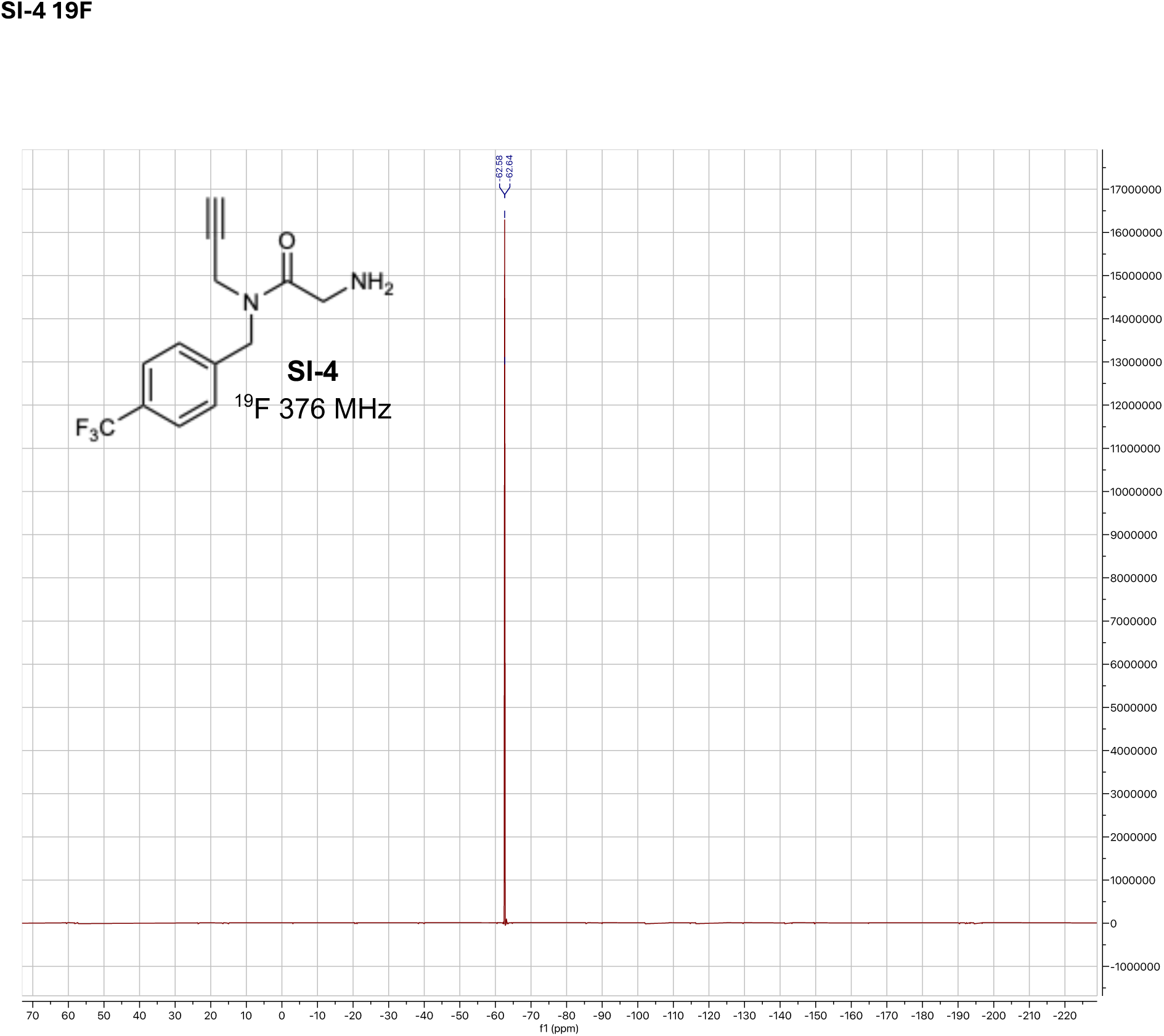

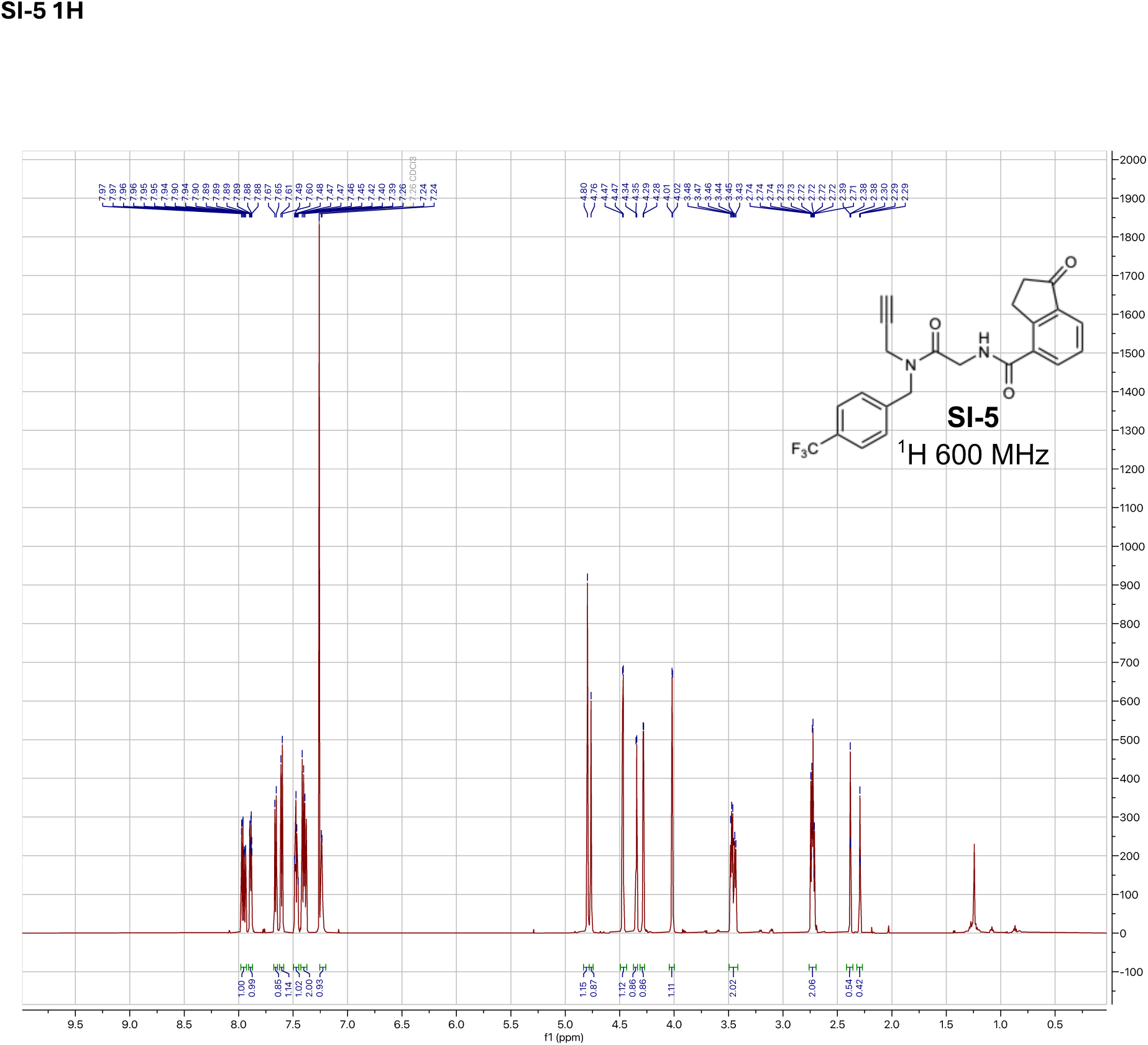

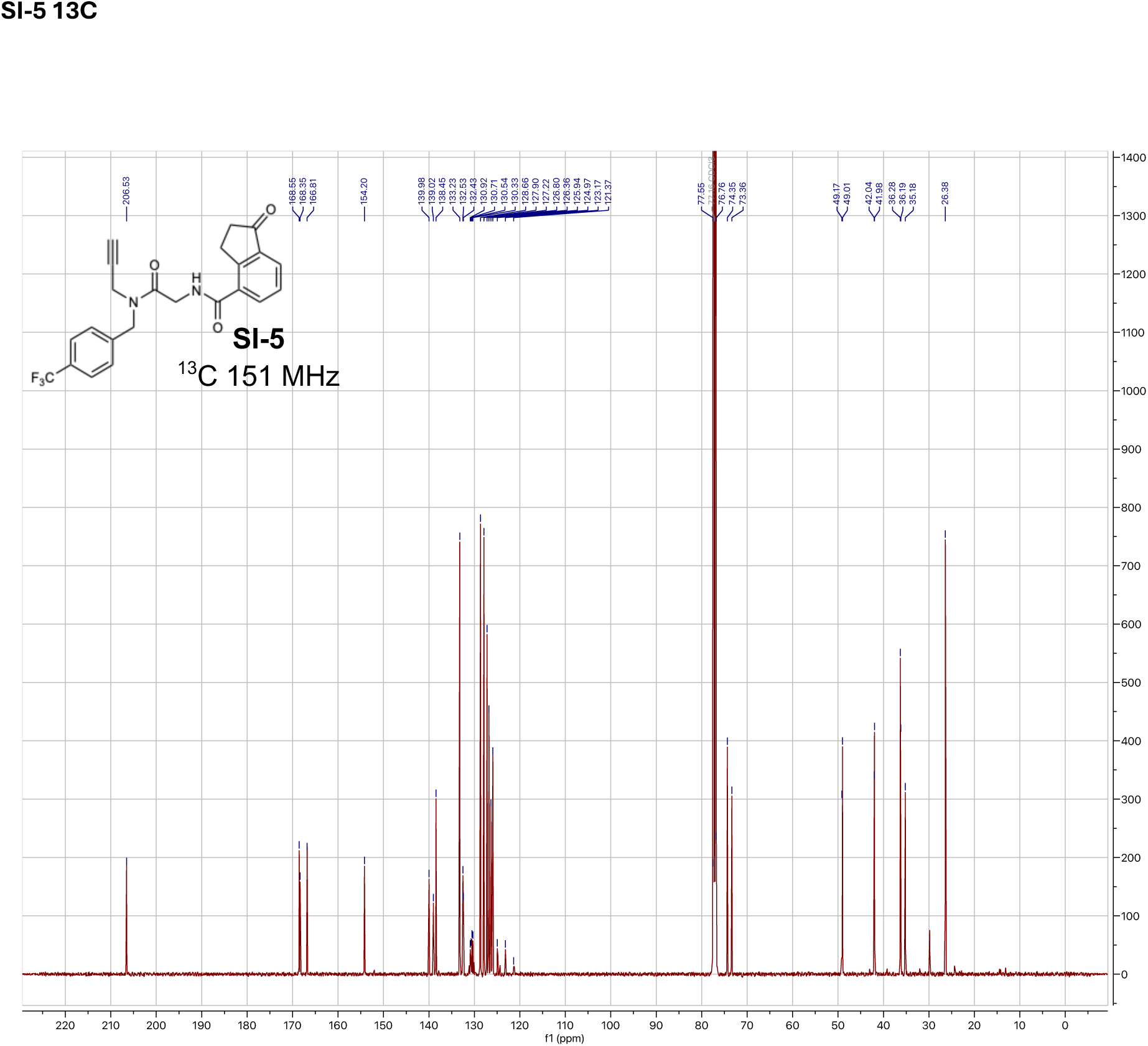

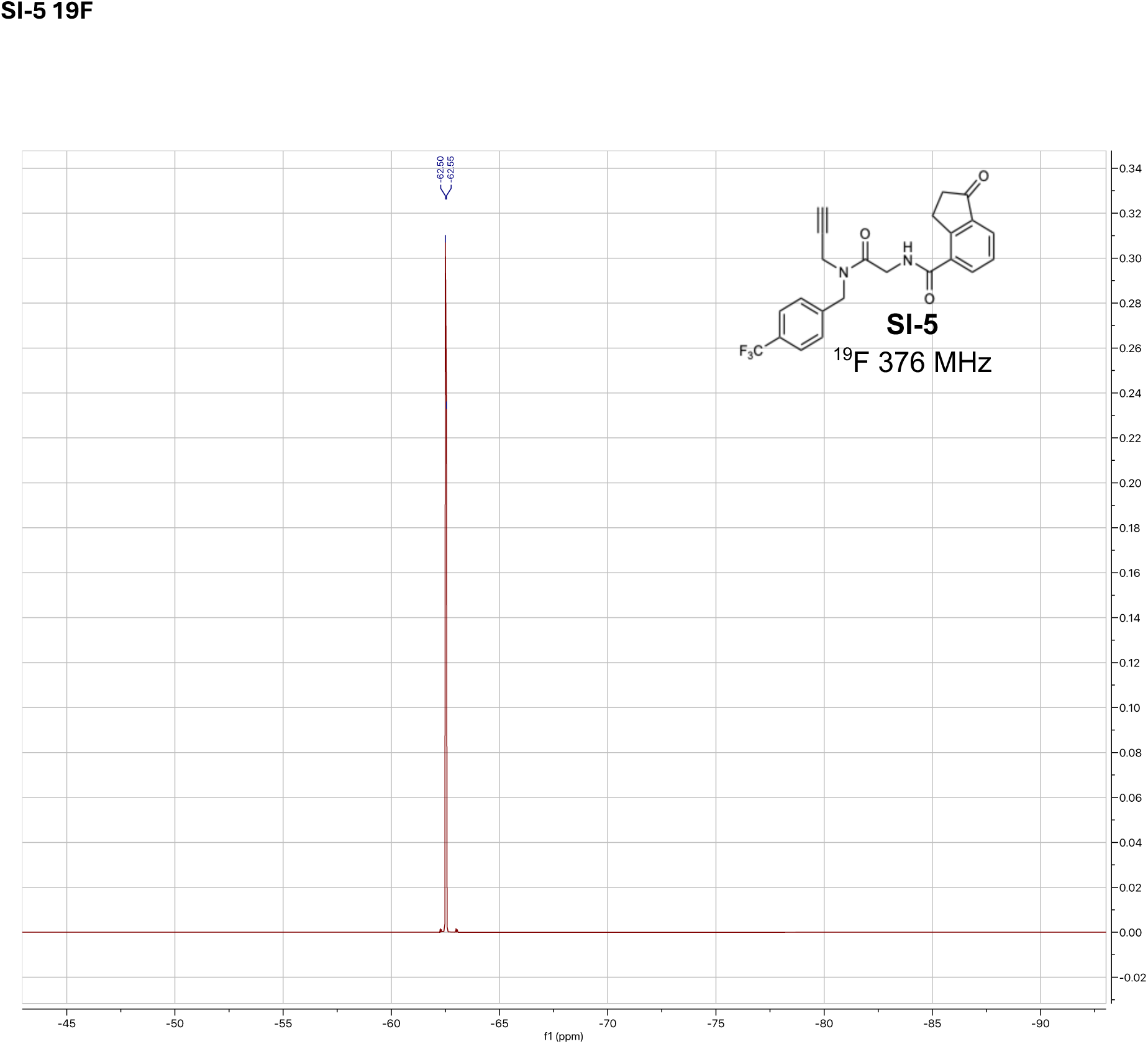

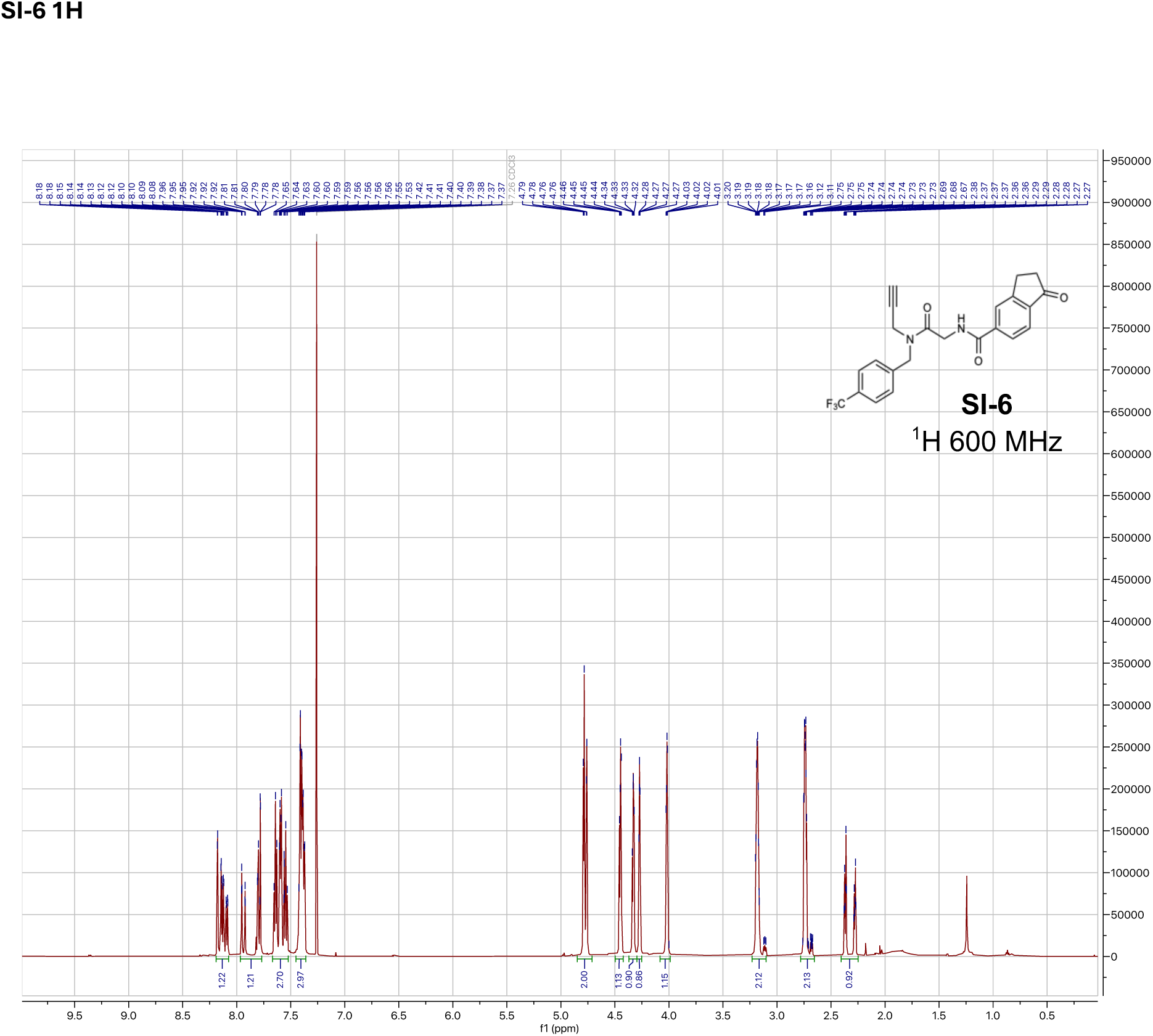

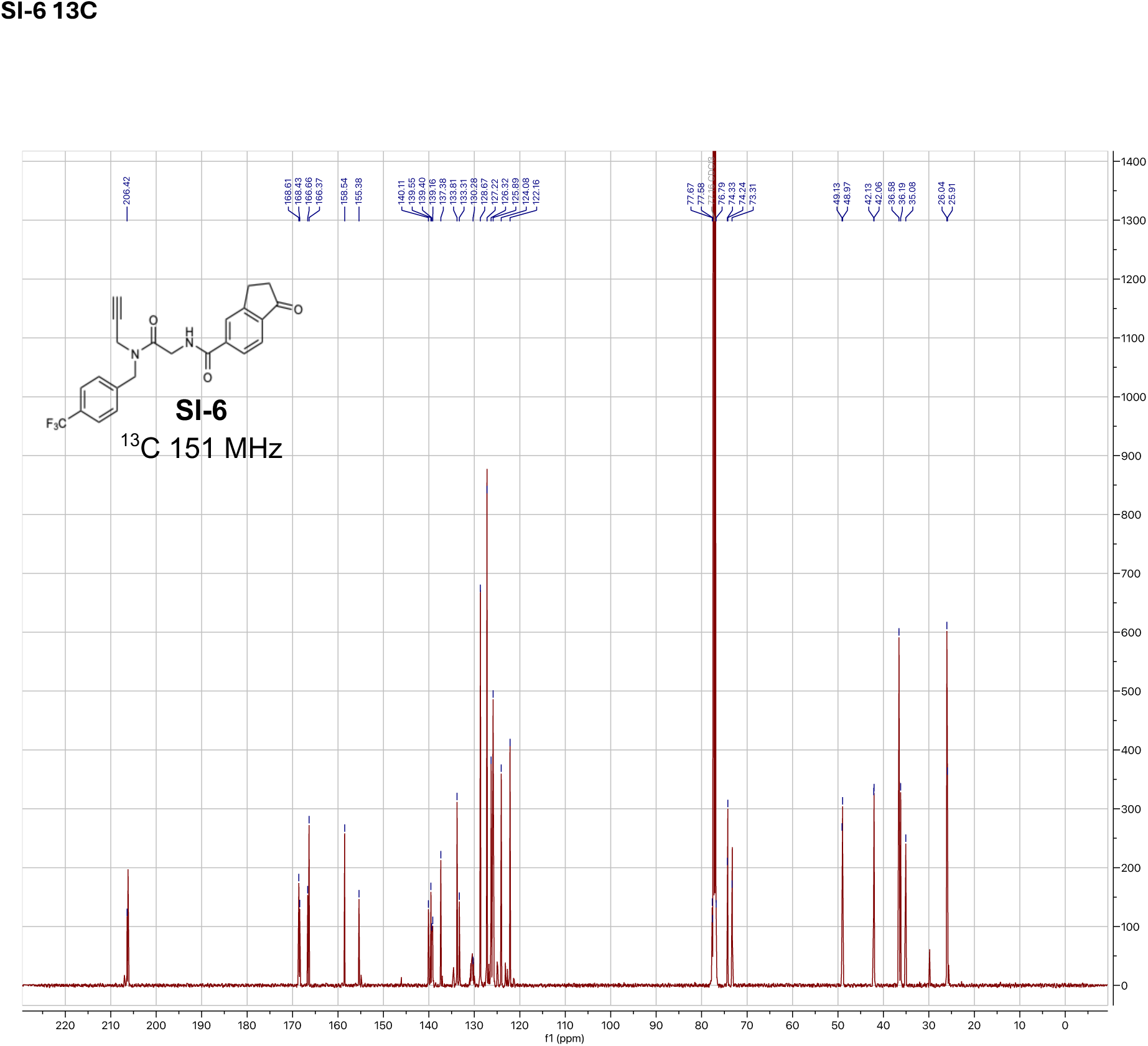

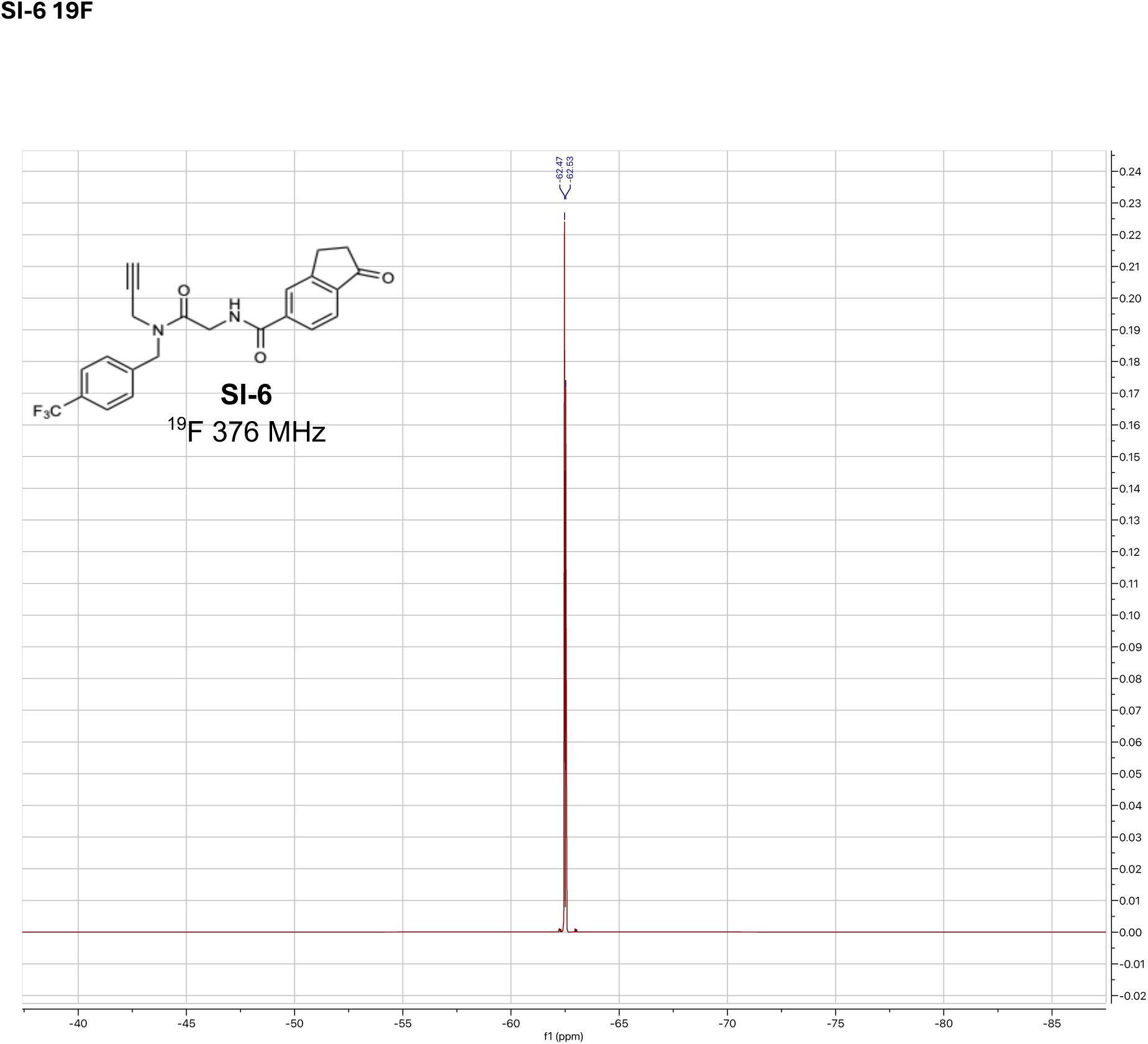

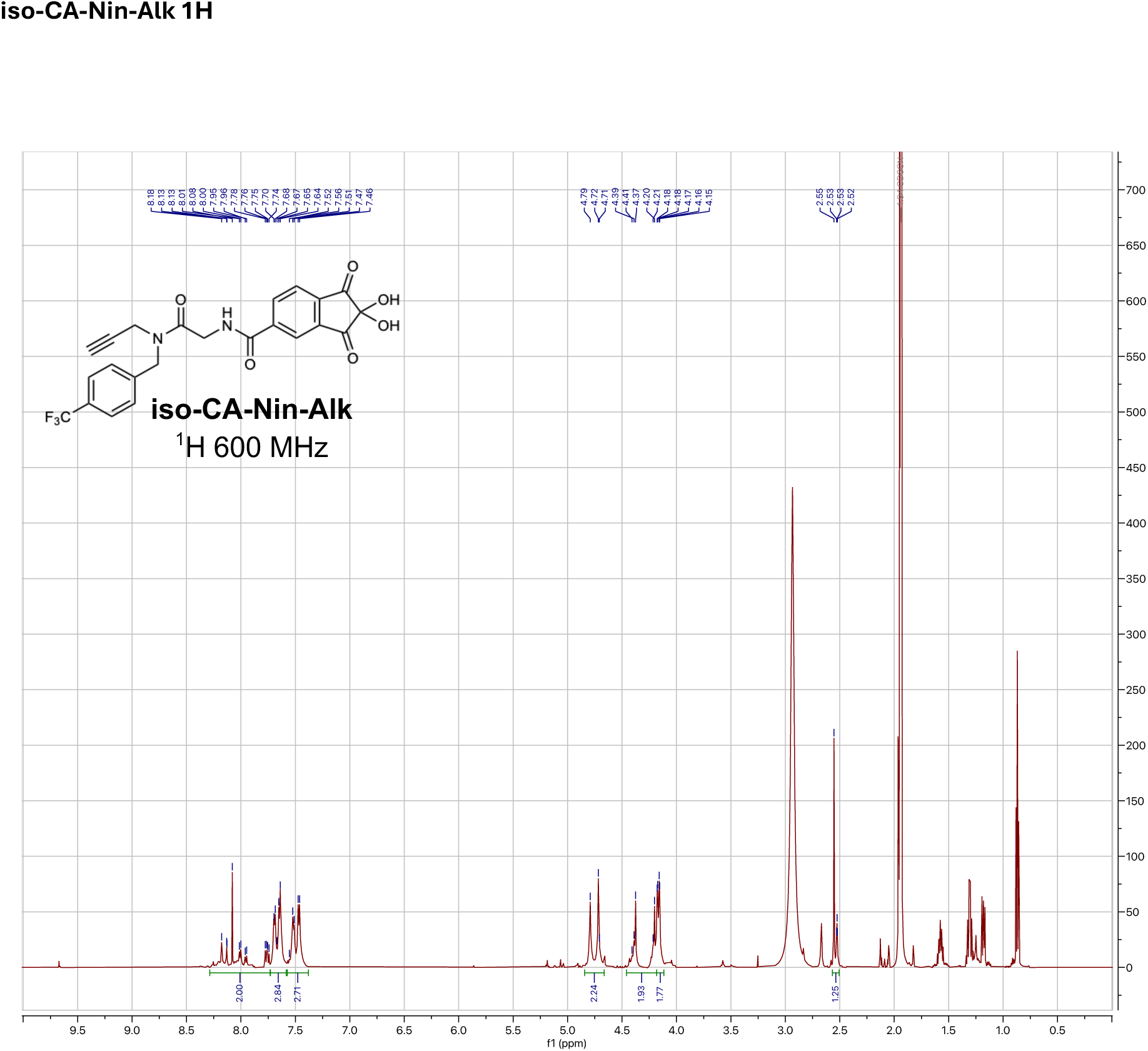

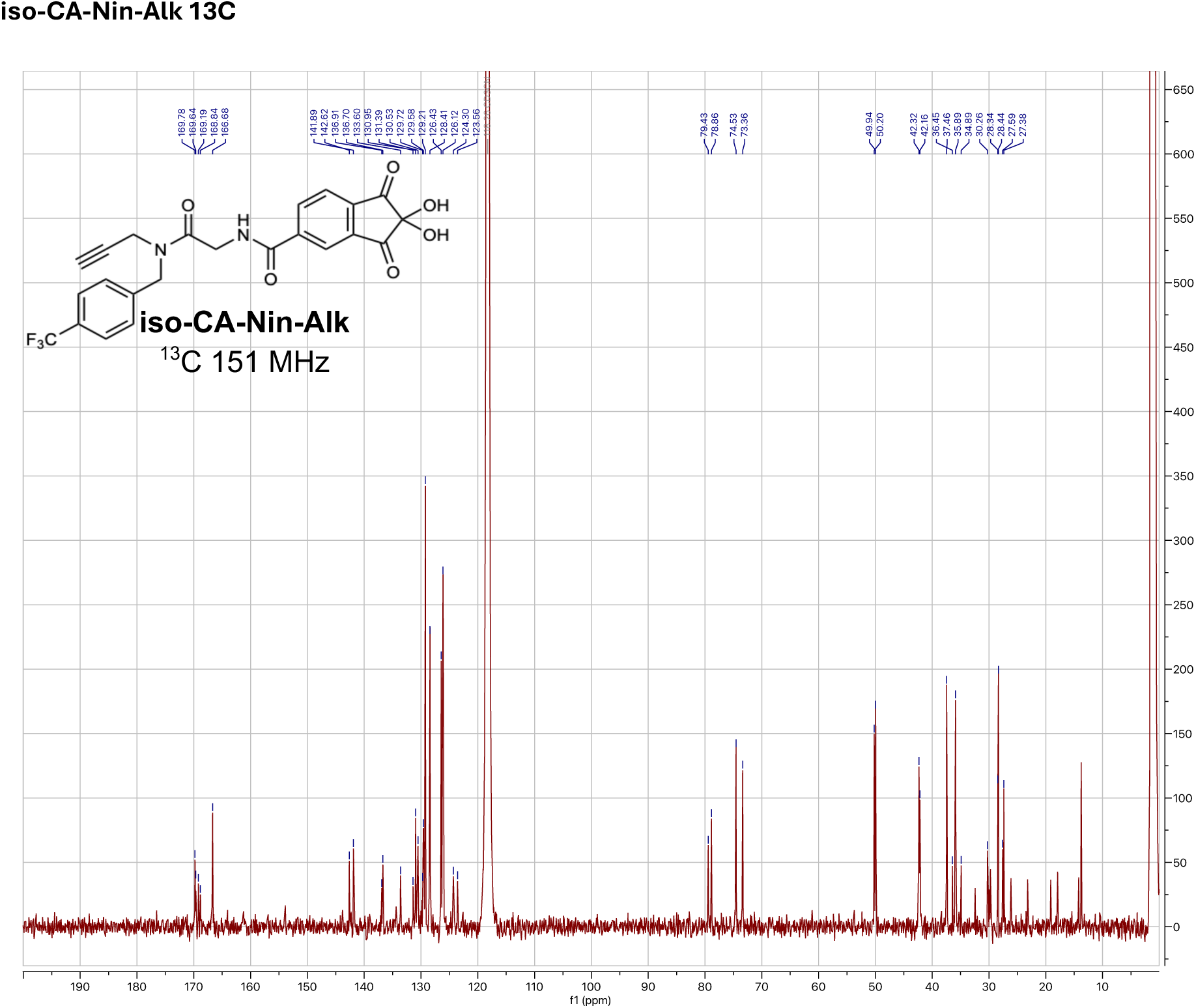

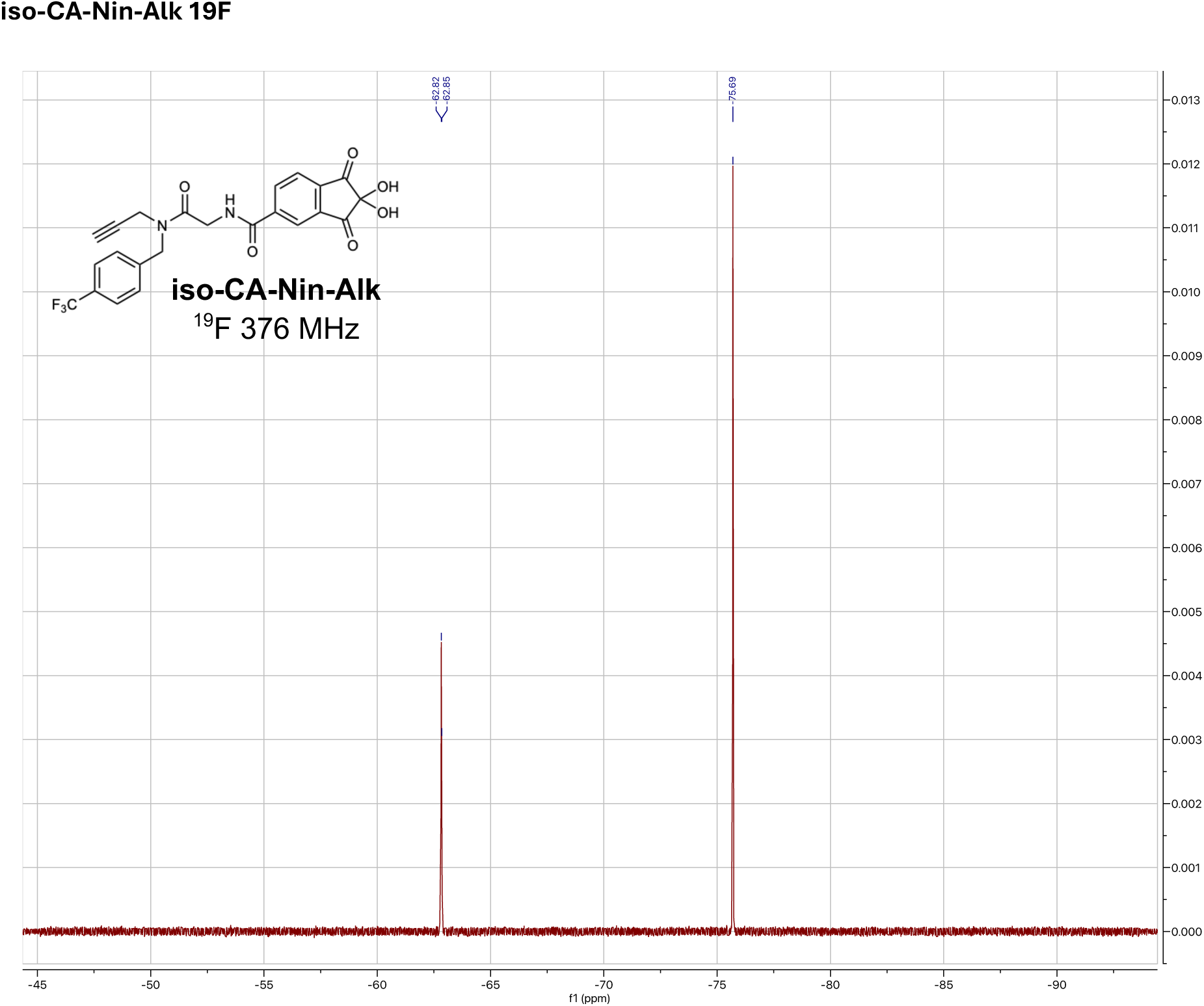

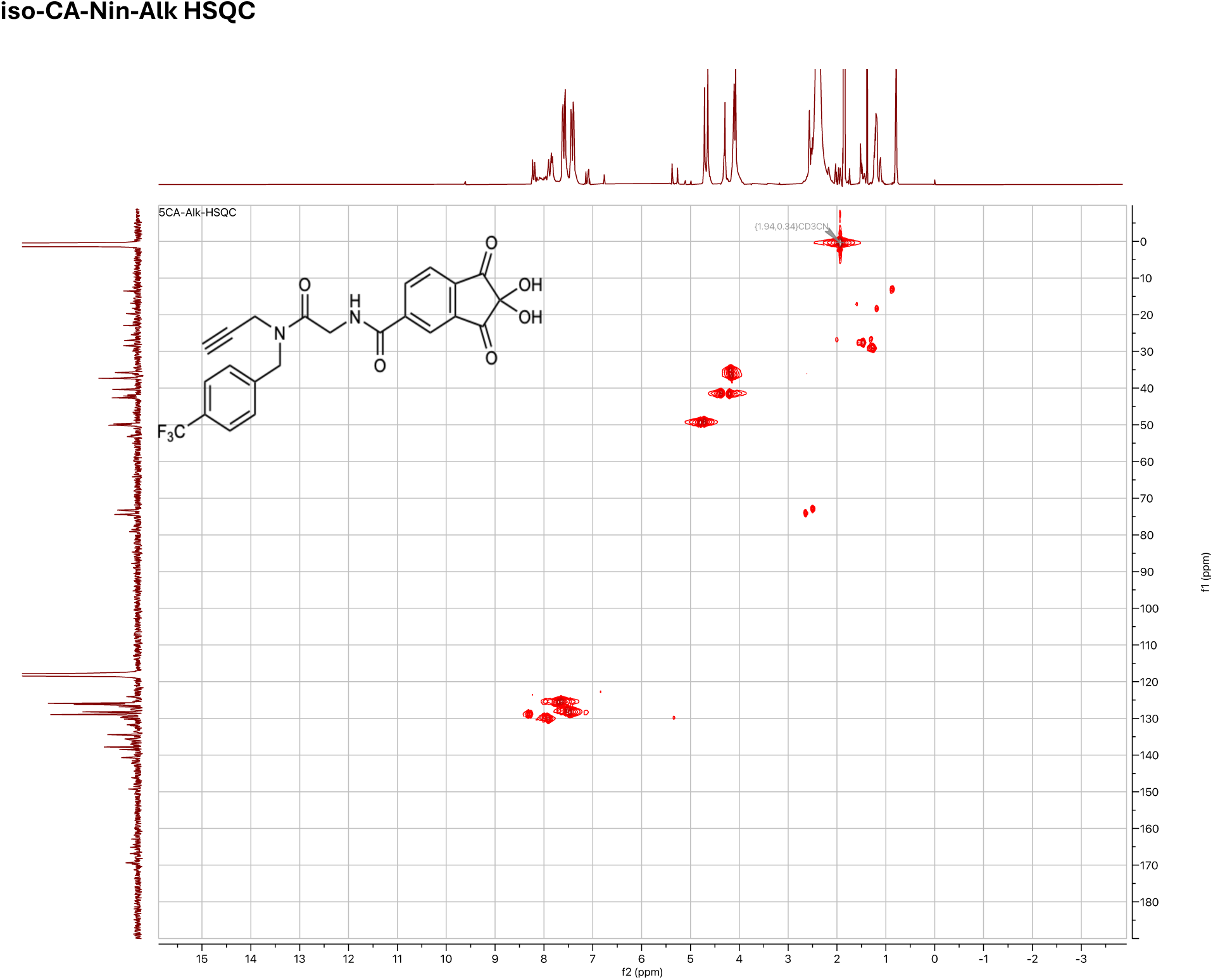

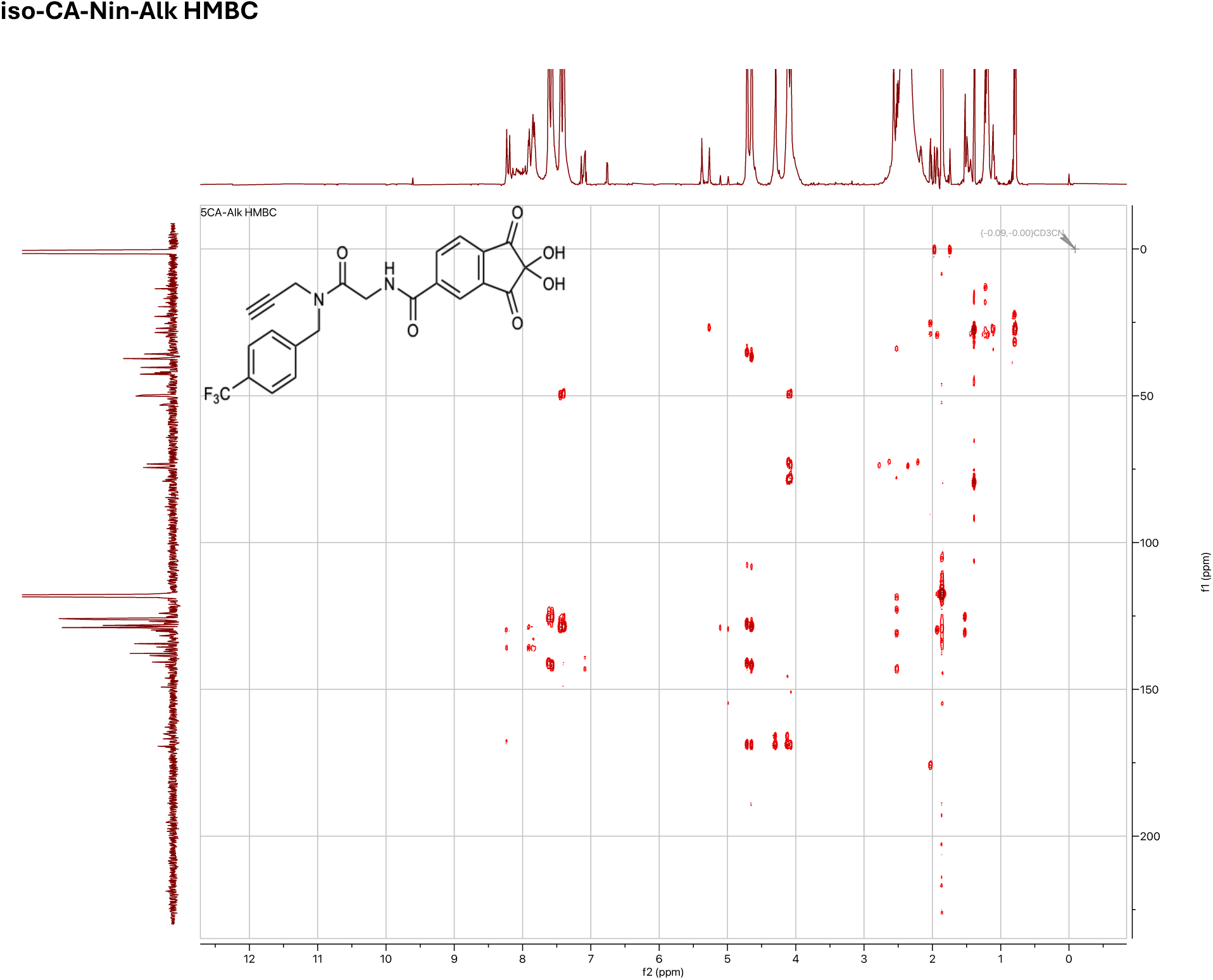

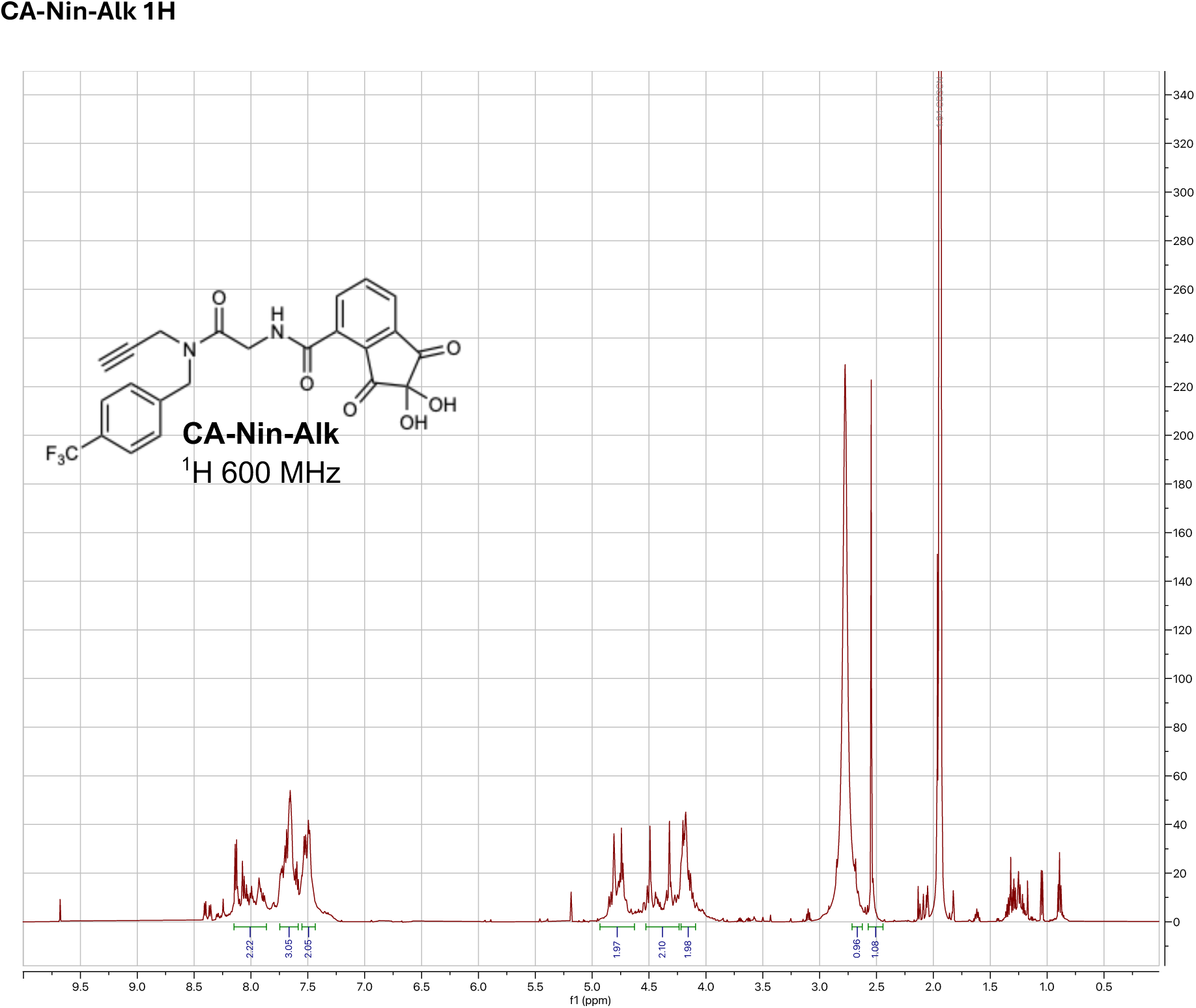

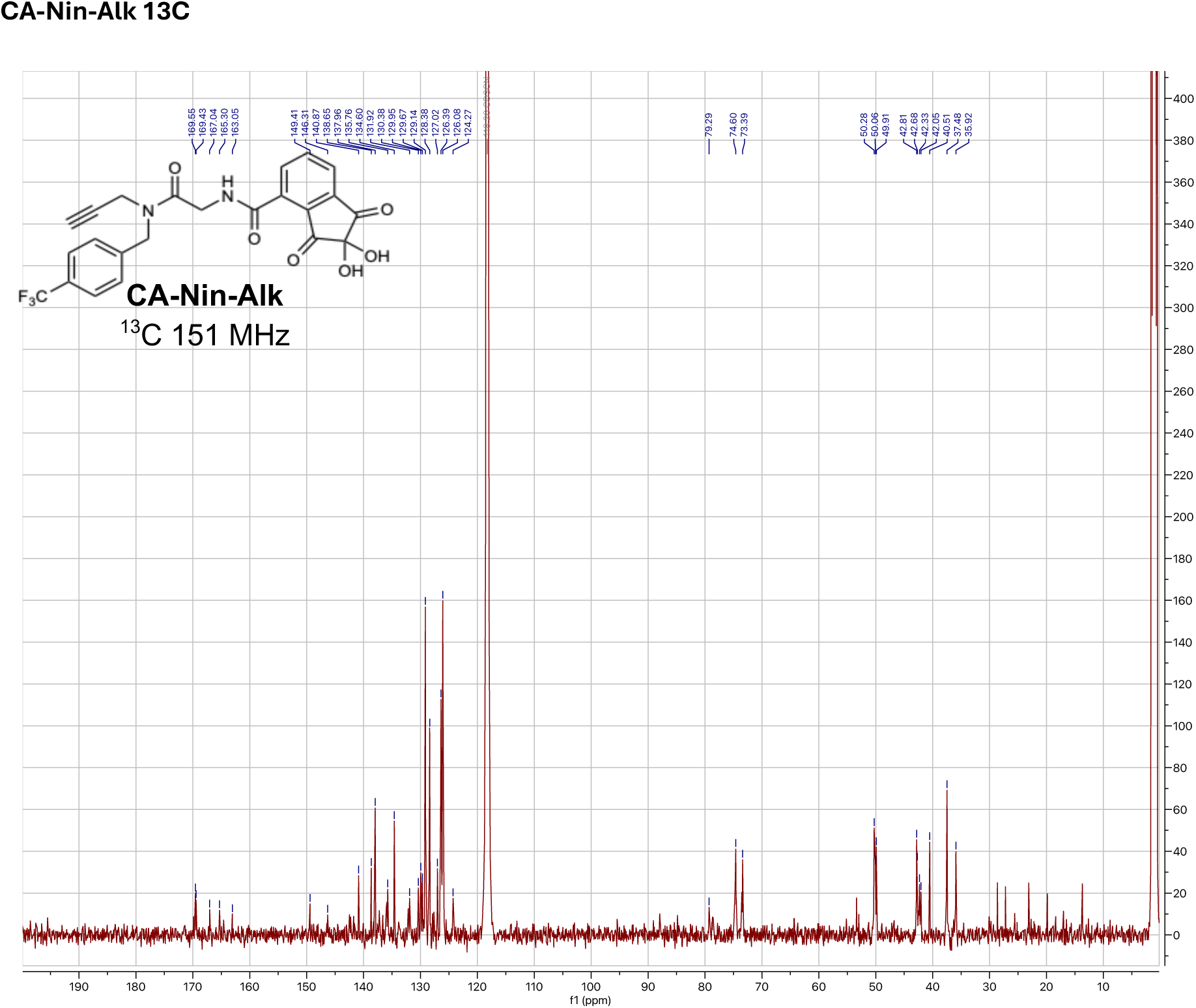

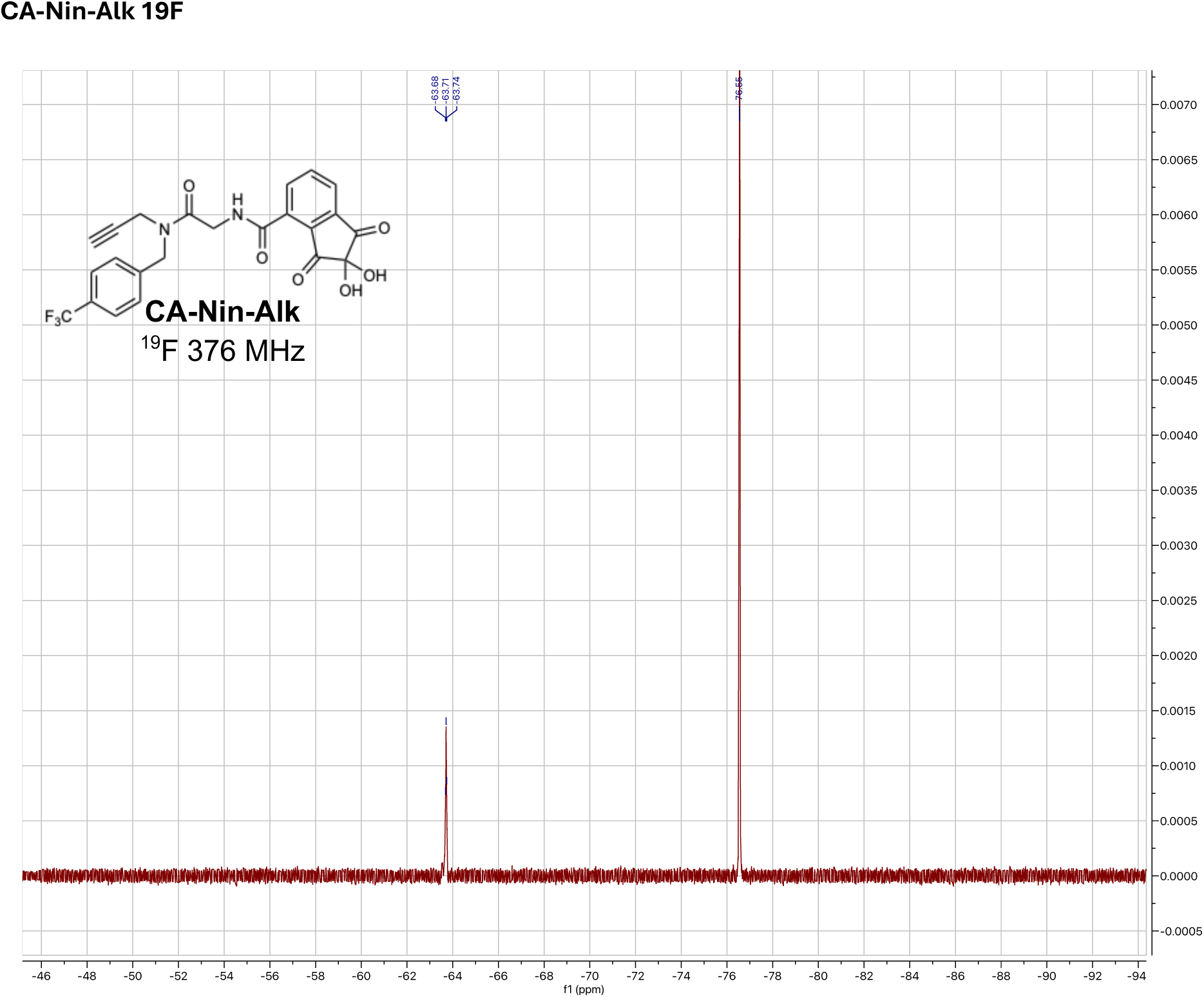

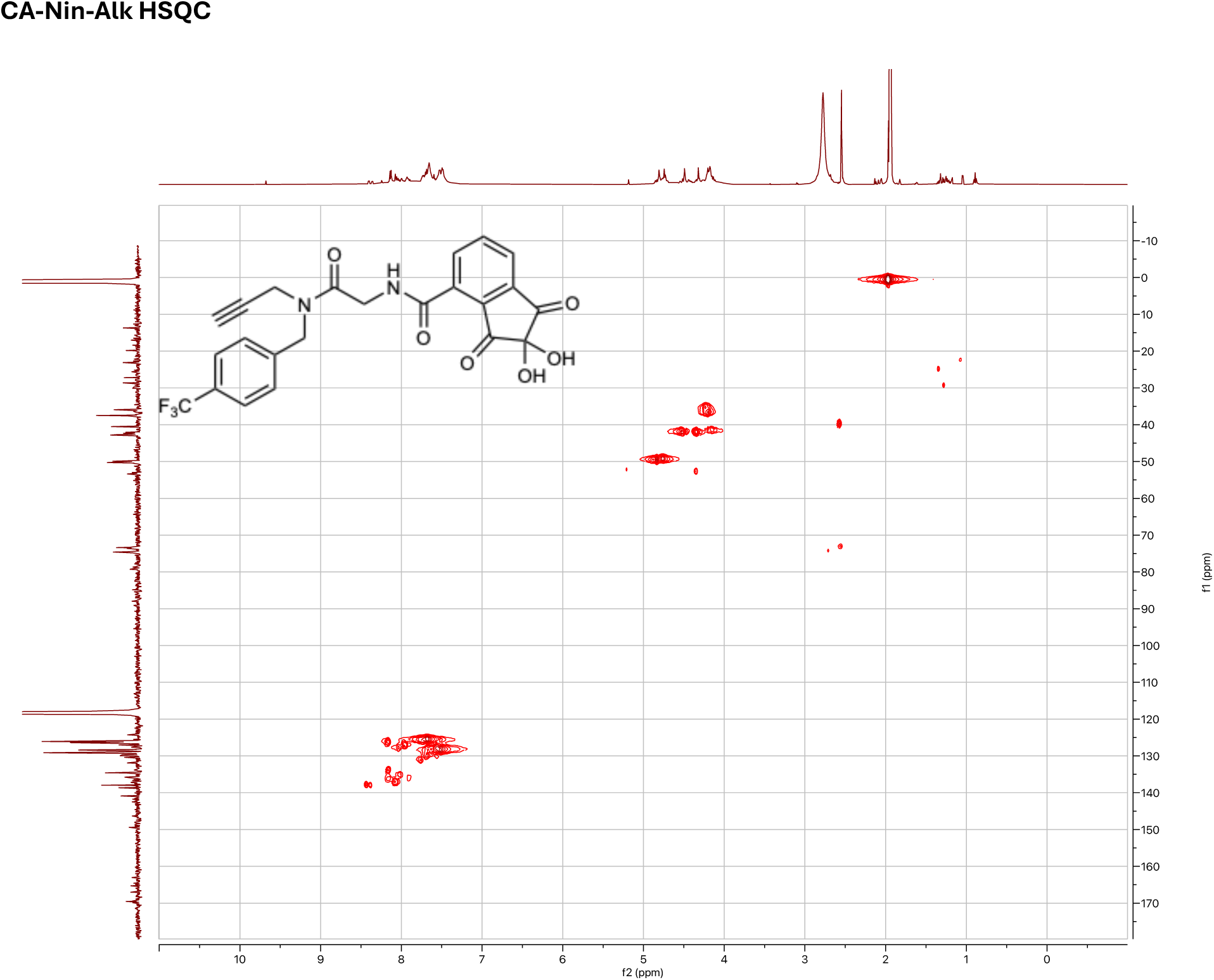

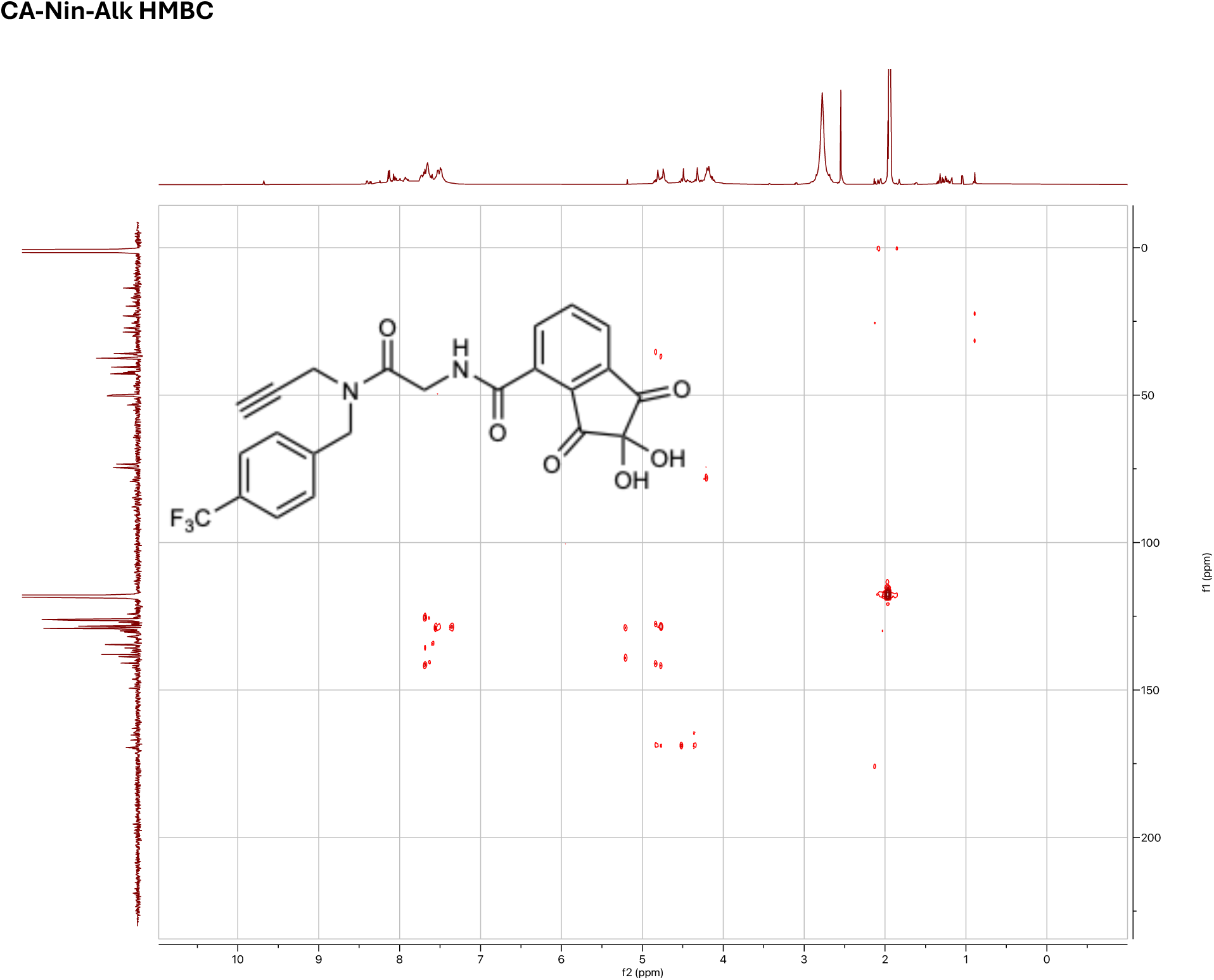

